# Influence of Energy Deficiency on the Molecular Processes of *Substantia Nigra Pars Compacta* Cell for Understanding Parkinsonian Neurodegeneration: A Comprehensive Biophysical Computational Model

**DOI:** 10.1101/2020.02.18.950337

**Authors:** Vignayanandam R. Muddapu, V. Srinivasa Chakravarthy

## Abstract

Parkinson’s disease (PD) is the second most prominent neurodegenerative disease around the world. Although it is known that PD is caused by the loss of dopaminergic cells in substantia nigra pars compacta (SNc), the decisive cause of this inexorable cell loss is not clearly elucidated. We hypothesize that “Energy deficiency at a sub-cellular/cellular/systems level can be a common underlying cause for SNc cell loss in PD.” Here, we propose a comprehensive computational model of SNc cell which helps us to understand the pathophysiology of neurodegeneration at subcellular level in PD. The proposed model incorporates a rich vein of molecular dynamics related to SNc neurons such as ion channels, active pumps, ion exchangers, dopamine turnover processes, energy metabolism pathways, calcium buffering mechanisms, alpha-synuclein aggregation, Lewy body formation, reactive oxygen species (ROS) production, levodopa uptake, and apoptotic pathways. The proposed model was developed and calibrated based on experimental data. The influx of glucose and oxygen into the model was controlled, and the consequential ATP variations were observed. Apart from this, the dynamics of other molecular players (alpha-synuclein, ROS, calcium, and dopamine) known to play an important role in PD pathogenesis are also studied. The aim of the study was to see how deficits in supply of energy substrates (glucose and oxygen) lead to a deficit in ATP, and furthermore, deficits in ATP are the common factor underlying the pathological molecular-level changes including alpha-synuclein aggregation, ROS formation, calcium elevation, and dopamine dysfunction. The model suggests that hypoglycemia plays a more crucial role in leading to ATP deficits than hypoxia. We believe that the proposed model provides an integrated modelling framework to understand the neurodegenerative processes underlying PD.

## INTRODUCTION

More than 200 years after it was first described by Dr. James Parkinson as ‘shaking palsy’ still we searching for a cure for Parkinson’s disease (PD), a neurodegenerative disorder characterized by the loss of dopaminergic cells in Substantia Nigra pars compacta (SNc) (McDonald et al., 2018). It is quite remarkable that loss of cells in a small nucleus like SNc can have wide-ranging devastating effects in all the four major domains of brain function – sensory-motor, cognitive, affective and autonomous (Goldman and Postuma, 2014). While existing treatments manage the symptoms of PD – sometimes with miraculous effect – a genuine cure demands an understanding of the root cause of SNc cell loss. Recently, a new approach towards PD etiology – that metabolic deficiencies at subcellular/cellular/network level can be a major cause of SNc cell loss in PD – was gaining attention (Bolam and Pissadaki, 2012; Muddapu et al., 2018, 2019; Muddapu and Chakravarthy, 2017; Pacelli et al., 2015; Wellstead and Cloutier, 2011).

In an earlier computational study, we have shown that metabolic deficiency at the systems/network level can lead to neurodegeneration of SNc neurons due to excitotoxicity caused by an overexcited Subthalamic Nucleus (STN) (Muddapu et al., 2018, 2019; Muddapu and Chakravarthy, 2017). As a further step in understanding the PD pathophysiology, in the present study, we proposed to model the effects of metabolic deficiencies in SNc at subcellular level. To this end, we need a comprehensive, holistic model of the SNc neuron with all the essential subcellular or molecular processes involved in PD pathogenesis. The model should include the standard molecular infrastructure like ion channels, active pumps, ion exchangers, dopamine turnover processes, energy metabolism pathways, and calcium buffering mechanisms and be able to simulate a rich vein of PD-related molecular processes such as, alpha-synuclein aggregation, Lewy body formation, reactive oxygen species (ROS) production, levodopa uptake, and apoptotic pathways. Several researchers had tried to model parts of the extensive chemical network involved in subcellular PD pathogenesis (Cloutier and Wellstead, 2012; Cullen and Wong-Lin, 2015; Francis et al., 2013; Reed et al., 2012; Tello-Bravo, 2012). From their modelling efforts it was evident that developing such a comprehensive model of SNc neuron would be a significant leap in understanding the subcellular mechanisms underlying neurodegeneration in PD. A comprehensive literature survey on modelling efforts related to PD pathogenesis was recently published (Bakshi et al., 2019; Lloret-Villas et al., 2017).

The proposed computational study aims to see how deficits in supply of energy substrates (glucose and oxygen) lead to a deficit in ATP, and furthermore, deficits in ATP are the common factor underlying the pathological changes in alpha-synuclein, ROS, calcium, and dopamine. Here, we propose a comprehensive computational model of SNc cell, which helps us in understand the pathophysiology of neurodegeneration in PD. The model is expected to help resolve several outstanding questions of PD pathology, e.g., the recurrent confusion of cause and effect – is alpha-synuclein aggregation a cause or an effect of PD? If the hypothesis that the model set out to investigate ultimately proves to be true, it will be demonstrated that energy deficiency underlies all the molecular level manifestations of PD. Such a demonstration, naturally, requires extensive and directed experimentation and the present model could perhaps serve as a blueprint for rolling out such an experimental program.

The model is developed as per the following stages. Firstly, each of the cellular processes in the model was calibrated by experimental data. Secondly, model responses to electrical and chemical stimulations were carried out to observe their effects on different vital molecular players in the SNc neuron. Finally, hypoglycemia and hypoxia conditions were simulated in the model to understand their adaptability to the energy crisis and to identify the different regimes, normal and pathological, in which the model operates.

## METHODS

The proposed comprehensive single-cell model of SNc dopaminergic neuron consists of ion-channel dynamics (Francis et al., 2013), calcium buffering mechanisms (Francis et al., 2013; Marhl et al., 2000), energy metabolism pathways (Cloutier and Wellstead, 2010, 2012), dopamine turnover processes (Tello-Bravo, 2012), LDOPA-uptake mechanisms (Reed et al., 2012), apoptotic pathways (Hong et al., 2012). and molecular pathways involved in PD pathology (Cloutier and Wellstead, 2012) (Figure 1).

**Figure 1:**
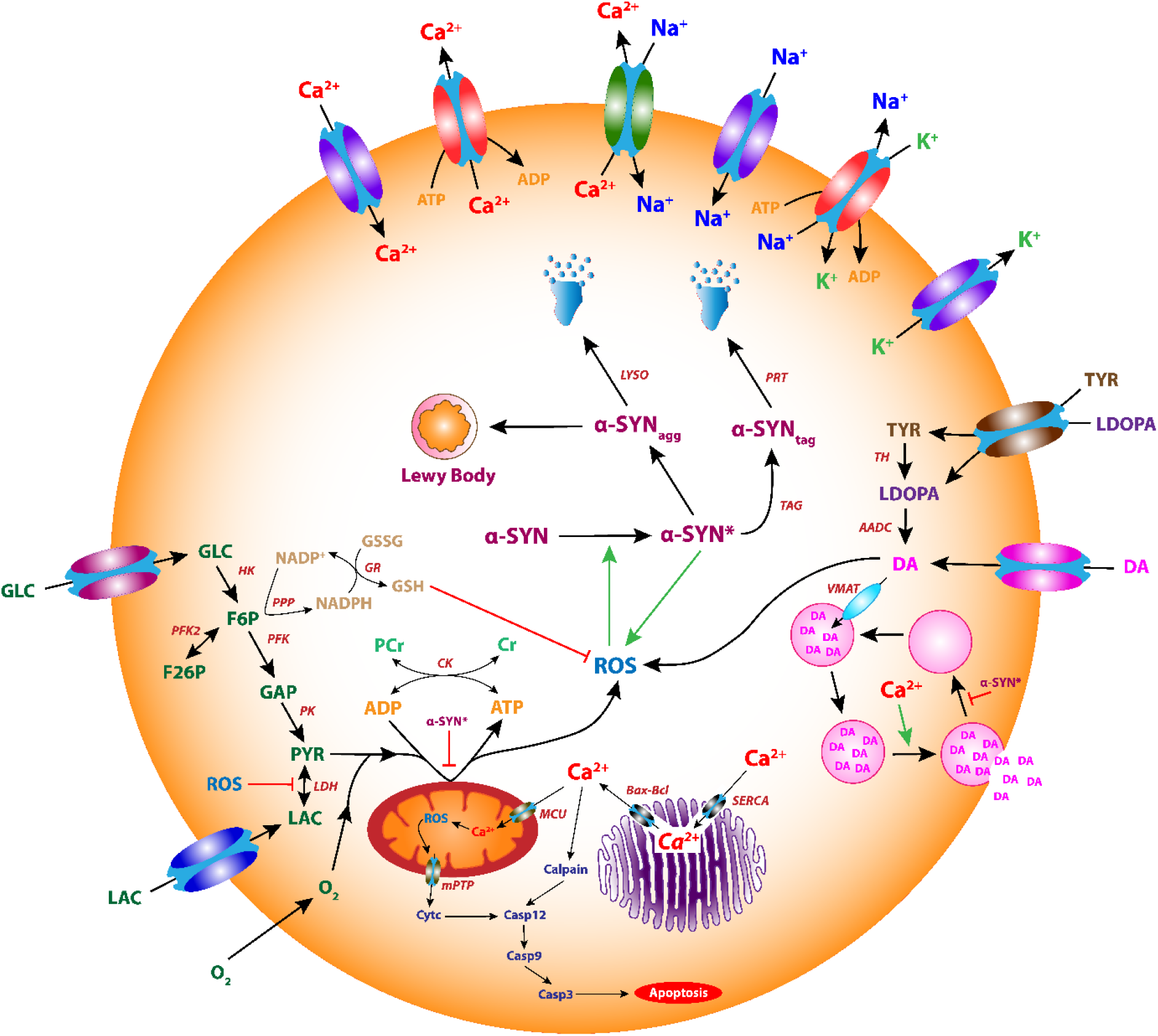
The proposed comprehensive model of the SNc neuron.

### Ion Channels

Modelling the behavior of a single neuron often requires detailed dynamics for a particular neuron type, since distinct electrophysiological and morphological features characterize each type of neuron. Dopaminergic neurons in substantia nigra exhibit two distinct firing patterns: low-frequency irregular tonic or background firing (1–5 *Hz*) and high-frequency regular phasic or burst firing (~ 20 *Hz*) (Grace and Bunney, 1984b, 1984a). Dopaminergic neurons are autonomously active and produce a constant background firing pattern on which bursts may be superimposed.

We have adapted the single-compartmental biophysical model of SNc (Francis et al., 2013) where ion-channel dynamics is dependent on ATP levels. Other previously published dopaminergic neuronal models are specified in *Supplementary material-1*. The ionic currents which were considered in the soma (Figure 2), voltage-dependent sodium currents (*I_Na_*), voltage-dependent potassium currents (*I_K_*), voltage-dependent L-type calcium current (*I_CaL_*), calcium-dependent potassium current (*I*_*K*(*Ca*)_), leak current (*I_L_*), sodium-potassium ATPase (*I_NaK_*), calcium ATPase (*I_pmca_*) and sodium-calcium exchanger (*I_NaCax_*).

**Figure 2:**
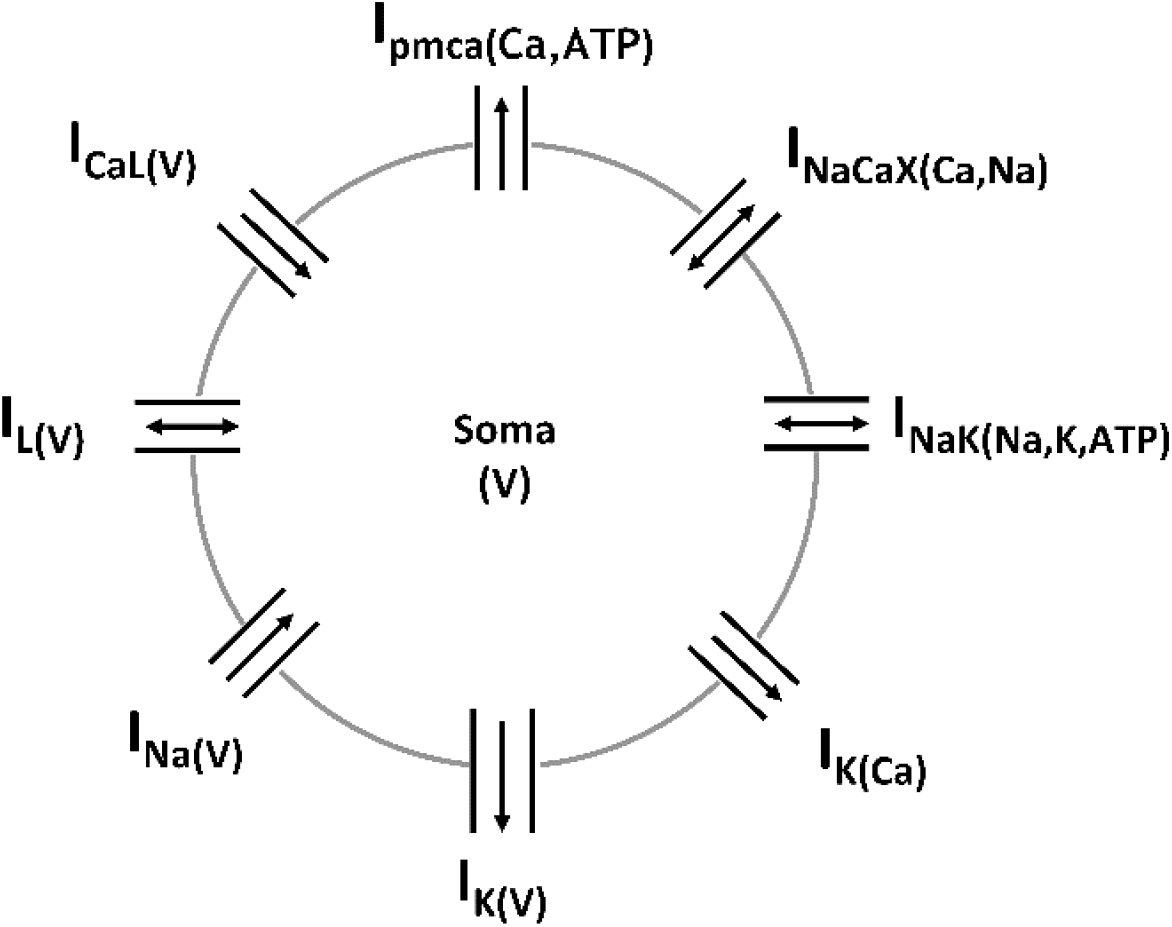
Schematic of the single-compartment DA neuron model demonstrating the various ion currents in the model. See text for description of the figure.

The membrane potential equation for the SNc soma (*V*) is given by,

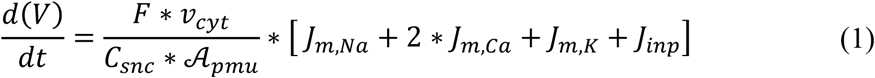

where, *F* is the Faraday’s constant, *C_snc_* is the SNc membrane capacitance, *v_cyt_* is the cytosolic volume, 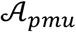 is the cytosolic area, *J_m,Na_* is the sodium membrane ion flux, *J_m,Ca_* is the calcium membrane ion flux, *J_m,K_* is the potassium membrane ion flux, *J_inp_* is the overall input current flux.

### Plasma Membrane Ion Channels

The intracellular calcium concentration dynamics ([*Ca_i_*]) is given by,

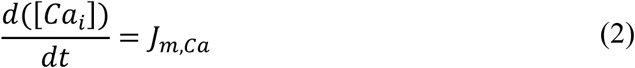

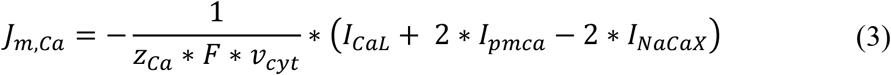

where, 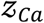 is the valence of calcium ion, *I_CaL_* is the L-type calcium channel current, *I_pmca_* is the ATP-dependent calcium pump current, *I_NaCax_* is the sodium-potassium exchanger current, *F* is the Faraday’s constant, *v_cyt_* is the cytosolic volume.

The voltage-dependent L-type calcium channel current (*I_CaL_*) is given by,

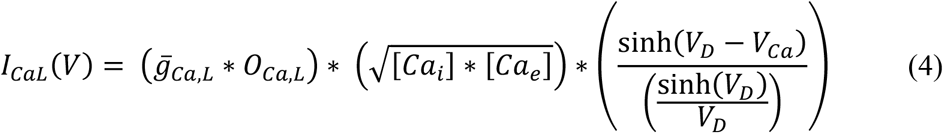

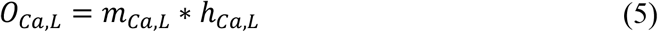

where, 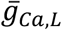 is the maximal conductance for calcium channel, *O_Ca,L_* is the gating variable of calcium channel, *m_Ca,L_* is the activation gate of the L-type calcium channel, *h_Ca,L_* is the inactivation gate of L-type calcium channel, [*Ca_i_*] is the intracellular calcium concentration, [*Ca_e_*] is the extracellular calcium concentration, *V_Ca_* is the reversal potential for calcium ion, *V_D_* is the voltage defined thermodynamic entity.

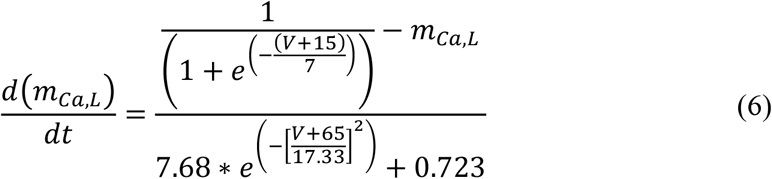

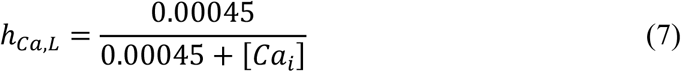

The intracellular sodium concentration ([*Na_i_*]) dynamics is given by,

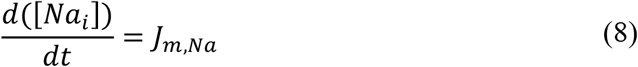

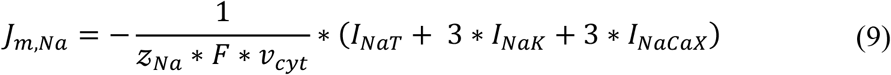

where, 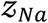 is the valence of sodium ion, *I_NaT_* is the total sodium channel current, *I_NaK_* is the ATP-dependent sodium-potassium pump current, *I_NaCax_* is the sodium-potassium exchanger current, *F* is the Faraday’s constant, *v_cyt_* is the cytosolic volume.

The total sodium channel current is given by,

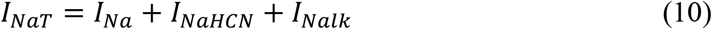

where, *I_Na_* is the voltage-dependent sodium channel current, *I_NaHCN_* is the hyperpolarization-activated cyclic nucleotide-gated sodium channel current, and *I_Nalk_* is the leaky sodium channel current.

The voltage-dependent sodium channel current (*I_Na_*) is given by,

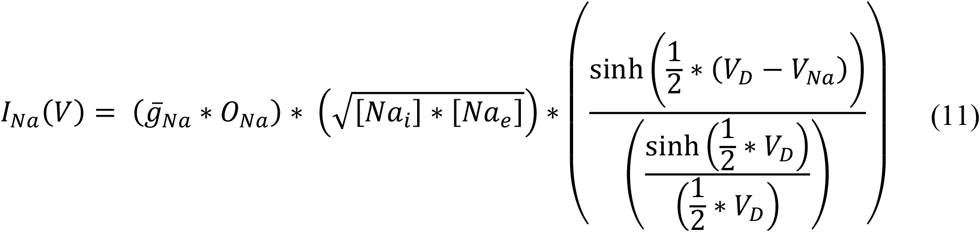

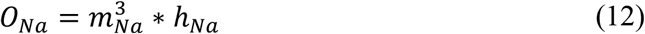

where, 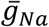 is the maximal conductance for sodium channel, *O_Na_* is the gating variable of sodium channel, *m_Na_* is the activation gate of the sodium channel, *h_Na_* is the inactivation gate of the sodium channel, [*Na_i_*] is the intracellular sodium concentration, [*Na_e_*] is the extracellular sodium concentration, *V_Na_* is the reversal potential for sodium ion, *V_D_* is a voltage-defined thermodynamic entity.

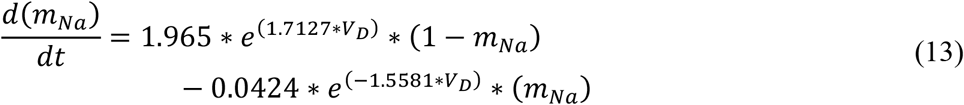

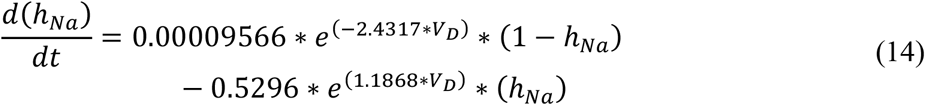

The hyperpolarization-activated cyclic nucleotide (HCN) gated sodium channel current (*I_NaHCN_*) is given by,

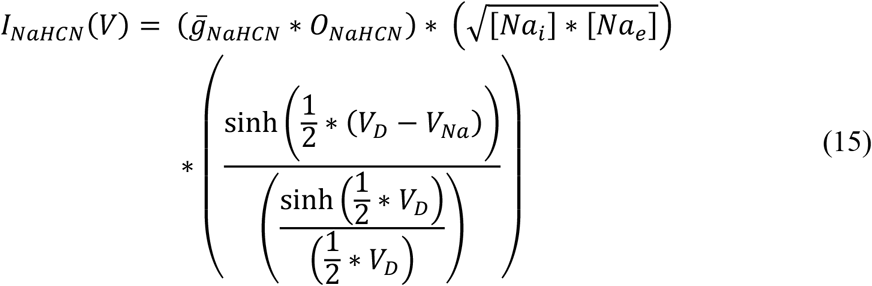

where, 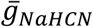 is the maximal conductance for sodium HCN channel, *O_NaHCN_* is the gating variable of sodium HCN channel, [*Na_i_*] is the intracellular sodium concentration, [*Na_e_*] is the extracellular sodium concentration, *V_Na_* is the reversal potential for sodium ion, *V_D_* is the voltage defined thermodynamic entity, [*cAMP*] is the cyclic adenosine monophosphate concentration.

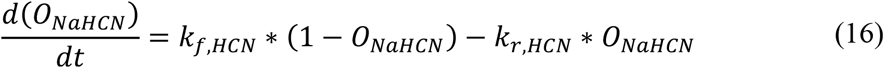

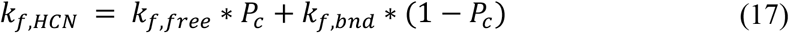

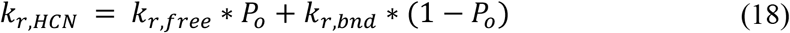

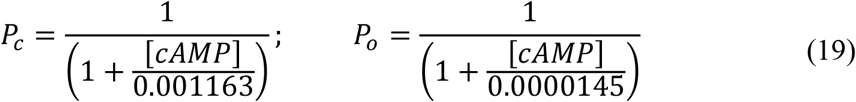

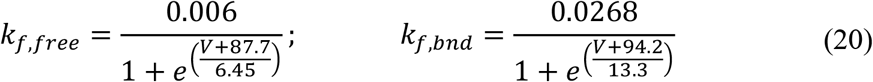

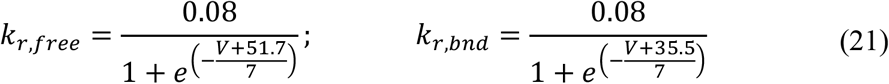

The leaky sodium channel current (*I_Nalk_*) is given by,

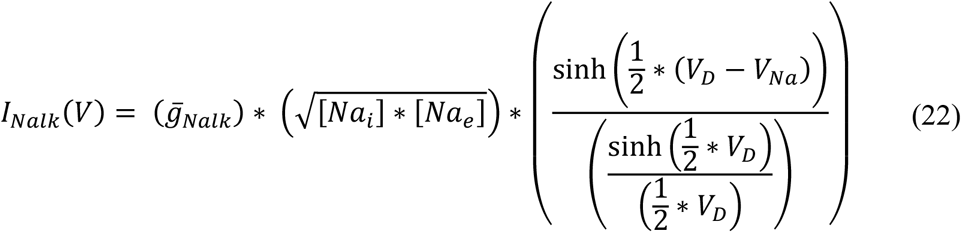

where, 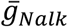 is the maximal conductance for leaky sodium channel, [*Na_i_*] is the intracellular sodium concentration, [*Na_e_*] is the extracellular sodium concentration, *V_Na_* is the reversal potential for sodium ion, *V_D_* is the voltage defined thermodynamic entity.

The intracellular potassium concentration dynamics ([*K_i_*]) is given by,

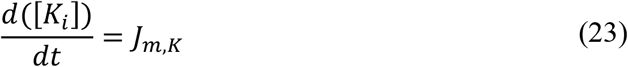

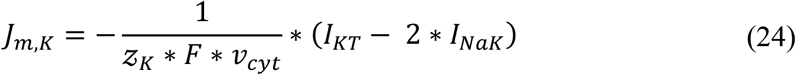

where, 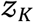 is the valence of potassium ion, *I_KT_* is the total potassium channel current, *I_NaK_* is the ATP-dependent sodium-potassium pump current, *F* is the Faraday’s constant, *v_cyt_* is the cytosolic volume.

The total potassium channel current is given by,

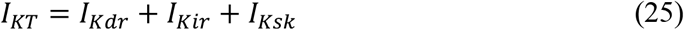

where, *I_Kdr_* is the voltage-dependent (delayed rectifying, DR) potassium channel current, *I_Kir_* is the voltage-dependent (inward rectifying, IR) potassium channel current, *I_Ksk_* is the calcium-dependent (small conductance, SK) potassium channel current.

The voltage-dependent (delayed rectifying) potassium channel current (*I_Kdr_*) is given by,

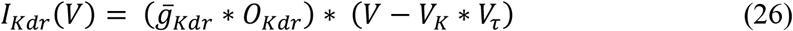

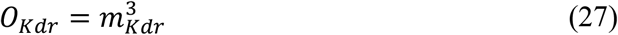

where, 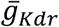 is the maximal conductance for delayed rectifying potassium channel, *O_Kdr_* is the gating variable of voltage-dependent (delayed rectifying) potassium channel, *V_K_* is the reversal potential for potassium ion, *V_τ_* is the temperature defined thermodynamic entity.

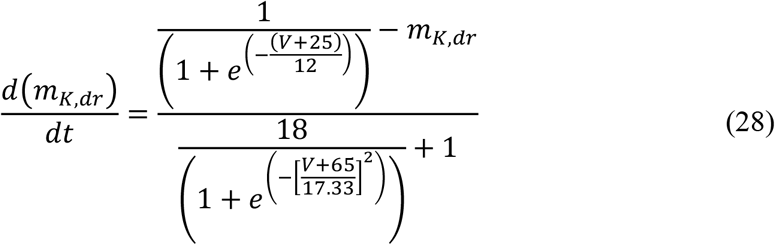

The voltage-dependent (inward rectifying) potassium channel current (*I_Kir_*) is given by,

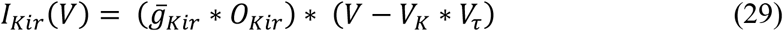

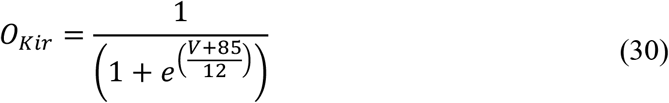

where, 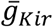 is the maximal conductance for inward rectifying potassium channel, *O_Kir_* is the gating variable of voltage-dependent (inward rectifying) potassium channel, *V_K_* is the reversal potential for potassium ion, *V_τ_* is the temperature defined thermodynamic entity.

The calcium-dependent (small conductance) potassium channel current (*I_Ksk_*) is given by,

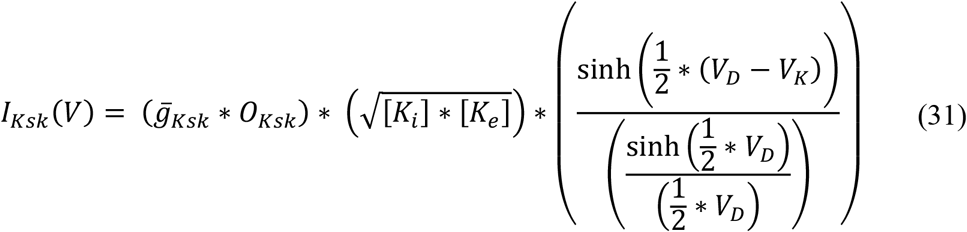

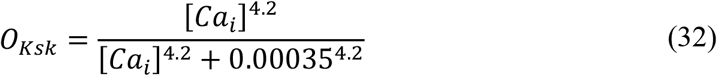

where, 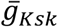 is the maximal conductance for small conductance potassium channel, *O_Ksk_* is the gating variable of calcium-dependent (small conductance) potassium channel, [*K_i_*] is the intracellular potassium concentration, [*K_e_*] is the extracellular potassium concentration, [*Ca_i_*] is the intracellular calcium concentration, *V_K_* is the reversal potential for potassium ion, *V_D_* is the voltage defined thermodynamic entity.

The overall synaptic input current flux (*J_syn_*) to SNc neuron is given by,

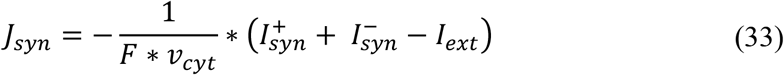

where, 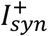 is the excitatory synaptic current, 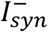 is the inhibitory synaptic current, *I_ext_* is the external current applied, *F* is the Faraday’s constant, *v_cyt_* is the cytosolic volume. The different types of synaptic receptors were modeled as similar to (Destexhe et al., 1998) and given in *Supplementary material-4*.

### Plasma Membrane ATPases

The plasma membrane sodium-potassium ATPase (*I_NaK_*) is given by,

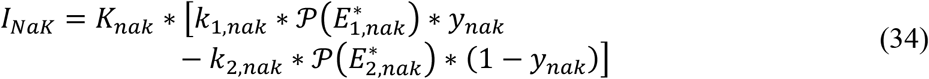

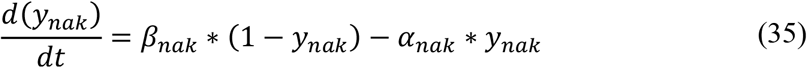

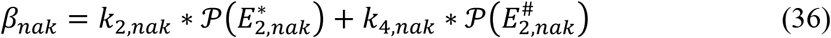

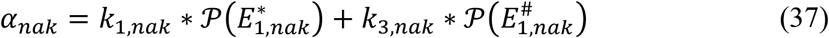

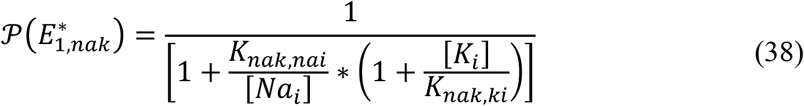

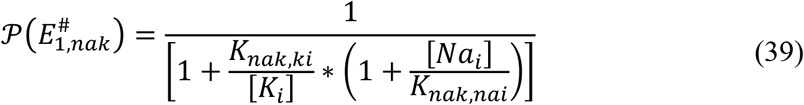

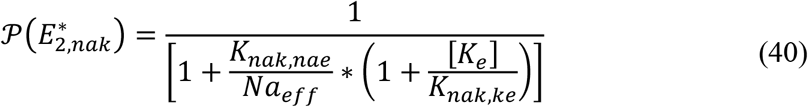

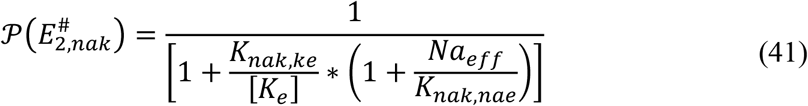

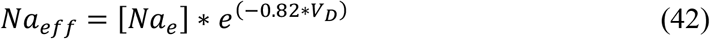

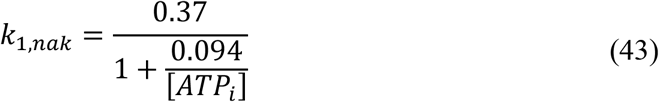

where, *K_nak_* is the maximal conductance for sodium-potassium ATPase, [*Na_i_*] is the intracellular concentration of sodium ion, [*Na_e_*] is the extracellular concentration of sodium ion, [*K_i_*] is the intracellular concentration of potassium ion, [*K_e_*] is the extracellular concentration of potassium ion, (*k*_1,*nak*_, *k*_2,*nak*_, *k*_3,*nak*_, *k*_4,*nak*_) are the reaction rates, (*K_nak,nae_, K_nak,nai_, K_nak,ke_, K_nak,ki_*) are the dissociation constants, [*ATP_i_*] is the intracellular concentration of adenosine triphosphate (ATP), *V_D_* is the voltage defined thermodynamic entity.

The plasma membrane calcium ATPase (*I_pmca_*) is given by,

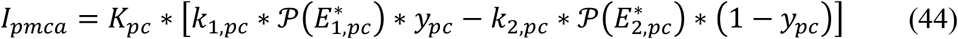

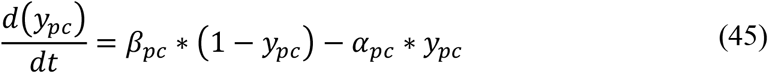

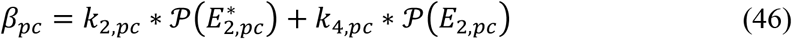

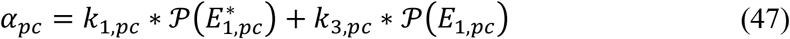

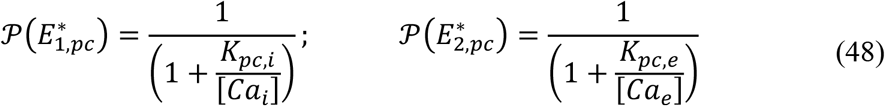

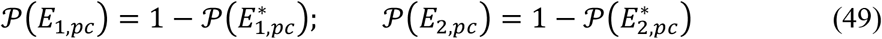

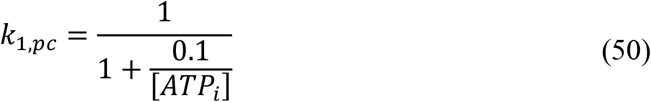

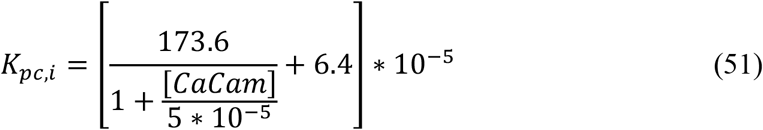

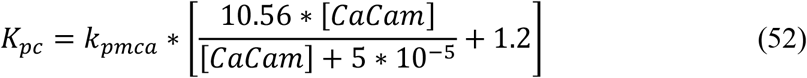

where, (*k*_1,*pc*_, *k*_2,*pc*_, *k*_3,*pc*_, *k*_4,*pc*_) are the reaction rates, *k_pmca_* is the maximal conductance for calcium ATPase, (*K_pc,e_*, *K_pc,i_*) are the dissociation constants, [*ATP_i_*] is the intracellular concentration of ATP, [*Ca_i_*] is the intracellular calcium concentration, [*CaCam*] is the intracellular calcium-bound calmodulin concentration.

### Plasma Membrane Exchangers

The plasma membrane sodium-calcium exchanger (*I_NaCax_*) is given by,

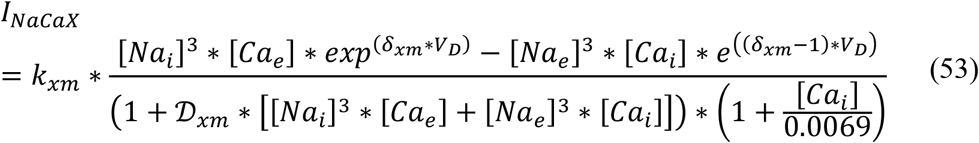

where, *k_xm_* is the maximal conductance for sodium-calcium exchanger, [*Na_e_*] is the extracellular sodium concentration, [*Na_i_*] is the intracellular sodium concentration, [*Ca_e_*] is the extracellular calcium concentration, [*Ca_i_*] is the intracellular calcium concentration, *δ_xm_* is the energy barrier parameter, 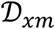 is the denominator factor, *V_D_* is the voltage defined thermodynamic entity.

### Calcium Buffering Mechanisms

The intracellular calcium plays an essential role in the normal functioning of the cell. In order to maintain calcium homeostasis, the intracellular calcium levels are tightly regulated by calcium buffering mechanisms such as calcium-binding proteins, endoplasmic reticulum (ER), and mitochondria (MT) (Figure 3) (Alzheimer, 2003).

**Figure 3:**
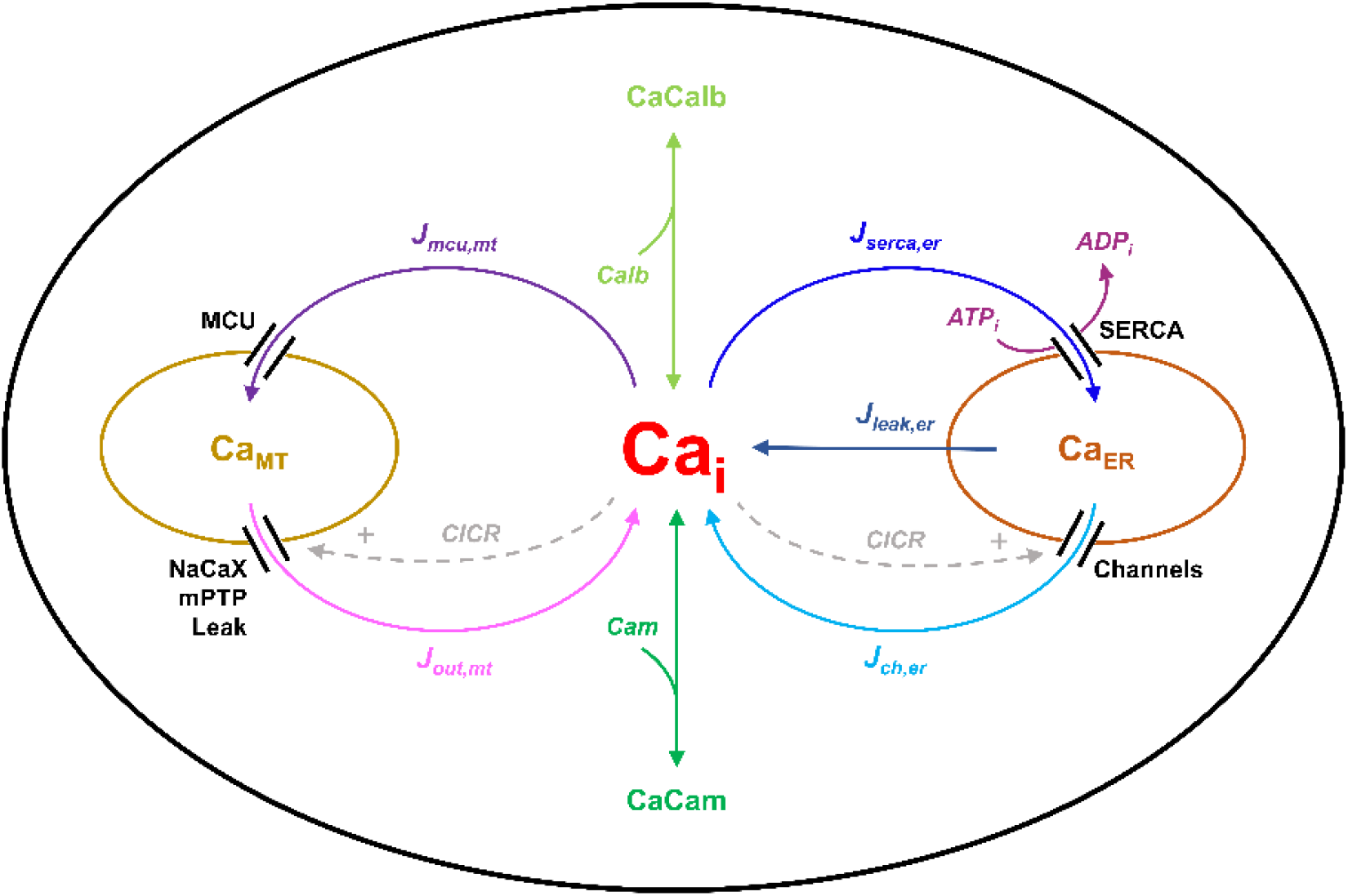
Schematic of calcium buffering mechanisms in the SNc cell model. See text for description of the figure.

The intracellular calcium concentration dynamics ([*Ca_i_*]) after including calcium buffering mechanisms (Francis et al., 2013; Marhl et al., 2000) (Figure 3) is given by,

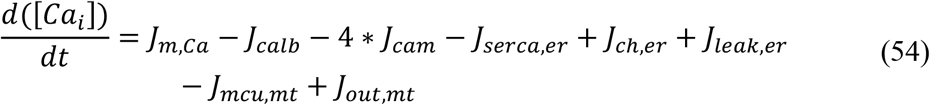

where, *J_m,Ca_* is the flux of calcium ion channels, *J_calb_* is the calcium buffering flux by calbindin, *J_cam_* is the calcium buffering flux by calmodulin, *J_serca,er_* is the calcium buffering flux by ER uptake of calcium through sarco/endoplasmic reticulum calcium-ATPase (SERCA), *J_ch,er_* is the calcium efflux from ER by calcium-induced calcium release (CICR) mechanism, *J_leak,er_* is the calcium leak flux from ER, *J_mcu,mt_* is the calcium buffering flux by MT uptake of calcium through mitochondrial calcium uniporters (MCUs), *J_out,mt_* is the calcium efflux from MT through sodium-calcium exchangers, mitochondrial permeability transition pores (mPTPs) and non-specific leak flux.

The calcium buffering flux by calbindin (*J_calb_*) is given by,

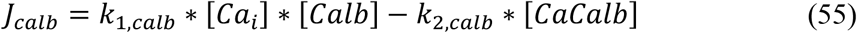

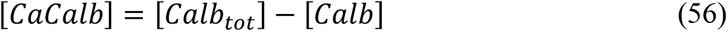

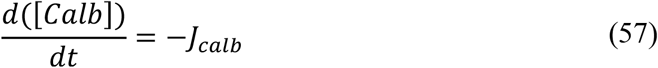

where, (*k*_1,*calb*_, *k*_2,*calb*_) are the calbindin reaction rates, [*Ca_i_*] is the intracellular calcium concentration, [*Calb*] is the calbindin concentration, [*CaCalb*] is the calcium-bound calbindin concentration, [*Calb_tot_*] is the total cytosolic calbindin concentration.

The calcium buffering flux by calmodulin (*J_cam_*) is given by,

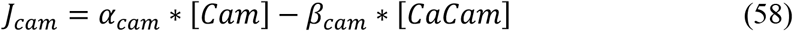

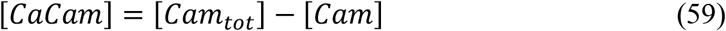

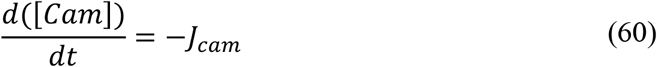

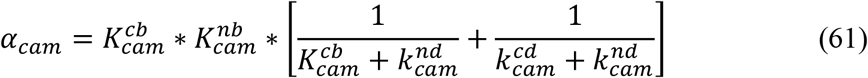

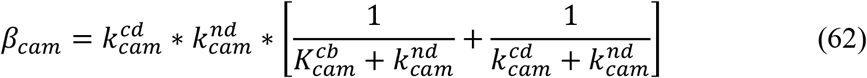

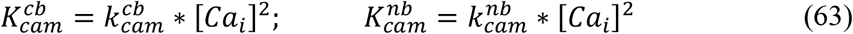

where, 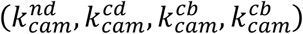 are the calmodulin reaction rates, [*Ca_i_*] is the intracellular calcium concentration, [*Cam*] is the calmodulin concentration, [*CaCam*] is the calcium-bound calmodulin concentration, [*Cam_tot_*] is the total cytosolic calmodulin concentration.

The calcium buffering flux by ER uptake of calcium through SERCA (*J_serca,er_*) is given by,

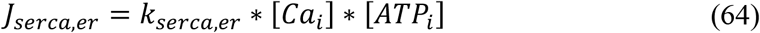

where, *k_serca,er_* is the maximal rate constant of SERCA, [*Ca_i_*] is the intracellular calcium concentration, [*ATP_i_*] is the intracellular ATP concentration.

The calcium efflux from ER by CICR (*J_cicr,er_*) is given by,

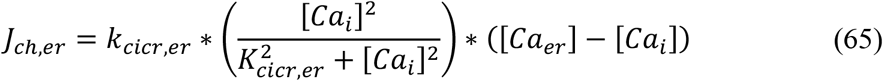

where, *k_ch,er_* is the maximal permeability of calcium channels in the ER membrane, *K_ch,er_* is the half-saturation for calcium, [*Ca_i_*] is the intracellular calcium concentration, [*Ca_er_*] is the ER calcium concentration.

The calcium leak flux from ER (*J_leak,er_*) is given by,

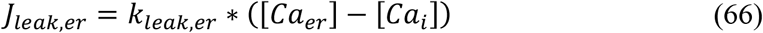

where, *k_leak,er_* is the maximal rate constant for calcium leak flux through the ER membrane, [*Ca_i_*] is the intracellular calcium concentration, [*Ca_er_*] is the ER calcium concentration.

The ER calcium concentration ([*Ca_er_*]) dynamics is given by,

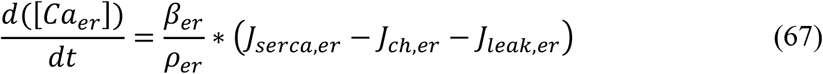

where, *β_er_* is the ratio of free calcium to total calcium concentration in the ER, *ρ_er_* is the volume ratio between the ER and cytosol, *J_serca,er_* is the calcium buffering flux by ER uptake of calcium through SERCA, *J_ch,er_* is the calcium efflux from ER by CICR mechanism, *J_leak,er_* is the calcium leak flux from ER.

The calcium buffering flux by MT uptake of calcium through MCUs (*J_mcu,mt_*) is given by,

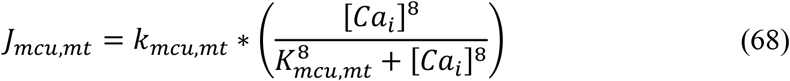

where, *k_mcu,mt_* is the maximal permeability of mitochondrial membrane calcium uniporters, *K_mcu,mt_* is the half-saturation for calcium, [*Ca_i_*] is the intracellular calcium concentration.

The calcium efflux from MT through sodium-calcium exchangers, mPTPs and non-specific leak flux (*J_out,mt_*) is given by,

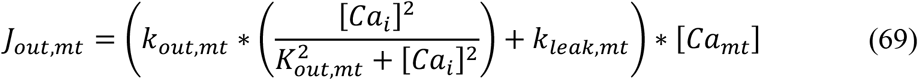

where, *k_out,mt_* is the maximal rate of calcium flux through sodium-calcium exchangers and mitochondrial permeability transition pores, *K_out,mt_* is the half-saturation for calcium, *k_leak,mt_* is the maximal rate constant for calcium leak flux through the MT membrane, [*Ca_i_*] is the intracellular calcium concentration, [*Ca_mt_*] is the MT calcium concentration.

The MT calcium concentration ([*Ca_mt_*]) dynamics is given by,

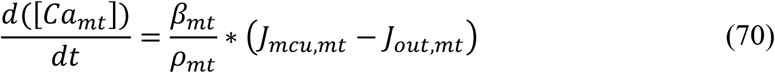

where, *β_mt_* is the ratio of free calcium to total calcium concentration in the ER, *ρ_mt_* is the volume ratio between the MT and cytosol, *J_mcu,mt_* is the calcium buffering flux by MT uptake of calcium through MCUs, *J_out,mt_* is the calcium efflux from MT through sodium-calcium exchangers, mPTPs and non-specific leak flux.

The total instantaneous concentration of calcium ([*Ca_tot_*]) in a SNc cell at a given time *t* is given by,

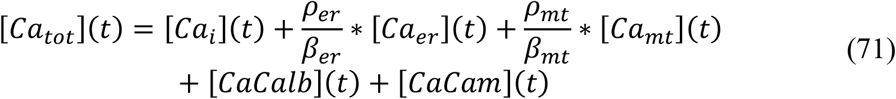

where, *β_er_* is the ratio of free calcium to total calcium concentration in the ER, *ρ_er_* is the volume ratio between the ER and cytosol, *β_mt_* is the ratio of free calcium to total calcium concentration in the ER, *ρ_mt_* is the volume ratio between the MT and cytosol, [*Ca_i_*](*t*), [*Ca_er_*](*t*), [*Ca_mt_*](*t*), [*CaCalb*](*t*), and [*CaCam*](*t*) are instantaneous concentration of intracellular (cytoplasmic) calcium, ER calcium, MT calcium, calcium-bound calbindin, and calcium-bound calmodulin, respectively.

### Energy Metabolism Pathways

The energy metabolism pathways which were included in the comprehensive model of SNc were adapted from (Cloutier and Wellstead, 2010) (Figure 4). Extracellular glucose (*GLC_e_*) is taken up into the neuron through glucose transporters and phosphorylated into fructose-6-phosphate (F6P) by hexokinase (HK) enzyme using adenosine triphosphate (ATP). F6P is broken down into glyceraldehyde-3-phosphate (GAP) by phosphofructokinase (PFK) enzyme using ATP. At steady state, F6P (fructose-2,6-bisphosphate (F26P)) is phosphorylated (dephosphorylated) to F26P (F6P) by dephosphorylating (phosphorylating) ATP (ADP) using phosphofructokinase-2 (PFK2) enzyme. GAP is dephosphorylated into pyruvate (PYR) by producing ATP using pyruvate kinase (PK). ATP is produced by MT through oxidative phosphorylation (OP) by utilizing PYR and oxygen (*O*_2_). Parallel to glycolysis, F6P is utilized to produce Nicotinamide adenine dinucleotide phosphate hydrogen (NADPH) through pentose phosphate pathway. Synthesized NADPH is used to produce glutathione (GSH) by glutathione reductase (GR) which scavenges reactive oxygen species (ROS). ATP is replenished by oxidative phosphorylation independent pathway where phosphocreatine is broken to produce ATP and creatine by creatine kinase (CK).

**Figure 4:**
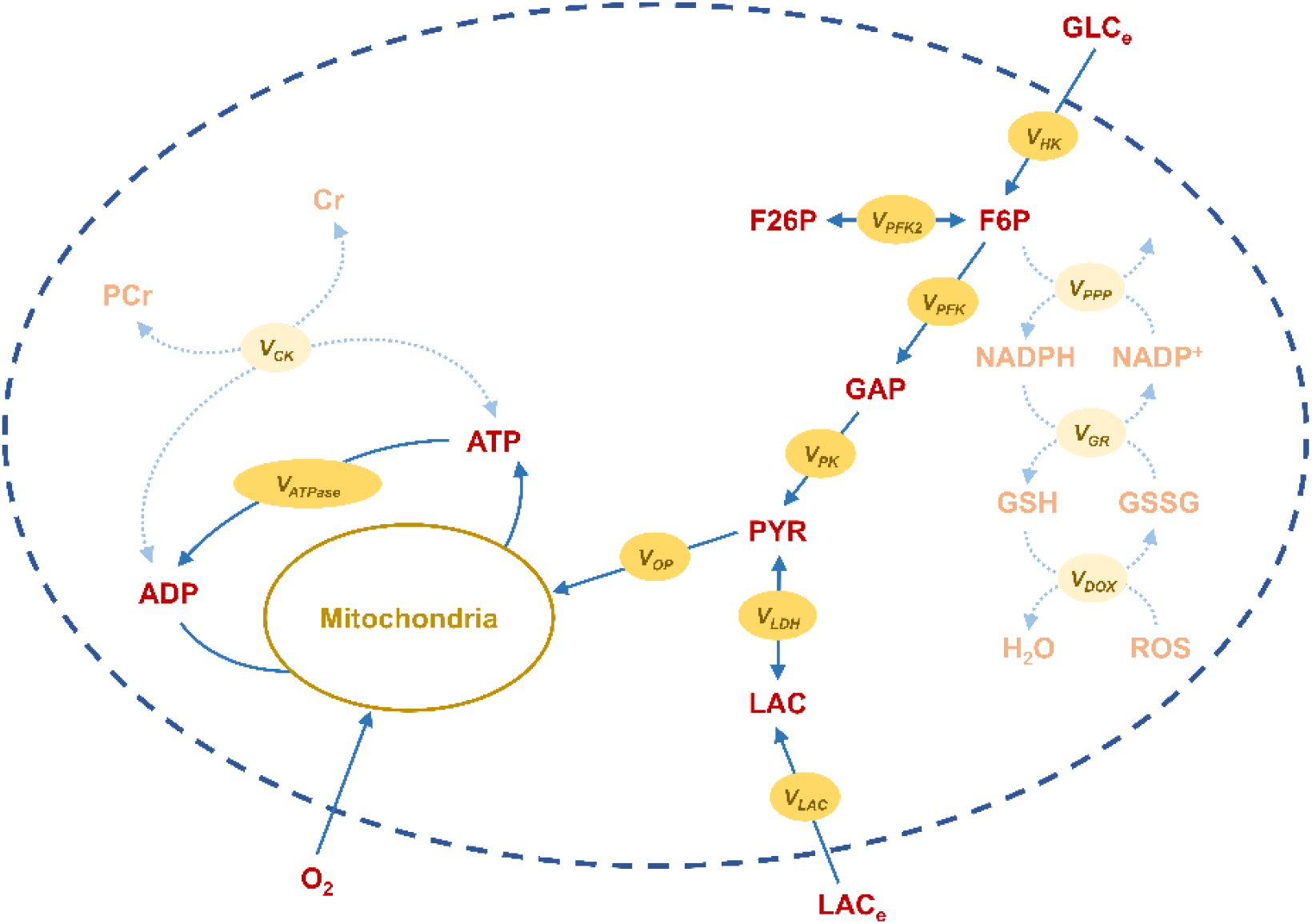
Schematic of energy mechanism pathways in the SNc cell model. See text for description of the figure.

The following equations give a concise view of all metabolite dynamics in the energy metabolism pathway:

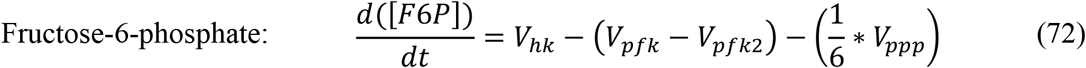

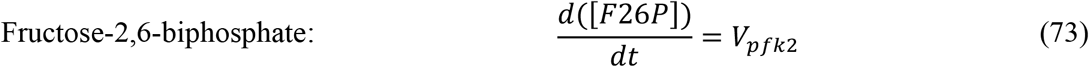

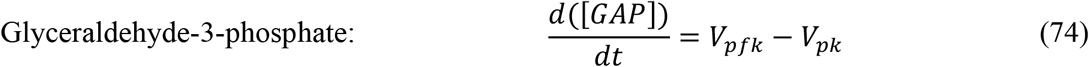

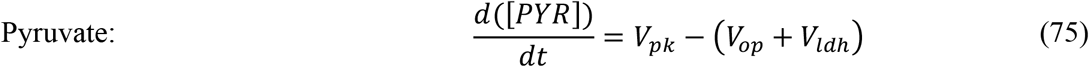

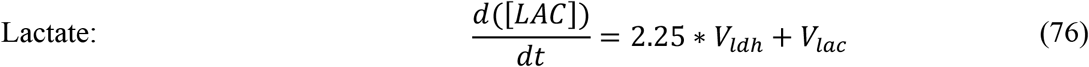

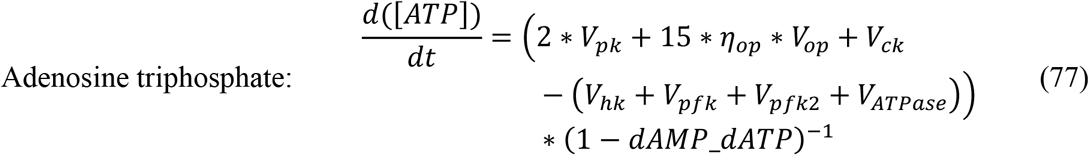

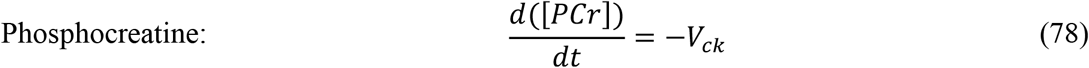

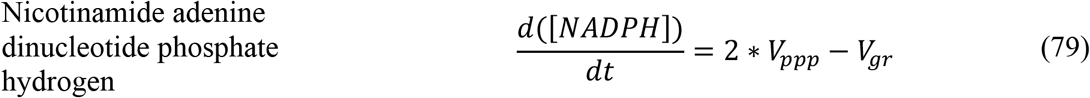

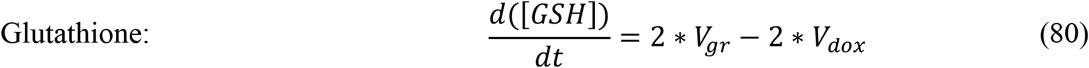

where, *V_hk_* is the irreversible flux of hexokinase enzyme where glucose was phosphorylated to F6P by using ATP, *V_pfk_* is the irreversible flux of phosphofructokinase enzyme where F6P was broken down to GAP using ATP, *V_pfk2_* is the reversible flux of phosphofructokinase-2 enzyme where F6P (F26P) was phosphorylated (dephosphorylated) to F26P (F6P) by dephosphorylating (phosphorylating) ATP (ADP), *V_ppp_* is the irreversible flux of the pentose phosphate pathway where NADP+ was reduced to NADPH, *V_pk_* is the irreversible flux of pyruvate kinase enzyme where GAP was dephosphorylated to PYR by phosphorylating adenosine diphosphate (ADP), *V_op_* is the irreversible flux of the oxidative phosphorylation pathway where PYR was utilized to produce ATP, *η_op_* is the electron transport chain efficiency, *V_ldh_* is the reversible flux of lactate dehydrogenase where LAC (PYR) was dehydrogenase (hydrogenase) to PYR (LAC), *V_lac_* is the reversible flux of monocarboxylate transporters where LAC from extracellular (intracellular) was transported into (out of) the cell, *V_ck_* is the reversible flux of creatine kinase where PCr (creatine (Cr)) was dephosphorylated (phosphorylated) to Cr (PCr) by phosphorylating (dephosphorylating) ADP (ATP), *V_gr_* is the irreversible flux of glutathione reductase where glutathione disulfide (GSSG) was reduced to GSH, *V_dox_* is the irreversible flux of anti-oxidative pathway where reactive oxygen species (ROS) was reduced to water, *V_ATPase_* is the irreversible flux of ATPases where ion equilibrium was maintained by utilizing ATP.

The flux of hexokinase (*V_hk_*) is given by,

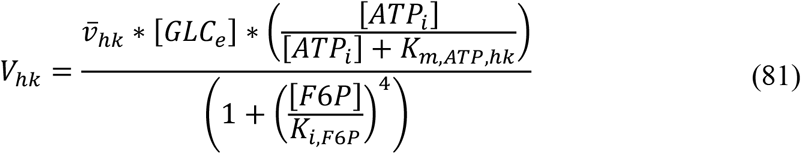

where, 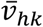 is the maximal hexokinase flux, [*ATP_i_*] is the intracellular ATP concentration, [*F6P*] is the F6P concentration, *K_m,ATP,hk_* is the affinity constant for ATP, *K_i,F6P_* is the inhibition constant for F6P, [*GLC_e_*] is the extracellular glucose concentration.

The flux of phosphofructokinase (*V_pfk_*) is given by,

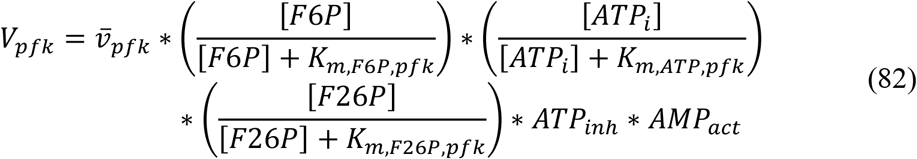

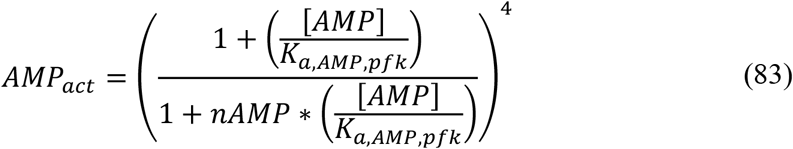

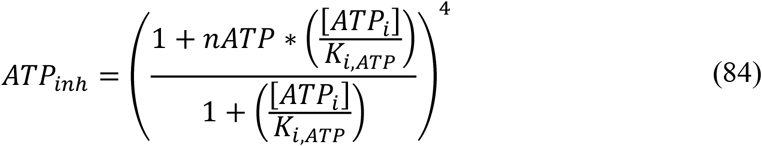

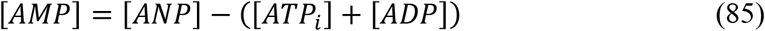

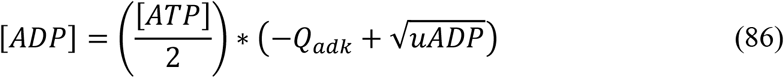

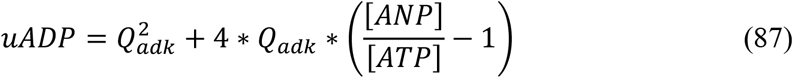

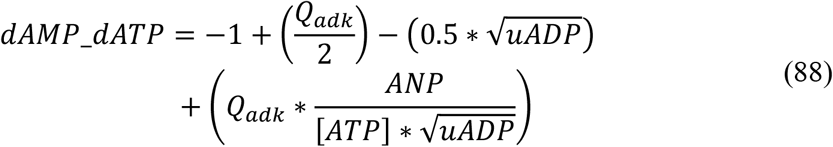

where, 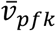 is the maximal phosphofructokinase flux, [*ATP_i_*] is the intracellular ATP concentration, [*F*6*F*] is the F6P concentration, [*F*26*P*] is the F26P concentration, *K*_*m*,*F*6*P,pfk*_ is the affinity constant for F6P, *K_m,ATP,pfk_* is the affinity constant for ATP, *K*_*m*,*F*26*P*,*pfk*_ is the affinity constant for F26P, [*AMF*] is the adenosine monophosphate (AMP) concentration, [*ADP*] is the adenosine diphosphate (ADP) concentration, [*ANP*] is the total energy shuttle’s (ANP) concentration, *K_a,AMP,pfk_* is the activation constant for AMP, *K_i,ATP_* is the inhibition constant for ATP, *nAMP* is the coefficient constant for AMP, *nATP* is the coefficient constant for ATP, *Q_adk_* is the coefficient constant for ADP.

The flux of phosphofructokinase-2 (*V*_*pfk*2_) is given by,

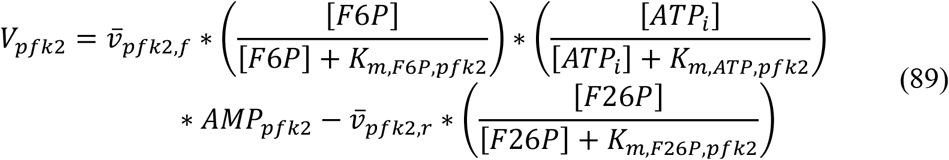

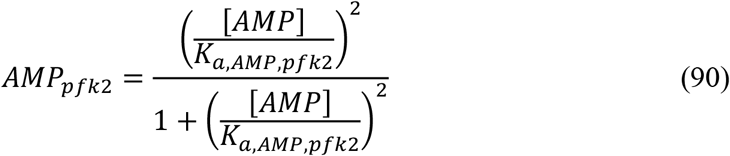

where, 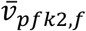 is the maximal phosphofructokinase-2 forward flux, 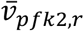 is the phosphofructokinase-2 maximal reverse flux, [*ATP_i_*] is the intracellular ATP concentration, [F6P] is the F6P concentration, [F26P] is the F26P concentration, [AMP] is the AMP concentration, *K_m,F6P,pfk2_* is the affinity constant for F6P, *K_m,ATP,pfk2_* is the affinity constant for ATP, *K_m,F26P,pfk2_* is the affinity constant for F26P, *K_a,AMP,pfk2_* is the activation constant for AMP.

The flux of pyruvate kinase (*V_pk_*) is given by,

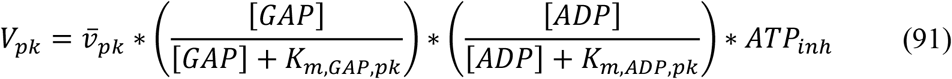

where, 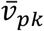 is the pyruvate kinase maximal flux, [*GAP*] is the GAP concentration, [*ADP*] is the ADP concentration, *K_m,GAP,pk_* is the affinity constant for GAP, *K_m,ADP,pk_* is the affinity constant for ADP, *ATP_inh_* is the ATP inhibition term.

The flux of the oxidative phosphorylation pathway (*V_op_*) is given by,

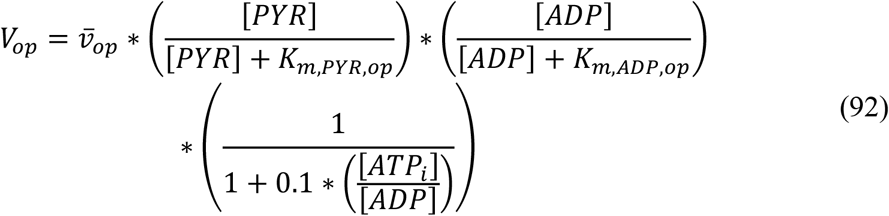

where, 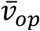 is the oxidative phosphorylation pathway maximal flux, [*PYR*] is the PYR concentration, [*ADP*] is the ADP concentration, [*ATP_i_*] is the ATP concentration, *K_m,PYR,op_* is the affinity constant for PYR, *K_m,ADP,op_* is the affinity constant for ADP.

In the absence of protein aggregation, the electron transport chain efficiency is given by,

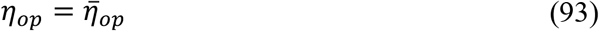

Moreover, in the presence of protein aggregation, the electron transport chain efficiency is given by,

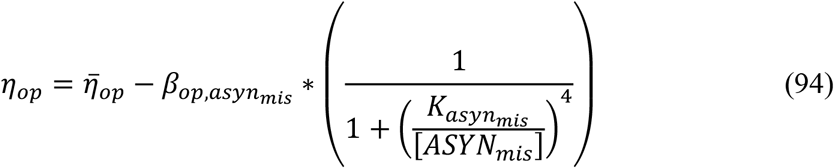

where, 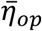 is the maximal electron transport chain efficiency, *β_op,asyn_mis__* is the maximum fractional decrease in the oxidative phosphorylation efficiency through misfolded alpha-synuclein (*ASYN_mis_*), [*ASYN_mis_*] is the misfolded alpha-synuclein concentration, *K_asyn_mis__* is the threshold concentration for mitochondrial damage by *ASYN_mis_*.

The flux of lactate dehydrogenase (*V_ldh_*) is given by,

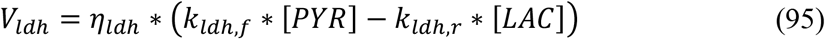

where, *η_ldh_* is the lactate fermentation efficiency, [*PYR*] is the PYR concentration, [*LAC*] is the LAC concentration, *k_ldh,f_* is the forward reaction constant of lactate dehydrogenase (LDH), *k_ldh,r_* is the reverse reaction constant of lactate dehydrogenase.

In the absence of oxidative stress, the lactate fermentation efficiency is given by,

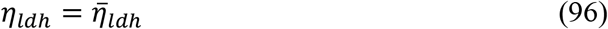

Moreover, in the presence of oxidative stress, the lactate fermentation efficiency is given by,

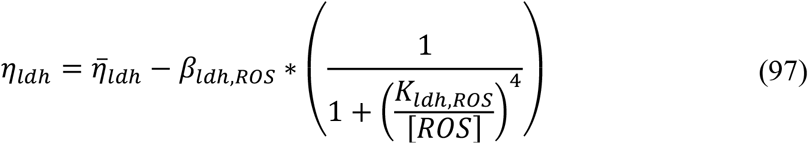

where, 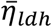 is the maximal lactate fermentation efficiency, *β_ldh,ROS_* is the maximum fractional decrease in the lactate fermentation efficiency through reactive oxygen species (*ROS*), *K_ldh,ROS_* is the threshold concentration for lactate fermentation damage by [*ROS*], [*ROS*] is the ROS concentration.

The flux of monocarboxylate transporters (*V_lac_*) is given by,

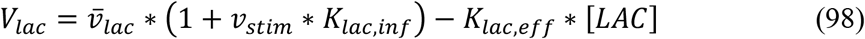

where, 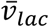 is the monocarboxylate transporters (MCTs) maximal inward flux, [*LAC*] is the LAC concentration, *v_stim_* is the stimulation pulse, *K_lac,inf_* is the coefficient constant for the inward flux of MCT, *K_lac,eff_* is the reaction constant for lactate efflux.

The flux of ATPases (*V_ATPase_*) is given by,

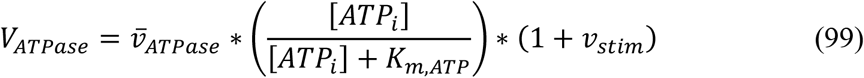

where, 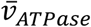 is the ATPase maximal flux, [*ATP_i_*] is the intracellular ATP concentration, *K_m,ATP_* is the affinity constant for ATP, *v_stim_* is the stimulation pulse.

The flux of the pentose phosphate pathway (*V_ppp_*) is given by,

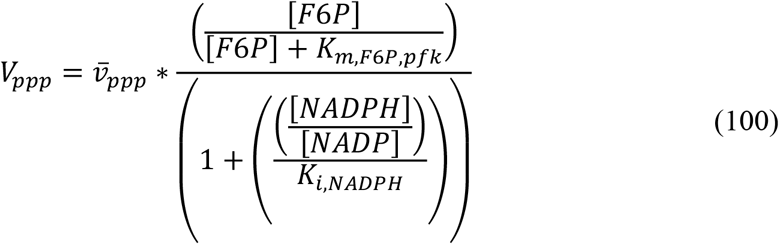

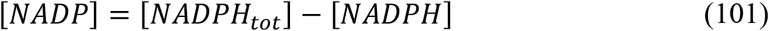

where, 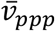 is the pentose phosphate pathway (PPP) maximal flux, [*F*6*P*] is the F6P concentration, [*NADPH*] is the NADPH concentration, [*NADP*] is the nicotinamide adenine dinucleotide phosphate (NADP) concentration, [*NADPH_tot_*] is the total NADPH and NADP concentration, *K_m,F6P,pfk_* is the affinity constant for F6P, *K_i,NADPH_* is the inhibition constant of PPP by NADPH to NADP ratio.

The flux of glutathione reductase (*V_gr_*) is given by,

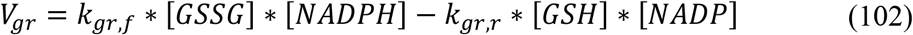

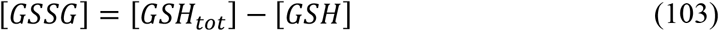

where, *k_gr,f_* is the forward reaction constant of glutathione reductase, *k_gr,r_* is the reverse reaction constant of glutathione reductase, [*NADPH*] is the NADPH concentration, [*NADP*] is the NADP concentration, [*GSH*] is the GSH concentration, *GSSG* is the GSSG concentration, [*GSH_tot_*] is the total GSH and GSSG concentration.

The flux of anti-oxidative pathway (*V_dox_*) is given by,

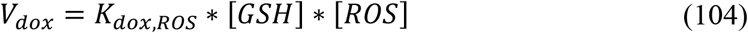

where, *K_dox,ROS_* is the reaction constant for ROS reduction by glutathione, [*GSH*] is the GSH concentration, [*ROS*] is the ROS concentration.

The flux of creatine kinase (*V_ck_*) is given by,

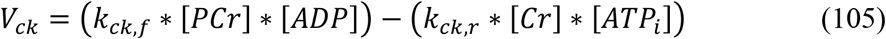

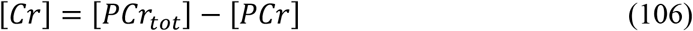

where, *k_ck,f_* is the forward reaction constant of creatine kinase, *k_ck,r_* is the reverse reaction constant of creatine kinase, [*PCr*] is the PCr concentration, [*Cr*] is the Cr concentration, [*PCr_tot_*] is the total PCr and Cr concentration, [*ADP*] is the ADP concentration, [*ATP_i_*] is the intracellular ATP concentration.

### Dopamine Turnover Processes

The whole dopamine turnover process has been modelled as a three-compartment biochemical model based on Michaelis-Menten kinetics (Tello-Bravo, 2012). The three compartments are intracellular compartment representing cytosol, extracellular compartment representing extracellular space (ECS), and vesicular compartment representing a vesicle. Previously published dopaminergic terminal models are specified in *Supplementary material-2*. In dopamine turnover processes, L-tyrosine (TYR) is converted into L-3,4-dihydroxyphenylalanine or levodopa (LDOPA) by tyrosine hydroxylase (TH) which in turn is converted into dopamine (DA) by aromatic L-amino acid decarboxylase (AADC) (Figure 5–1). The cytoplasmic DA (*DA_c_*) is stored into vesicles by vesicular monoamine transporter 2 (VMAT-2) (Figure 5–2). Upon arrival of action potential, vesicular DA (*DA_v_*) is released into extracellular space (Figure 5–3). Most of the extracellular DA (*DA_e_*) is taken up into the terminal through DA plasma membrane transporter (DAT) (Figure 5–4) and remaining extracellular DA is metabolized by catechol-O-methyltransferase (COMT) and homovanillic acid (HVA) (Figure 5–5). The DA that enters the terminal is again packed into vesicles, and the remaining cytoplasmic DA is metabolized by COMT and MAO enzymes (Figure 5–5). It is known that a DA neuron self-regulates its firing, neurotransmission and synthesis by autoreceptors (Anzalone et al., 2012; Ford, 2014). In the present model, we included autoreceptors that regulate the synthesis and release of dopamine (Figure 5–6,7). Along with TYR, external LDOPA compete for transporting into the terminal through aromatic L-amino acid transporter (AAT) (Figure 5–8).

**Figure 5:**
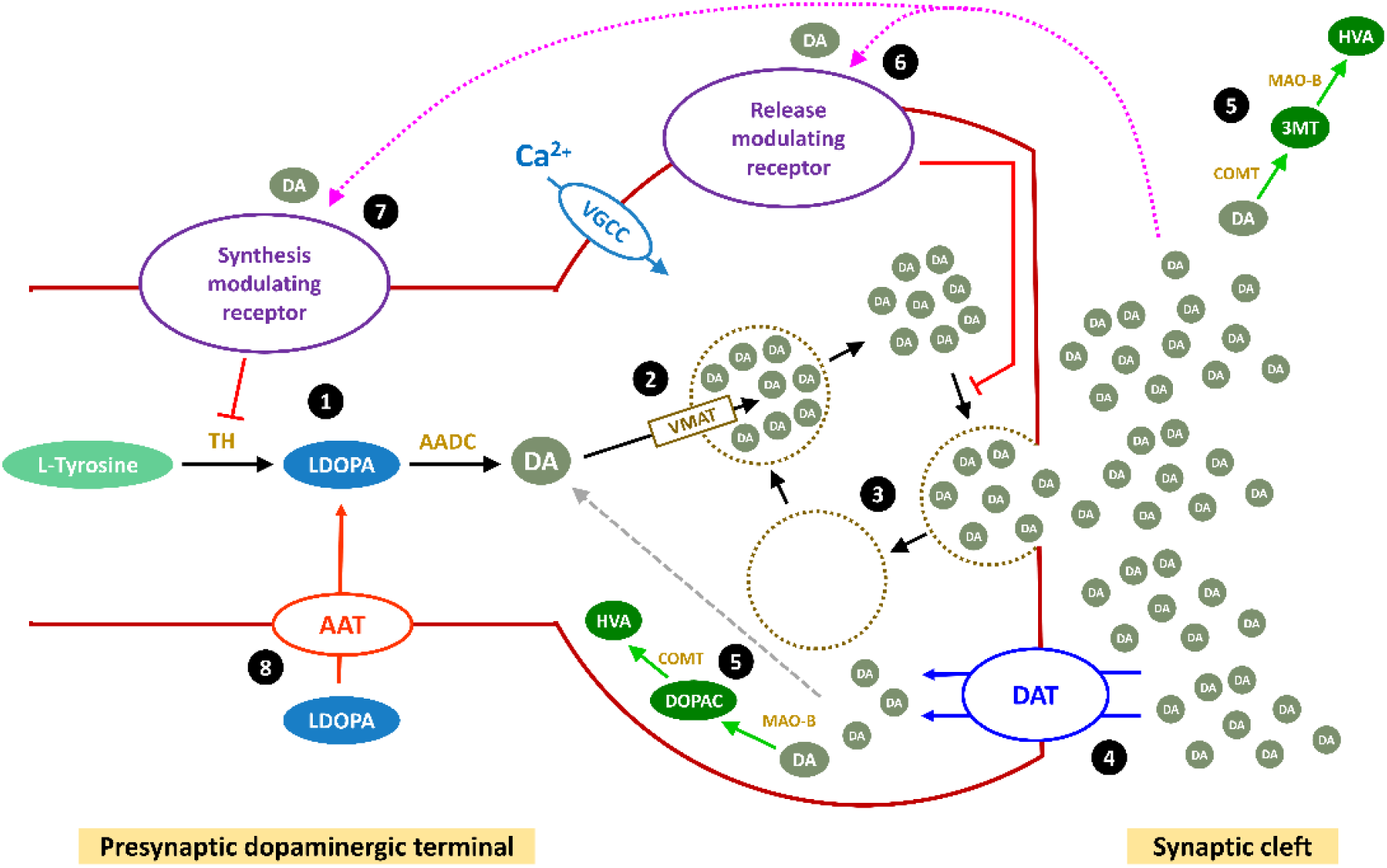
Schematic of Dopamine turnover processes in the SNc cell model. See text for description of the figure.

**Figure 6:**
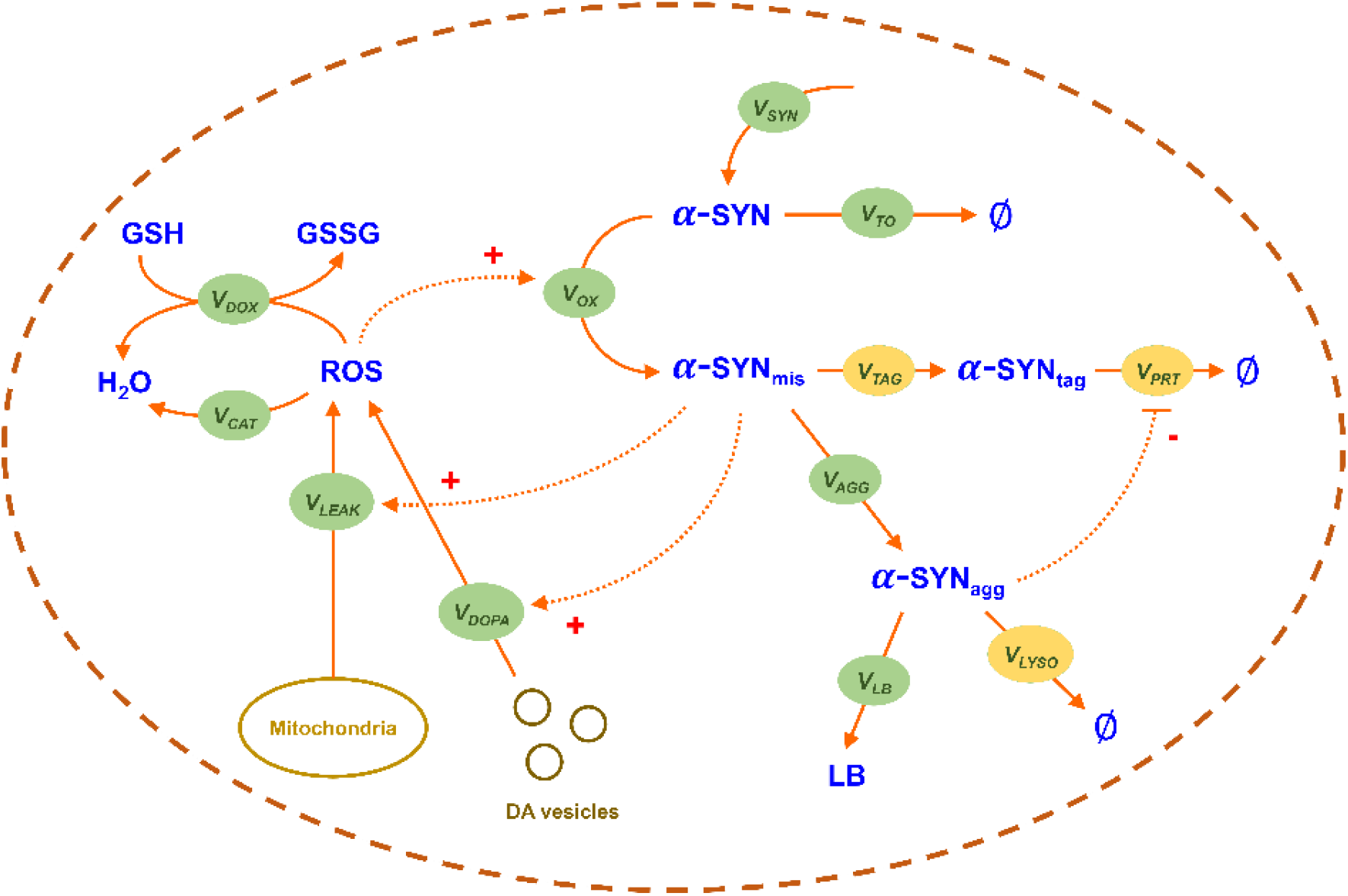
Schematic of molecular pathways in PD pathology in the SNc cell model. See text for description of the figure.

**Figure 7:**
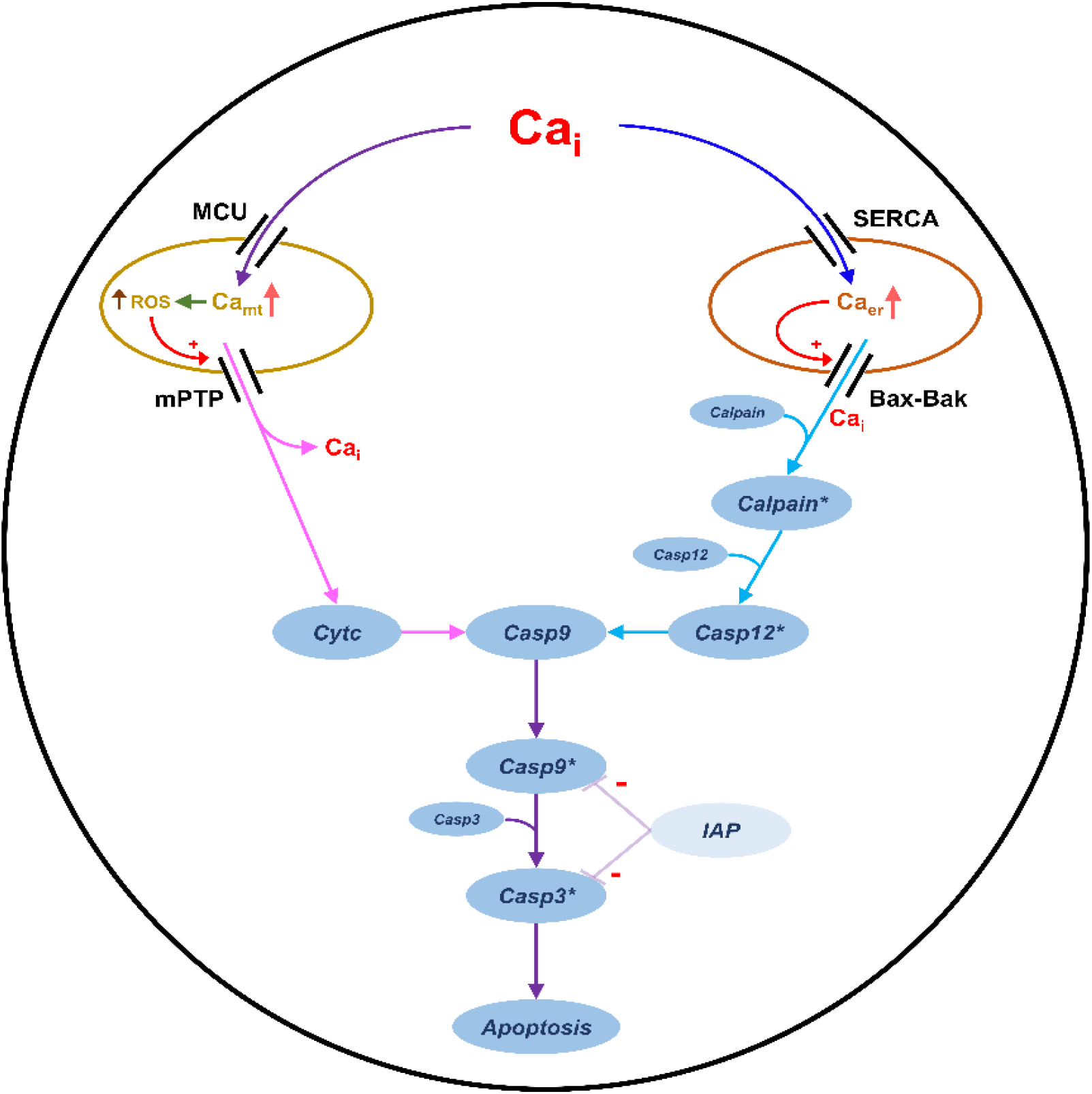
Schematic of Apoptotic pathways in SNc cell model. See text for description of the figure.

**Figure 8:**
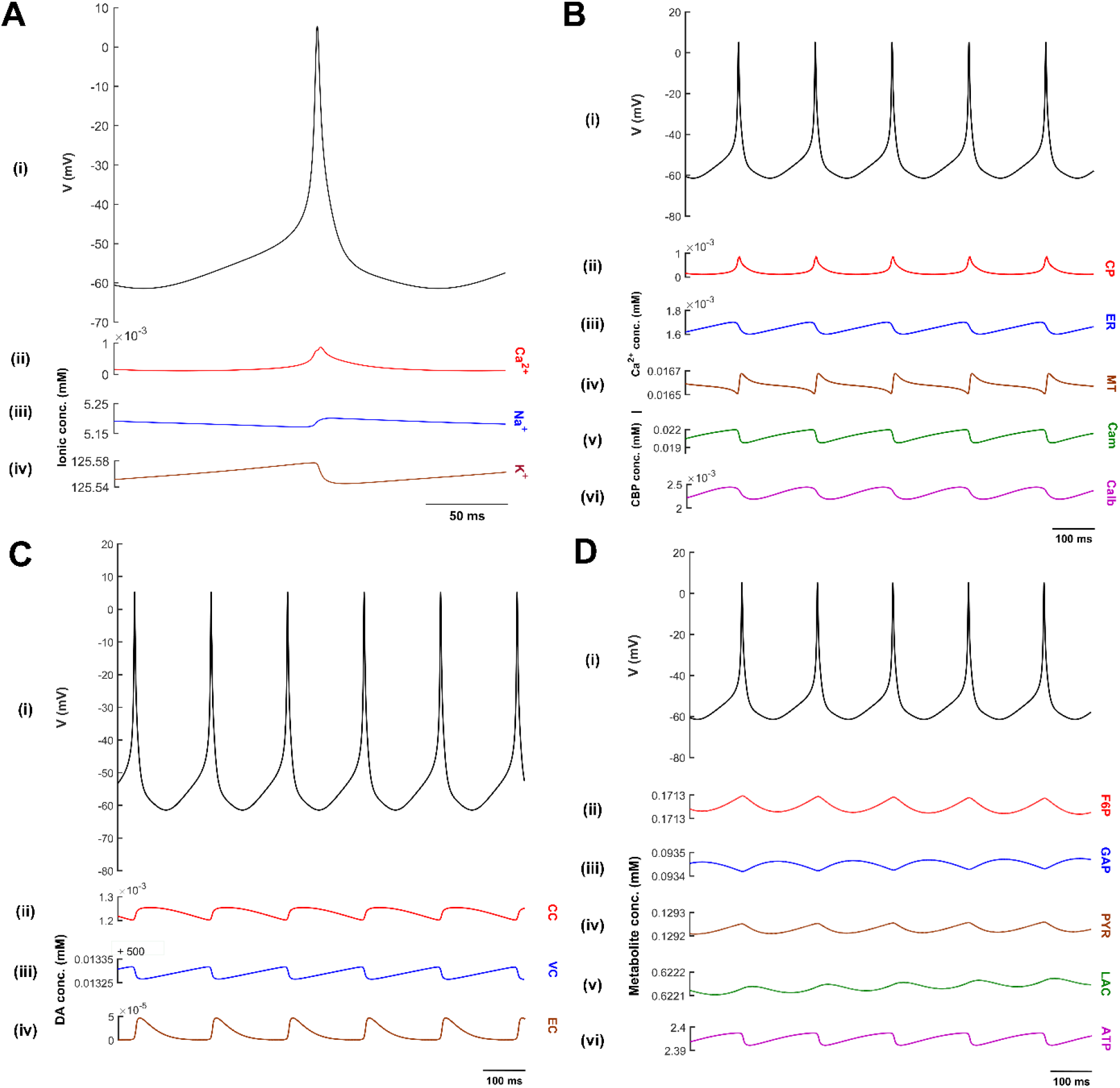
Oscillations in intracellular molecular concentrations in relation to the oscillations of the membrane potential. **(A)** Oscillations in the membrane potential (V) and the corresponding variations of intracellular sodium (Na^+^), potassium (K^+^) and calcium (Ca^2+^) concentrations, **(B)** Oscillations in cytoplasmic (CP), endoplasmic reticulum (ER) and mitochondrial (MT) calcium concentrations and calcium-binding proteins (CBP – Cam & Calb) concentration in relation to the variation of the membrane potential, **(C)** Oscillations in cytoplasmic (CC), vesicular (VC) and extracellular (EC) dopamine concentrations in relation to the membrane potential, **(D)** Oscillations in fructose-6-phosphate (F6P), glyceraldehyde-3-phosphate (GAP), pyruvate (PYR), lactate (LAC) and adenosine triphosphate (ATP) concentrations in relation to the membrane potential. Cam, Calmodulin; Calb, Calbindin; conc, concentration; mM, millimolar; mV; millivolt.

### Modelling Extracellular DA in the ECS

The three major mechanisms that determine the dynamics of extracellular DA ([*DA_e_*]) in the ECS given by,

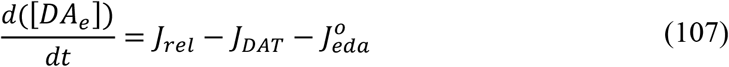

where, *J_rel_* represents the flux of calcium-dependent DA release from the DA terminal, *J_DAT_* represents the unidirectional flux of DA translocated from the extracellular compartment (ECS) into the intracellular compartment (cytosol) via DA plasma membrane transporter (DAT), and 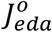 represents the outward flux of DA degradation, which clears DA from ECS.

### Calcium-Dependent DA Release Flux

Assuming that calcium-dependent DA release occurs within less than a millisecond after the calcium channels open, the flux of DA release (*J_rel_*) from the DA terminal is given by,

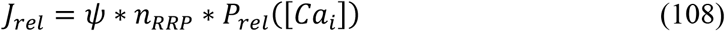

where, [*Ca_i_*] is the intracellular calcium concentration in the DA terminal, *P_rel_* is the release probability as a function of intracellular calcium concentration, *n_RRP_* is the average number of readily releasable vesicles and *ψ* is the average release flux per vesicle within a single synapse.

The flux of calcium-dependent DA release depends on extracellular DA concentration, and intracellular ATP acts as a feedback mechanism, assuming this regulation as extracellular DA and intracellular ATP controls the number of vesicles in the readily releasable vesicle pool (*n_RRP_*).

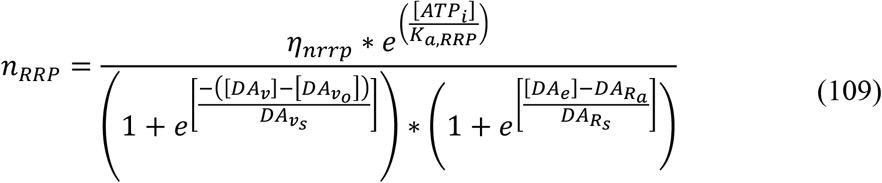

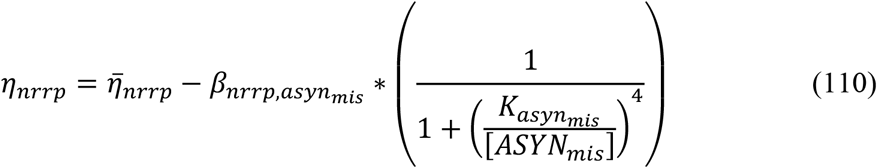

where, [*DA_v_o__*] is the initial vesicular DA concentration, *DA_v_s__* is the sensitivity to vesicular concentration, *DA_R_a__* is the high-affinity state for DA binding to receptors and *DA_R_s__* is the binding sensitivity, [*ATP_i_*] is the intracellular ATP concentration, *K_a,RRP_* is the activation constant for ATP, *η_nrrp_* is the effect of misfolded alpha-synuclein on vesicle recycling (Venda et al., 2010), 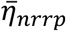 is the maximal vesicle recycling efficiency, *β_nrrp,asyn_mis__* is the maximum fractional decrease in the vesicle recycling efficiency through *ASYN_mis_, K_asyn_mis__* is the threshold concentration for damage by *ASYN_mis_*, [*ASYN_mis_*] is the misfolded alpha-synuclein concentration.

The release probability of DA as a function of intracellular calcium concentration is given by,

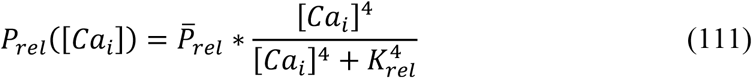

where, 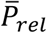 is the maximum release probability and *K_rel_* is the sensitivity of calcium concentration, [*Ca_i_*] is the intracellular calcium concentration.

### Unidirectional Reuptake Flux of DA

The unidirectional reuptake flux of extracellular DA into the presynaptic terminal is given by,

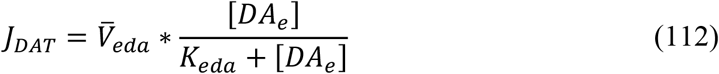

where, 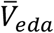 is the maximal velocity of dopamine transporter (DAT), *K_eda_* is the DA concentration at half-maximal velocity, [*DA_e_*] is the extracellular DA concentration.

### Outward Extracellular Flux

The flux of extracellular DA enzymatic degradation in the synaptic cleft (ECS) is given by,

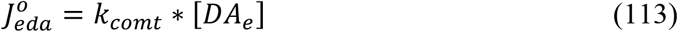

where, *k_comt_* is the rate at which extracellular DA cleared from ECS, [*DA_e_*] is the extracellular DA concentration.

### Modelling Intracellular DA in the Terminal

The intracellular DA dynamics ([*DA_i_*]) is determine as the sum of dopamine concentration in cytosolic and vesicular compartments is given by,

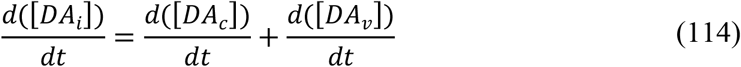

The cytosolic DA dynamics ([*DA_c_*]) is given by,

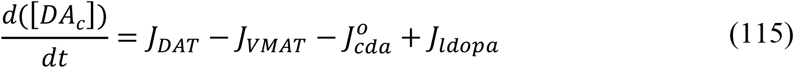

where, *J_DAT_* represents the unidirectional flux of DA translocated from ECS into the cytosol through DAT, *J_VMAT_* represents the flux of cytosolic DA into vesicle through vesicular monoamine transporters (VMAT), 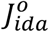 represents the outward flux of DA degradation which clears DA from the cytosol, *J_ldopa_* represents the flux of synthesized cytosol DA from levodopa (LDOPA) drug therapy.

The vesicular DA dynamics ([*DA_v_*]) is given by,

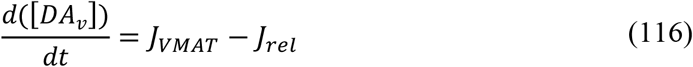

where, *J_rel_* represents the flux of calcium-dependent DA release from the DA terminal, *J_VMAT_* represents the flux of cytosolic DA into a vesicle.

### LDOPA Synthesis Flux

The flux of synthesized LDOPA whose velocity is the function of intracellular calcium concentration and LDOPA synthesis is regulated by the substrate (TYR) itself, extracellular DA (via autoreceptor) and intracellular DA concentrations are given by,

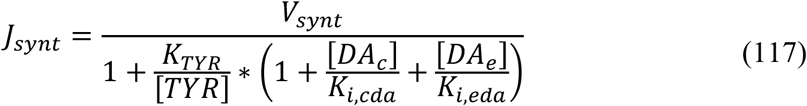

where, *V_synt_* is the velocity of synthesizing LDOPA, [*TYR*] is the tyrosine concentration in terminal bouton, *K_TYR_* is the tyrosine concentration at which half-maximal velocity was attained, *K_i,cda_* is the inhibition constant on *K_TYR_* due to cytosolic DA concentration, *K_i,eda_* is the inhibition constant on *K_TYR_* due to extracellular DA concentration, [*DA_c_*] is the cytoplasmic DA concentration, [*DA_e_*] is the extracellular DA concentration.

In (Chen et al., 2003) neuronal stimulation was linked to DA synthesis through an indirect event which starts with calcium influx into the terminal bouton. In this model, the velocity of LDOPA synthesis as a function of calcium levels in the terminal bouton is expressed as,

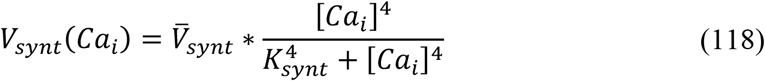

where, *K_synt_* is the calcium sensitivity, 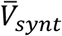 is the maximal velocity for LDOPA synthesis, [*Ca_i_*] is the intracellular calcium concentration.

### Storage Flux of DA into the Vesicle

The flux of transporting DA in the cytosol into the vesicles which depend on the intracellular ATP is given by,

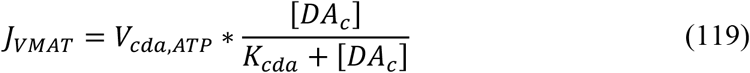

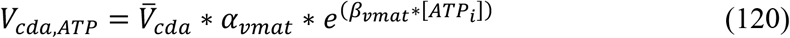

where, *K_cda_* is the cytosolic DA concentration at which half-maximal velocity was attained, 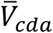 is the maximal velocity with which DA was packed into vesicles, [*DA_c_*] is the cytosolic DA concentration, *α_vmat_* is the scaling factor for VMAT, *β_vmat_* is the scaling factor for *ATP_i_*, [*ATP_i_*] is the intracellular ATP concentration.

### Outward Intracellular Flux

The flux of intracellular DA enzymatic degradation in synaptic bouton (cytosol) is given by,

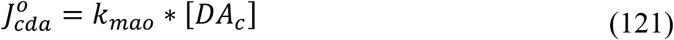

where, *k_mao_* is the rate at which intracellular DA cleared from the cytosol, [*DA_c_*] is the cytosolic DA concentration.

### LDOPA to DA Conversion Flux

The flux of LDOPA conversion to DA by aromatic L-amino acid decarboxylase (AADC) (Reed et al., 2012) is given by,

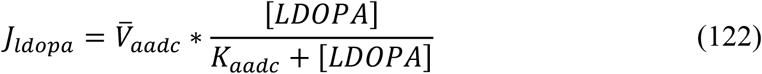

where, *K_aadc_* is the LDOPA concentration at which half-maximal velocity was attained, 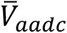 is the maximal velocity with which LDOPA was converted to DA, [*LDOPA*] is the LDOPA concentration.

### Transport Flux of Exogenous LDOPA into the Terminal

The flux of exogenous LDOPA transported into the terminal through aromatic L-amino acid transporter (AAT) while competing with other aromatic amino acids (Reed et al., 2012) is given by,

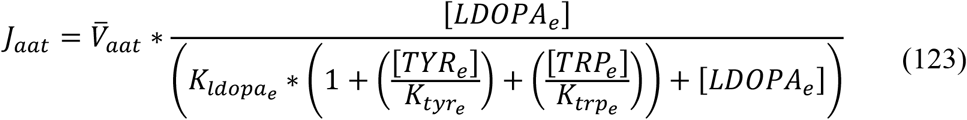

where, *K_ldopa_e__* is the extracellular LDOPA concentration at which half-maximal velocity was attained, 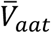 is the maximal velocity with which extracellular LDOPA was transported into the cytosol, [*LDOPA_e_*] is the extracellular LDOPA concentration, [*TYR_e_*] is the extracellular TYR concentration, [*TRP_e_*] is the extracellular tryptophan (TRP) concentration, *K_tyr_e__* is the affinity constant for [*TYR_e_*], *K_trp_e__* is the affinity constant for [*TRP_e_*].

When LDOPA drug therapy initiated,

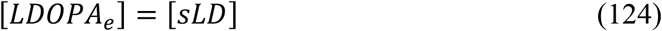

Moreover, when no LDOPA drug therapy initiated,

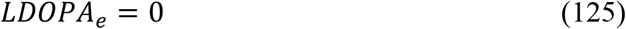

The LDOPA concentration ([*LDOPA*]) dynamics inside the terminal is given by,

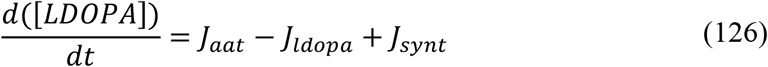

where, *J_aat_* represents the flux of exogenous LDOPA transported into the cytosol, *J_ldopa_* represents the conversion flux of exogenous LDOPA into DA, *J_synt_* represents the flux of synthesized LDOPA from tyrosine, [*sLD*] is the serum LDOPA concentration.

### Molecular Pathways Involved in PD Pathology

The molecular pathways in PD pathology were adapted from (Cloutier and Wellstead, 2012) and incorporated in the comprehensive model of SNc cell. The ROS formation occurs due to leakage from mitochondria during oxidative phosphorylation for ATP production, auto-oxidation of excess freely available DA in the cytoplasm, and misfolded alpha-synuclein (*ASYN_mis_*). In the present model, excess ROS is scavenged by glutathione. Under pathological conditions such as elevated ROS levels, normal alpha-synuclein (*ASYN*) undergoes conformation changes into misfolded alpha-synuclein. The misfolded alpha-synuclein is tagged (*ASYN_tag_*) and degraded by the ubiquitous-proteasome pathway using ATP. Excess misfolded alpha-synuclein forms aggregates, which in turn gets degraded by the lysosomal degradation pathway using ATP. In some scenarios, these alpha-synuclein aggregates (*ASYN_agg_*) form Lewy bodies (*LB*).

The model consists of ROS formation from different processes, ROS scavenging mechanism, alpha-synuclein aggregation, proteasomal and lysosomal degradation of damaged protein, etc. The following equations give a concise view of all metabolite dynamics in the PD pathology pathways,

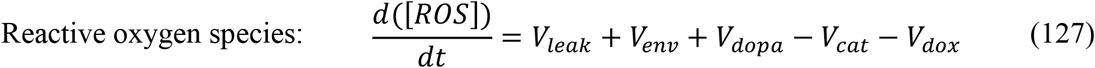

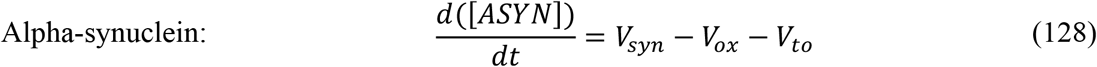

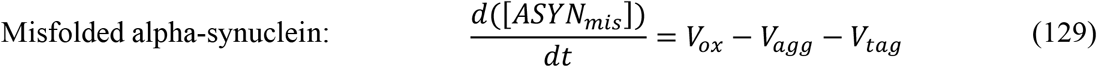

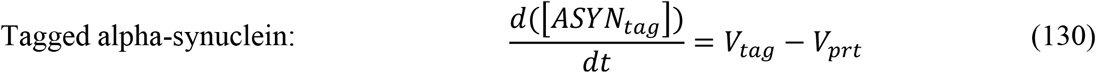

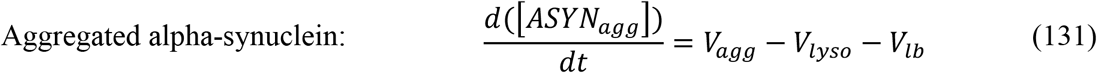

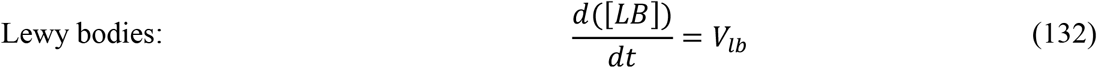

where, *V_leak_* is the flux of oxidative stress due to mitochondrial leakage, *V_env_* is the flux of external oxidative stress, *V_dopa_* is the flux of oxidative stress due to excess cytoplasmic dopamine, *V_cat_* is the catabolizing flux of ROS by catalase enzyme, *V_dox_* is the flux of GSH-dependent ROS scavenging pathway, *V_syn_* is the synthesizing flux of alpha-synuclein protein, *V_ox_* is the flux of alpha-synuclein misfolding due to ROS, *V_to_* is the usage flux of alpha-synuclein in other processes, *V_agg_* is the flux of alpha-synuclein aggregation, *V_tag_* is the flux of ATP-dependent ubiquitination of damaged protein for proteasomal degradation, *V_prt_* is the flux of ATP-dependent breakdown of damaged protein through proteasomal degradation, *V_lyso_* is the flux of ATP-dependent breakdown of aggregated protein through lysosomal degradation, *V_lb_* is the flux of Lewy bodies formation.

The flux of oxidative stress due to mitochondrial leakage (*V_leak_*) is given by,

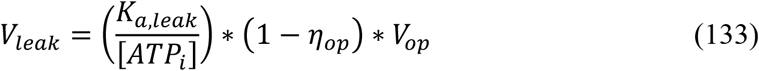

where, *V_op_* is the flux of the oxidative phosphorylation pathway, *η_op_* is the electron transport chain efficiency, [*ATP_i_*] is the intracellular ATP concentration, *K_a,ATP_* is the activation constant for ATP.

The flux of oxidative stress due to excess dopamine in the cytoplasm (*V_dopa_*) is given by,

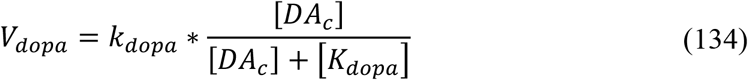

where, *k_dopa_* is the reaction constant for ROS production by excess dopamine, [*DA_c_*] is the cytoplasmic dopamine concentration, *K_dopa_* is the affinity constant for [*DA_c_*].

The catabolizing flux of ROS by catalase enzyme (*V_cat_*) is given by,

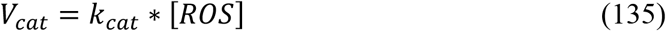

where, *k_cat_* is the reaction constant for catalase, [*ROS*] is the ROS concentration.

The synthesizing flux of alpha-synuclein protein (*V_syn_*) is given by,

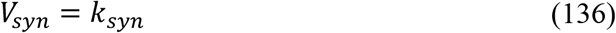

where, *k_syn_* is the reaction constant for alpha-synuclein synthesis.

The flux of alpha-synuclein misfolding due to ROS (*V_ox_*) is given by,

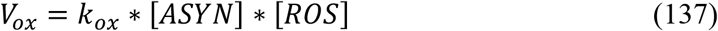

where, *k_ox_* is the reaction constant for alpha-synuclein oxidation, [*ASYN*] is the alpha-synuclein concentration, [*ROS*] is the ROS concentration.

The usage flux of alpha-synuclein in other processes (*V_to_*) is given by,

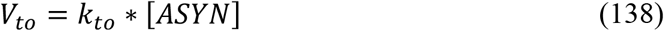

where, *k_to_* is the reaction constant for alpha-synuclein consumption, [*ASYN*] is the alpha-synuclein concentration.

The flux of alpha-synuclein aggregation (*V_agg_*) is given by,

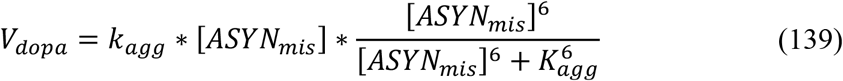

where, *k_agg_* is the reaction constant for alpha-synuclein aggregation, [*ASYN_mis_*] is the misfolded alpha-synuclein concentration, *K_agg_* is the affinity constant for [*ASYN_mis_*].

The flux of ATP-dependent ubiquitination of damaged protein for proteasomal degradation (*V_tag_*) is given by,

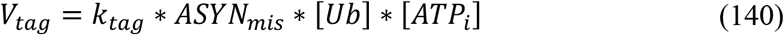

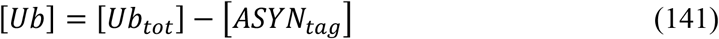

where, *k_tag_* is the reaction constant for ubiquitination of damaged protein, [*ASYN_mis_*] is the misfolded alpha-synuclein concentration, [*Ub*] is the ubiquitin concentration, [*ATP_i_*] is the intracellular ATP concentration, [*Ub_tot_*] is the total ubiquitin concentration, [*ASYN_tag_*] is the tagged alpha-synuclein concentration.

The flux of ATP-dependent breakdown of damaged protein through proteasomal degradation (*V_prt_*) is given by,

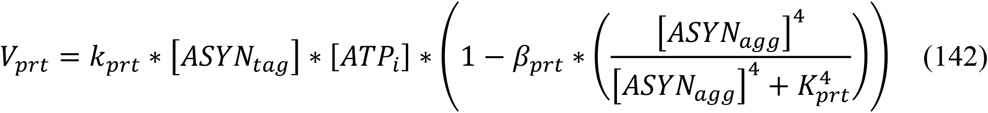

where, *k_prt_* is the reaction constant for damaged protein disposal by the proteasome, [*ASYN_tag_*] is the tagged alpha-synuclein concentration, [*ATP_i_*] is the intracellular ATP concentration, [*ASYN_agg_*] is the aggregated alpha-synuclein concentration, *K_prt_* is the affinity constant for [*ASYN_agg_*], *β_prt_* is the fraction reduction of proteasome activity by [*ASYN_agg_*].

The flux of ATP-dependent breakdown of aggregated protein through lysosomal degradation (*V_lyso_*) is given by,

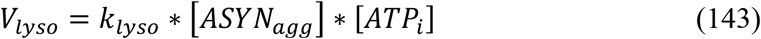

where, *k_lyso_* is the reaction constant for [*ASYN_agg_*] disposal by the lysosome, [*ATP_i_*] is the intracellular ATP concentration.

The flux of Lewy body formation (*V_lb_*) is given by,

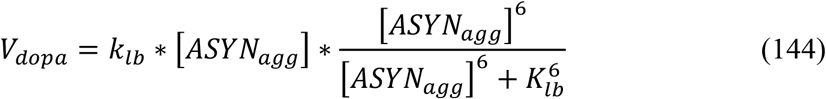

where, *k_lb_* is the reaction constant for Lewy bodies from [*ASYN_agg_*], [*ASYN_agg_*] is the aggregated alpha-synuclein concentration, *K_prt_* is the affinity constant for [*ASYN_agg_*].

### Apoptotic Pathways

The apoptotic pathways were adapted from (Hong et al., 2012) and incorporated in the comprehensive model of SNc cell. The model consists of ER stress-induced apoptotic activation and mitochondrial ROS-induced apoptotic activation (Figure 7) (El-Osta and Circu, 2016).

Under stress conditions, calcium from endoplasmic reticulum efflux and intracellular calcium (*Ca_i_*) builds up in the cytoplasm of SNc neurons which activates calcium-dependent calpain (*Calpain*) protease through ER stress-induced pathway (Rasheva and Domingos, 2009). Activated calpain (*Calpain*\*) proteases procaspase-12 (*Casp*12) to caspase-12 (*Casp*12*) through calpain-dependent activation of caspase-12 (Martinez et al., 2010). Activated caspase-12 cleaves procaspase-9 (*Casp*9) into caspase-9 (*Casp*9*) through cytochrome c-independent pathway (Morishima et al., 2002), caspase-9, in turn, activates procaspase-3 (*Casp*3) into caspase-3 (*Casp3*\*) (Brentnall et al., 2013). Activated caspase-3 eventually induces apoptotic mediators (*Apop*) (Porter and Jänicke, 1999).

Under stress conditions, the mitochondrial permeability increases through mitochondrial permeability transition pore complex (*PTP_mit_*) which leads to release of pro-apoptotic factors into the cytosol (Redza-Dutordoir and Averill-Bates, 2016) results in cytochrome c-dependent (*Cytc*) activation of apoptotic mediator caspase-9 (Jiang and Wang, 2004). Activated caspase-9, in turn, activates procaspase-3 (*Casp*3) into caspase-3 (*Casp*3*) (Brentnall et al., 2013). Activated caspase-3 eventually induces apoptotic mediators (*Apop*) (Porter and Jänicke, 1999).

*ER Stress-Induced Apoptosis:*

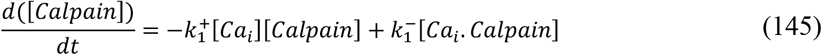

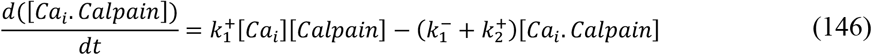

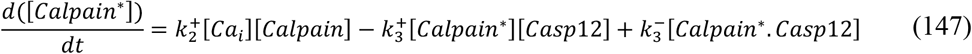

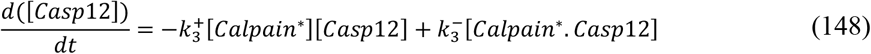

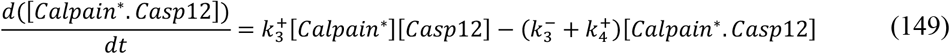

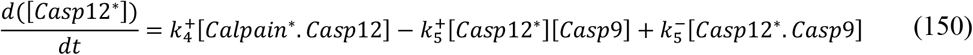

*MT-Induced Apoptosis:*

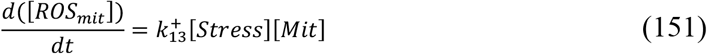

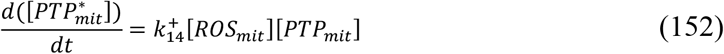

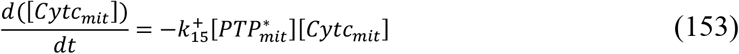

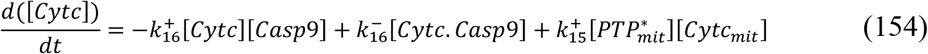

*Common Pathways for Both Apoptotic Signaling Pathways:*

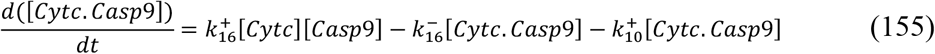

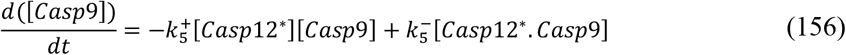

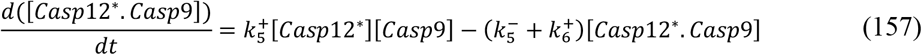

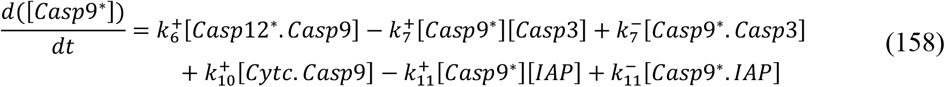

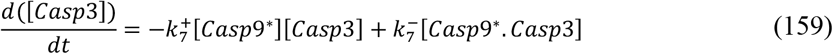

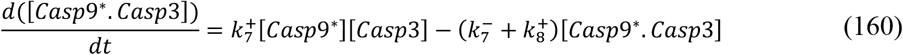

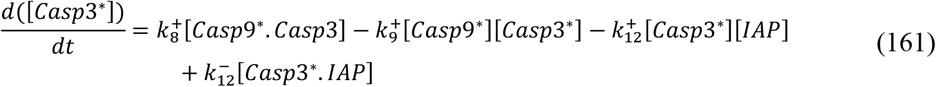

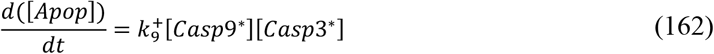

*Inhibitor of Apoptosis (IAP) Proteins:*

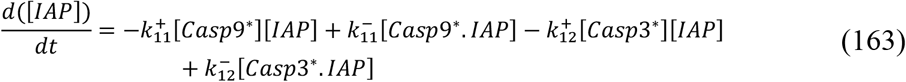

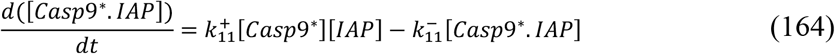

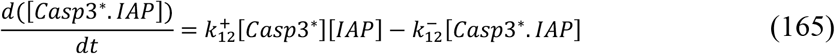

### Energy Consumption

The approximate ATP consumption in the propagation of action potential and recovery of membrane potential (*uATP_ep_*) is given by,

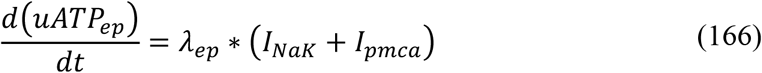

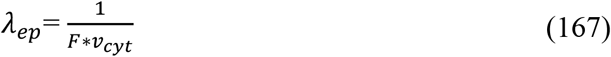

where, *λ_ep_* is the scaling factor for electrical processes, *I_NaK_* is the sodium-potassium pump current, *I_pmca_* is the calcium pump current, *F* is the Faraday’s constant, *v_cyt_* is the cytosolic volume.

The approximate ATP consumption in synaptic recycling and neurotransmitter packing into vesicles (*uATP_sp_*) is given by,

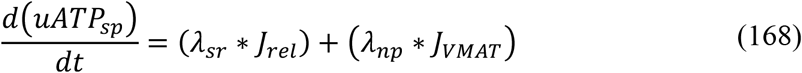

where, *λ_sr_* is the scaling factor for synaptic recycling, *λ_np_* is the scaling factor for neurotransmitter packing, *J_rel_* is the DA release flux from the terminal, *J_VMAT_* is the DA packing flux into the vesicles.

The approximate ATP consumption in calcium influx into the endoplasmic reticulum (*uATP_er_*) is given by,

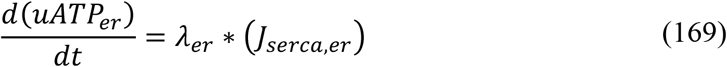

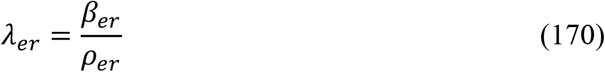

where, *λ_er_* is the scaling factor for endoplasmic reticulum processes, *J_serca,er_* is the calcium influx into endoplasmic reticulum through SERCA, *β_er_* is the ratio of free calcium to total calcium concentration in the ER, *ρ_er_* is the volume ratio between the ER and cytosol.

The approximate ATP consumption in damaged protein disposal mechanisms (*uATP_dm_*) is given by,

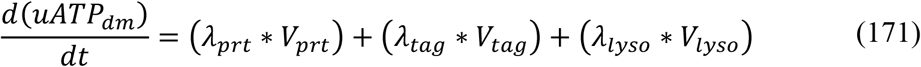

where, *λ_prt_* is the scaling factor for proteasomal degradation of damaged protein, *λ_tag_* is the scaling factor for ubiquitination of damaged protein, *λ_lyso_* is the scaling factor for lysosomal degradation of damaged protein, *V_prt_* is the flux of ATP-dependent breakdown of damaged protein through proteasomal degradation, *V_tag_* is the flux of ATP-dependent ubiquitination of damaged protein for proteasomal degradation, *V_lyso_* is the flux of ATP-dependent breakdown of aggregated protein through lysosomal degradation. All the initial values of the differential equations were taken as zero. All parameter and steady state values are given in *Supplementary material-3*.

## RESULTS

We developed a comprehensive model of SNc neuron which exhibits characteristic ionic dynamics (Figure 8-A), calcium dynamics (Figure 8-B), dopamine dynamics (Figure 8-C) and energy metabolite dynamics (Figure 8-D). The model also exhibits energy consumption by different cellular processes (Figure 9-A) and varying dopamine released extracellularly based on nRRP (Figure 9-B).

**Figure 9:**
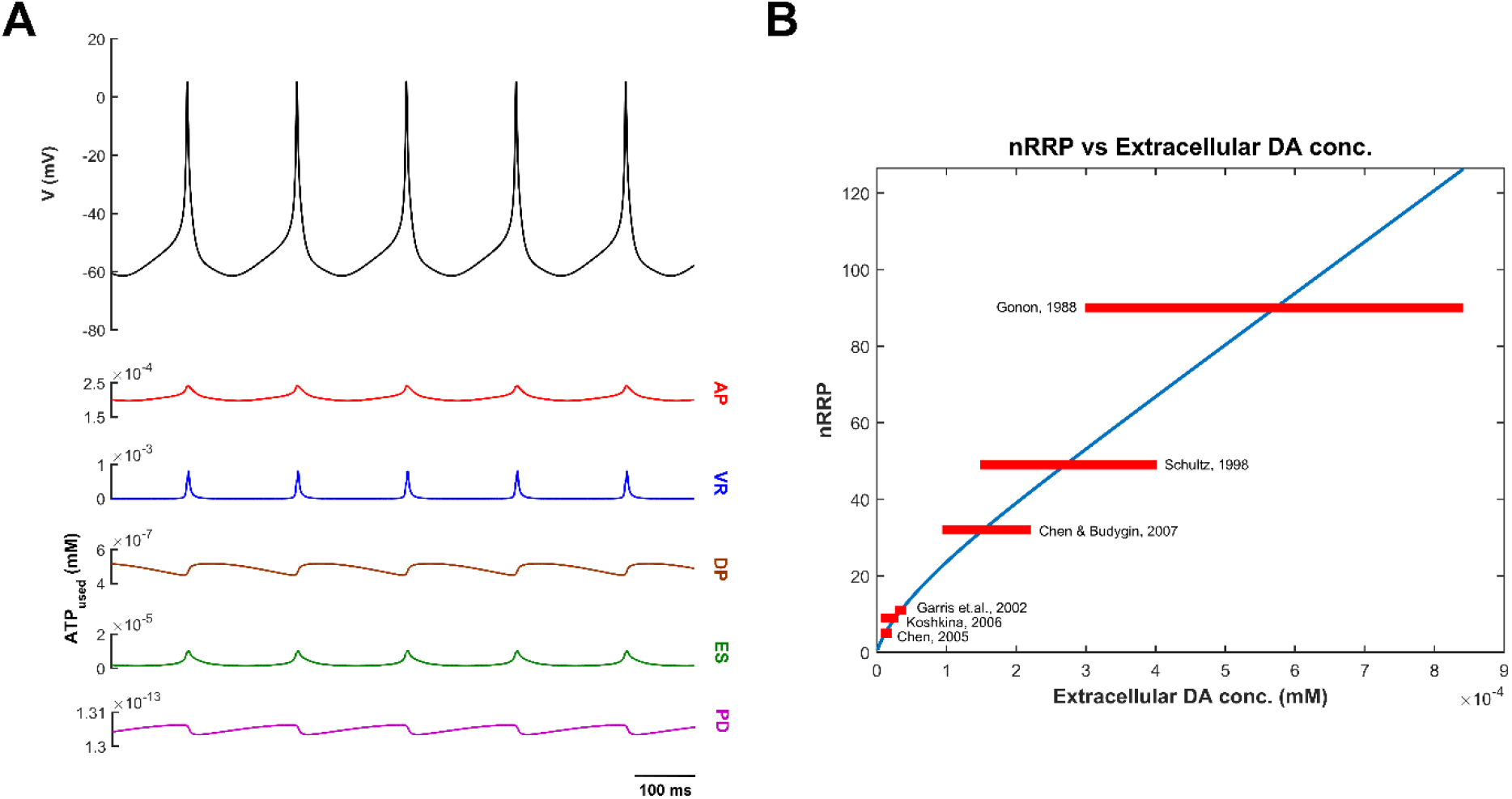
Energy consumption and extracellular dopamine release. **(A)** Energy consumption by different cellular processes in the SNc cell, **(B)** Range bar plot of extracellular dopamine (DA) concentration with respect to nRRP. ATP, Adenosine Triphosphate; AP, Action potential Propagation; VR, Vesicle Recycling; DP, Dopamine Packing; ES, Endoplasmic reticulum calcium Sequestering; PD, Protein Degradation; nRRP, number of Readily Releasable vesicle Pool.

Then, we studied the effect of electrical (Figure 10) and chemical (Figure 11) stimulation on the proposed model. Finally, we showed model responses to energy deficiency conditions (Figure 12, 13, 14).

**Figure 10:**
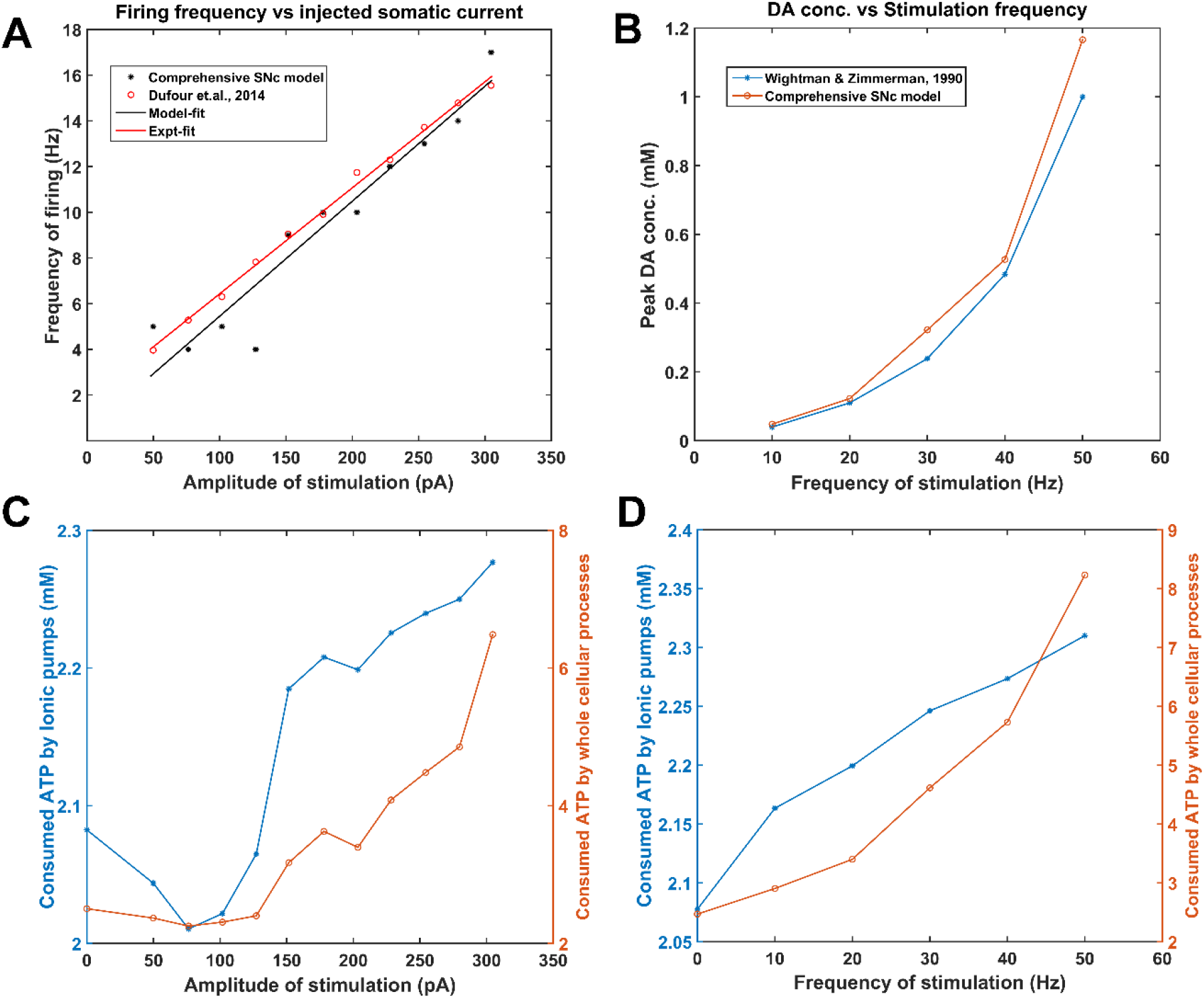
Model response to electrical stimulation. Frequency of firing **(A)** and Energy consumption **(C)** by ionic pumps (blue trace) and whole cellular processes (orange trace) of the model concerning the amplitude of stimulating depolarized current (1 sec), Extracellular dopamine (DA concentration **(B)** and Energy consumption **(D)** by ionic pumps (blue trace) and whole cellular processes (orange trace) of the model concerning the frequency of stimulating depolarized current (2 sec). ATP, Adenosine Triphosphate.

**Figure 11:**
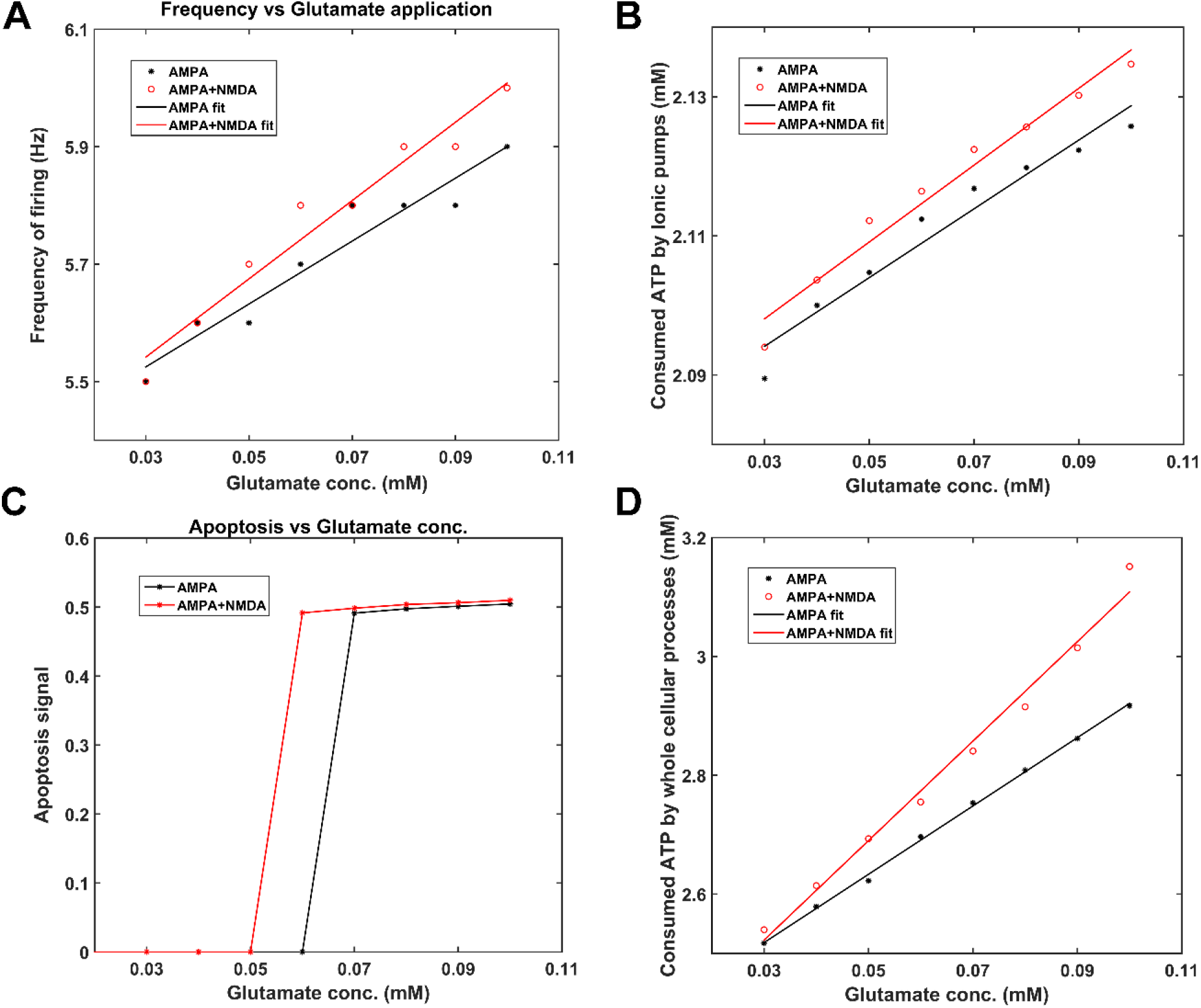
Model response to chemical stimulation (glutamate). Frequency of firing **(A)**, Apoptosis signal **(C)** due to excess stimulation, Energy consumption by ionic pumps **(B)** and whole cellular processes **(D)** of the model concerning the concentration of glutamate application (1 sec). ATP, Adenosine Triphosphate; AMPA, Alpha-amino-3-hydroxy-5-Methyl-4-isoxazole Propionic Acid; NMDA, N-Methyl-D-aspartic Acid.

**Figure 12:**
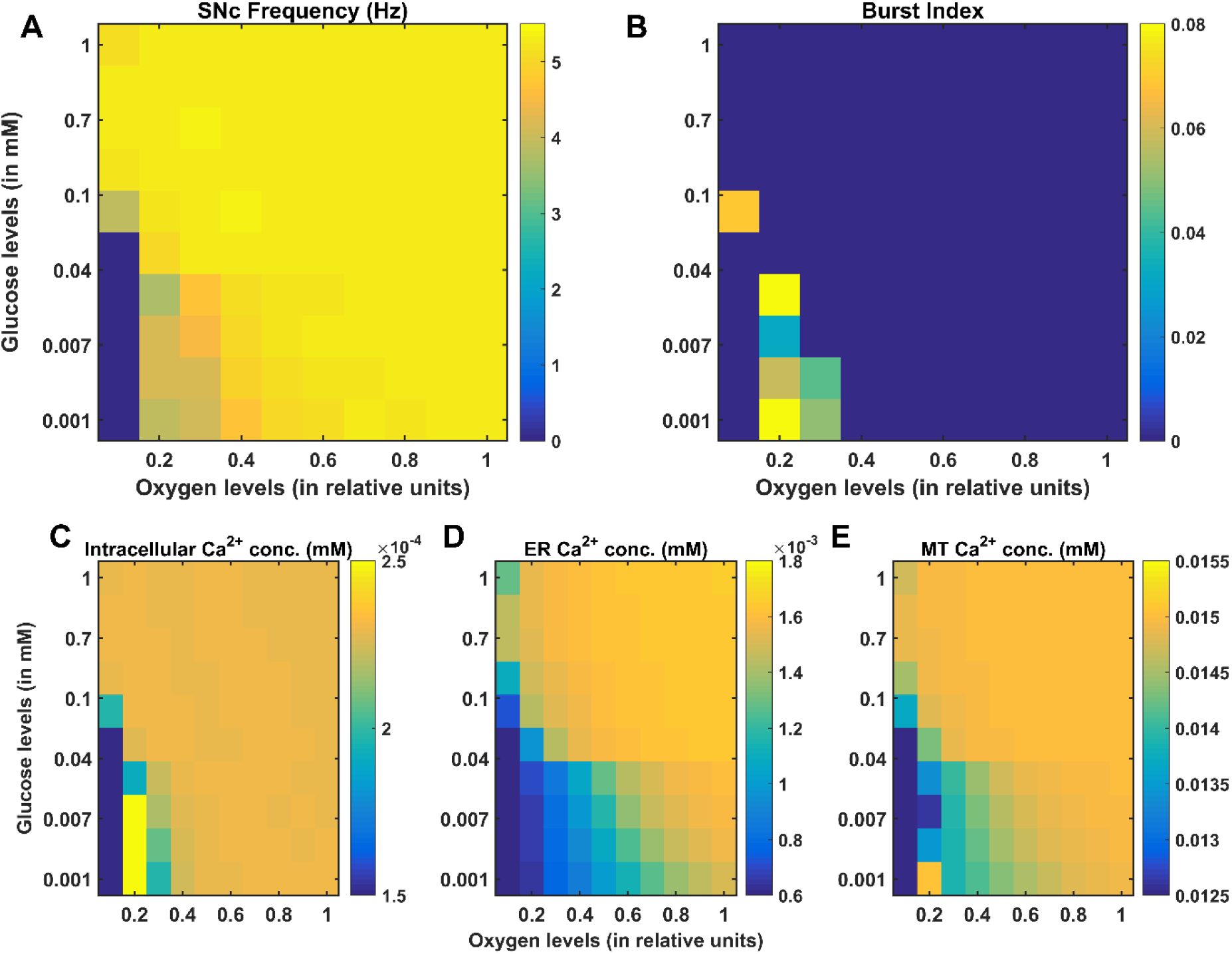
Model response to hypoglycemia and hypoxia conditions. Average frequency of firing **(A)**, Bursting **(B)**, average intracellular calcium (Ca^2+^) concentration **(C)**, average endoplasmic reticulum (ER) calcium concentration **(D)**, and average mitochondrial (MT) calcium concentration **(E)** of the model for varying glucose and oxygen concentrations. SNc, Substantia Nigra pars compacta; conc, concentration; mM, millimolar.

**Figure 13:**
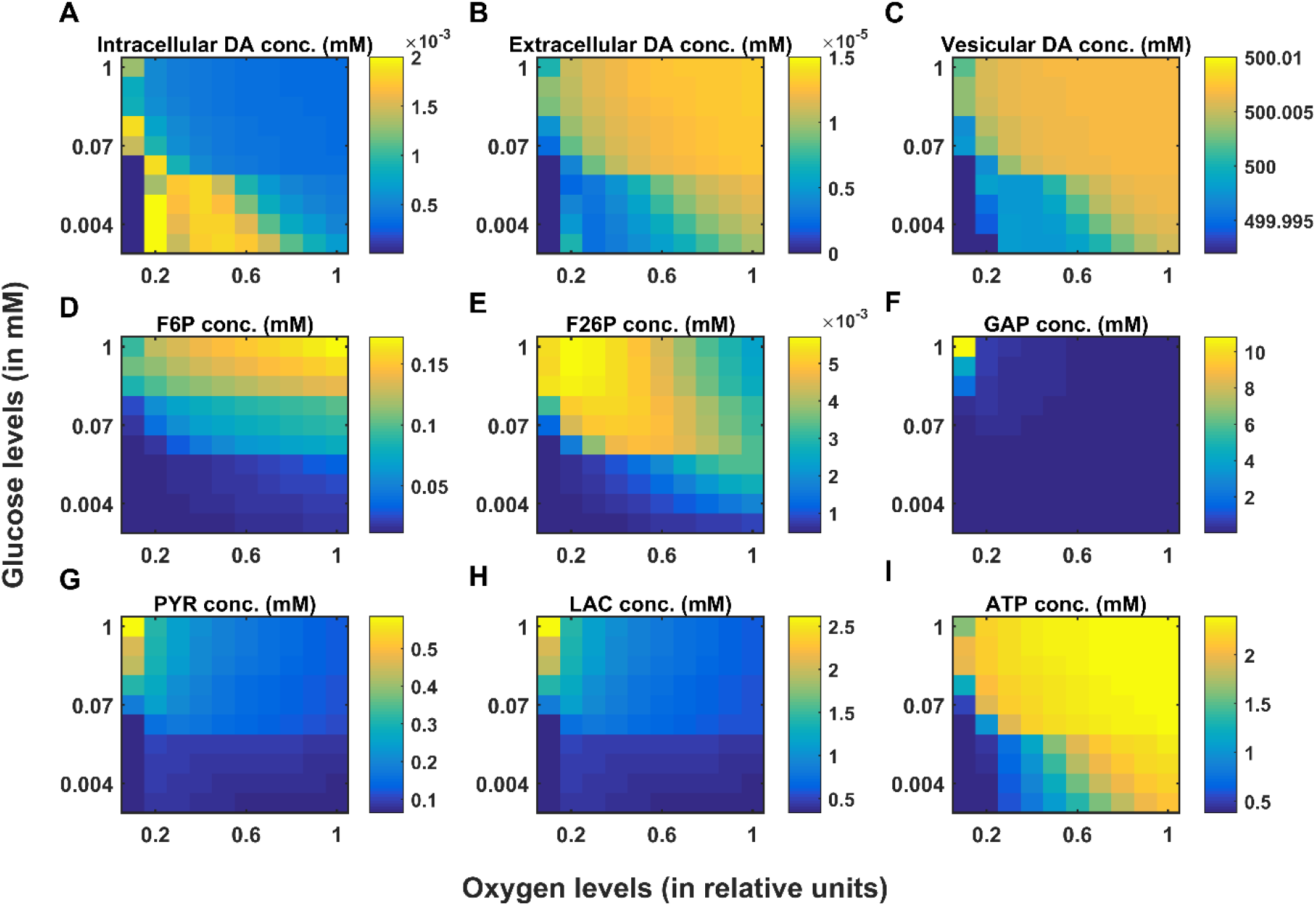
Model response to hypoglycemia and hypoxia conditions. Average intracellular dopamine (DA) concentration **(A)**, average extracellular dopamine concentration **(B)**, average vesicular dopamine concentration **(C)**, average fructose-6-phosphate (F6P) concentration **(D)**, average fructose-2,6-biphosphate (F26P) concentration **(E)**, average glyceraldehyde-3-phosphate (GAP) concentration **(F)**, average pyruvate (PYR) concentration **(G)**, average lactate (LAC) concentration **(H)**, average adenosine triphosphate (ATP) concentration **(I)** of the model for varying glucose and oxygen concentrations. conc, concentration; *mM*, millimolar.

**Figure 14:**
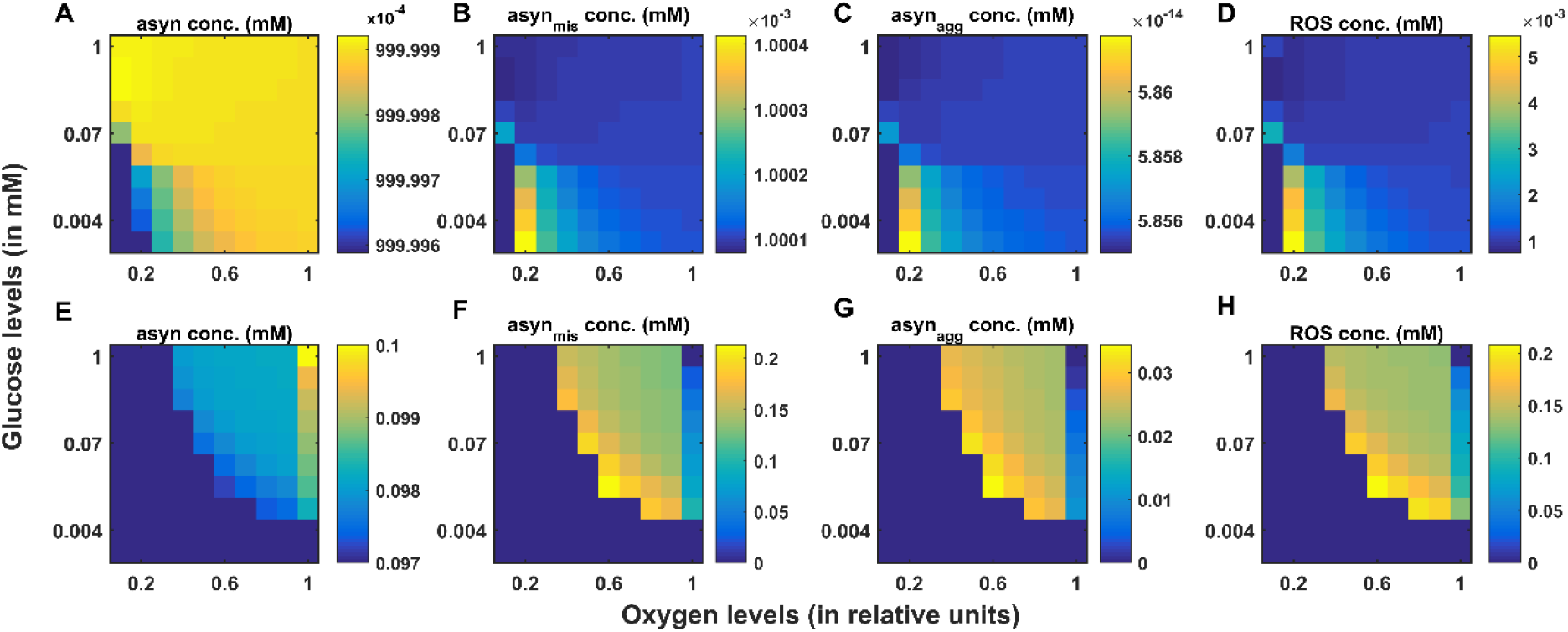
Whole (A, B, C, D) and reduced (E, F, G, H) models responses to hypoglycemia and hypoxia conditions. Average normal alpha-synuclein (α-syn) concentration **(A, E)**, average misfolded alpha-synuclein (α-syn_mis_) concentration **(B, F)**, average aggregated alpha-synuclein (α-syn_agg_) concentration **(C, G)**, and average reactive oxygen species (ROS) concentration **(D, H)** of the fast and slow dynamic models for varying glucose and oxygen concentrations. conc, concentration; mM, millimolar.

### Characteristic Ionic Dynamics of the SNc Neuron

The proposed comprehensive model of SNc exhibits the basal firing rate of 5 *Hz*, which is in the range of 3 to 8 *Hz* observed experimentally (Figure 8) (Grace and Bunney, 1984b). The bursting type of firing also observed in the proposed model with a different range of synaptic inputs (not shown here) (Grace and Bunney, 1984a). The ionic flux concentrations, which drive membrane potential, were in the range of values used in previous models (Francis et al., 2013; Oster and Gutkin, 2011). The intracellular calcium concentration during resting state was ~1*x*10^−4^ *mM*, which can rise to values greater than 1*x*10^−3^ *mM* upon arrival of the action potential (Figure 8B(ii)) (Ben-Jonathan and Hnasko, 2001; Dedman and Kaetzel, 1997; Wojda et al., 2008). The calcium concentration in the ER was ~1000 times higher than in the cytoplasm (Figure 8B(iii)) (Wojda et al., 2008). In general, the calcium concentration in the MT will be lesser than the cytoplasm, but due to the higher mitochondrial density (Pacelli et al., 2015) and higher calcium loading in the SNc cells (Sulzer, 2007; Surmeier et al., 2010), the SNc mitochondrial calcium concentration was much higher than other cells (Figure 8B(iv)). Accompanying slow calcium buffering mechanisms, calcium-binding proteins such as calbindin and calmodulin act as rapid calcium buffering mechanisms (mobile calcium buffers) (Schwaller, 2010), which are present near calcium hotspots and bind rapidly to excess cytoplasmic calcium (Figure 8B(v, vi)).

### Characteristic Dopamine Dynamics of the SNc Neuron

The link between membrane potential, which was driven by the exchange of ionic concentrations and extracellular release of dopamine, which was driven by that membrane potential was described in (Tello-Bravo, 2012) (Figure 8C). The extracellular dopamine was ~ 45%K)^−6^ *mM* which was in the range of (34 – 48) *x*10^−6^ *mM* (Garris et al., 1997) (Figure 8C(iv)) for a number of vesicles in the readily releasable pool (*nRRP* = 10). The amount of extracellular dopamine concentration after the quantal release was dependent on the nRRP parameter (Figure 9B). The cytoplasmic dopamine concentration was ~12×10^−4^ *mM* which was in the range of 10^−4^ *to* 10^−3^ *mM* (Liu and Edwards, 1997) (Figure 8C(ii)). The vesicular dopamine concentration was ~500 *mM* which was 10^3^ – 10^5^ greater than cytoplasmic dopamine concentration (Ben-Jonathan and Hnasko, 2001).

### Characteristic Energy Metabolite Dynamics of the SNc Neuron

Active pumps and exchangers maintained the ionic equilibrium across the cell membrane where ATP drives the sodium-potassium and calcium pumps. Utilizing glucose and oxygen, ATP was produced in the cell through stages of processes such as glycolysis and oxidative phosphorylation (Figure 8D). The average basal ATP concentration in the SNc cell was ~ 2.4 *mM*, which was in the range of 2 – 4 *mM* (Burnstock, 2007) (Figure 8D(vi)). Other intermediate metabolites in the energy metabolism were in the range similar to (Cloutier and Wellstead, 2010, 2012) (Figure 8D).

### Energy Consumption by Different Cellular Processes of the SNc Neuron

The energy consumption in the SNc neuron by different cellular processes namely action potential propagation, vesicle recycling, dopamine packing, ER calcium sequestration, and protein degradation was estimated using the proposed model (Figure 9A). The peak instantaneous ATP consumption for action potential propagation and synaptic transmission (vesicle recycling and dopamine packing) were ~ 2.42*x*10^−4^ *mM* and ~ 8.16%10^−3^ *mM*. The ratio of ATP consumption for action potential propagation to the synaptic transmission was 1:3 which was similar to (Sengupta et al., 2013).

### Model Responses to Electrical Stimulation

In order to study the effect of increased electrical stimulation on firing frequency and dopamine release, electrical stimulation was carried on the proposed SNc neuronal model. Upon electrical stimulation (pulse width = 10 *msec*, frequency = 20 *Hz* and duration = 1 *sec*) with varying amplitude of stimulation from 50 *pA* to 300 *pA* with similar step size to (Dufour et al., 2014), there was not much change in the firing frequency till 130 *pA* but increased linearly with increasing stimulation amplitude from 150 *pA* onwards (Figure 10A). Upon electrical stimulation, there was a sharp increase in consumed ATP by ionic pumps at 150 *pA* (Figure 10C, blue trace) clearly correlating with increased firing frequency (Figure 10A). There was not much change in the consumed ATP by the whole other cellular processes till 130 *pA* but starts to increase with the increase in stimulation amplitude from 150 *pA* onwards (Figure 10C, orange trace) correlating with increased firing frequency (Figure 10A).

Upon electrical stimulation (pulse width = 10 msec, amplitude = 144 *pA* and duration = 2 *sec*) with varying frequency of stimulation from 10 *Hz* to 50 *Hz* with similar step size to (Wightman and Zimmerman, 1990), there was an increase in peak dopamine concentration with increased frequency of stimulation (Figure 10B, orange trace) similar to (Wightman and Zimmerman, 1990) (Figure 10B, blue trace). The consumed ATP by ionic pumps and whole cellular processes increased with increased frequency of stimulation (Figure 10D).

### Model Responses to Chemical Stimulation

In order to study the effect of glutamate application on the different properties such as firing frequency, energy consumption, and apoptotic signal, chemical stimulation was carried on the proposed SNc neuronal model. Upon chemical application (duration of stimulation (1 *sec*)) with varying glutamate concentration from 0.03 *mM* to 0.1 *mM* (see Supplementary material-4), there was a greater increase in the firing frequency in the presence of both AMPA and NMDA receptors than AMPA receptor alone (Figure 11A). Similar trend was observed in the ATP consumption by ionic pumps and whole cellular processes, it was higher for both AMPA and NMDA receptors than AMPA receptor alone (Figure 11B, 4D). The apoptosis occurs at lower concentration of glutamate in the SNc neurons with both AMPA and NMDA receptors as opposed to neurons with AMPA receptors alone (Figure 11C) (Surmeier et al., 2010; Zhou et al., 2013).

### Hypoglycemia and Hypoxia Conditions

By introducing energy deficiency in the form of hypoxia and hypoglycemia, we now studied the effect of hypoglycemia and hypoxia on the various critical molecular players in the SNc neuron. The energy deficiency conditions were implemented by varying glucose and oxygen levels in the proposed comprehensive model of SNc. The firing frequency of the model decreases (Figure 12A), and the firing pattern changes from spiking to bursting (Figure 12B) under severe hypoglycemia (low glucose) and hypoxia (low oxygen) conditions. The average cytoplasmic calcium concentration was higher which might be due to the reduced outward flux of calcium by active calcium pump and sodium-calcium exchangers as a result of lesser ATP availability at higher extent of hypoglycemia and hypoxia conditions (Figure 12C). The average ER and mitochondrial calcium concentrations were low which might be due to reduced sequestration of calcium into ER and MT, which in turn happens due to lesser ATP availability under more severe hypoglycemia and hypoxia conditions (Figure 12D, 12E).

The average cytoplasmic DA concentration was higher, which might be due to reduced DA packing into the vesicles as a result of lesser ATP availability under more severe hypoglycemia and hypoxia conditions (Figure 13A). The average extracellular and vesicular DA concentrations were low which might be due to reduced readily releasable vesicle pool as a result of lesser ATP availability which might affect the DA packing into the vesicles under more severe hypoglycemia and hypoxia conditions (Figure 13B, 13C).

The average fructose-6-phosphate (F6P) concentration was more affected by reduced glucose than reduced oxygen, and F6P concentration becomes very low for glucose concentration reduced beyond 4*x*10^−2^ *mM* (Figure 13D). The average fructose-2,6-phosphate (F26P) accumulation was higher during high glucose and low oxygen, which was an integrator of metabolic stress (Cloutier and Wellstead, 2010) (Figure 13E). The average glyceraldehyde-3-phosphate (GAP), average pyruvate (PYR) and average lactate (LAC) concentrations were higher during high glucose and low oxygen due to GAP and PYR being the intermediate metabolites in the glycolytic pathway and LAC being the by-product of anaerobic respiration (in the absence of oxygen) (Figure 13F, 13G, 13H). The average ATP concentration under normal condition was ~ 2.4 *mM* which was in the range of 2-4 *mM*(Burnstock, 2007) and ATP concentration becomes significantly low for glucose concentration reduced beyond 4*x*10^−2^ *mM* (Figure 13I). At low glucose and low oxygen, ATP level reaches a point where SNc neuron might adapt and starts bursting (Figure 12A) to transmit maximum information with minimal usage of energy (Balduzzi and Tononi, 2013; Sandhu et al., 2015) (Figure 13I). At low glucose (< 5%10^−2^ *mM*) and very low oxygen (< 0.2) (relative units) levels, the SNc neuron undergoes degeneration (Figure 13).

In the whole (fast dynamics) model simulation, the healthy alpha-synuclein protein (asyn) was misfolded, and the available healthy alpha-synuclein protein was low at low glucose and low oxygen (Figure 14A, 14E). Under low glucose and low oxygen conditions, the accumulation of misfolded alpha-synuclein (asyn_mis_) and alpha-synuclein aggregates (asyn_agg_) was higher due to lesser ATP availability which leads to reduced proteolysis or protein degradation (Figure 14B, 14C). The average ROS concentration was increased at low glucose and low oxygen levels due to misfolded alpha-synuclein, thereby inducing further release of ROS by hindering mitochondrial functioning (Figure 14D). For a better representation of molecular markers under pathological conditions, the reduced (slow dynamics) model was simulated which was obtained by assuming fast substrates reaching their steady states rapidly, and associated differential equations were transformed into functions (that is, at steady state values). The average normal alpha-synuclein concentration decreases with a decrease in glucose and oxygen levels due to increased ROS-induced misfolding of alpha-synuclein (Figure 14E). The deleterious effect of ROS/α-syn_mis_ leads to a vicious cycle where the formation of ROS and α-syn_mis_ was facilitated by each other (Cloutier and Wellstead, 2012), which was evident from simulation results also. The average ROS concentration during normal condition was in the range of 1*x*10^−3^ – 5×10^−3^ *mM* and during hypoglycemia and hypoxia conditions it reached beyond the concentrations (0.01–0.015 *mM*) (Desagher et al., 1996) which was observed in the disease state (Figure 14H). Due to higher ROS concentration, alpha-synuclein misfolding and aggregation were prominent, and the concentrations are reaching values similar to high-stress conditions (Cloutier and Wellstead, 2012) (Figure 14F, 14G).

## DISCUSSION

The central objective of this computational study is to show that metabolic deficiency is the root cause that connects various molecular level pathological manifestations of PD in SNc cells. More importantly, we want to investigate the hypothesis that metabolic deficit is perhaps the root cause of SNc cell loss in PD. The proposed model is one of its kind which explains how deficits in supply of energy substrates (glucose and oxygen) can lead to the pathological molecular changes including alpha-synuclein aggregation, ROS production, calcium elevation, and dopamine deficiency. The proposed model is compared to other models, that at least had more than one cellular process modeled together (Table-1).

**Table-1:** Comparison of the proposed model with previously published models.

| Model | Ion channels | Calcium buffering | Energy metabolism | Dopamine turnover | Levodopa uptake | ROS/α-syn | Apoptosis |
| --- | --- | --- | --- | --- | --- | --- | --- |
| (Tello-Bravo, 2012) | ✓ | ✗ | ✗ | ✓ | ✗ | ✗ | ✗ |
| (Reed et al., 2012) | ✗ | ✗ | ✗ | ✓ | ✓ | ✗ | ✗ |
| (Cloutier and Wellstead, 2012) | ✗ | ✗ | ✓ | ✗ | ✗ | ✓ | ✗ |
| (Francis et al., 2013) | ✓ | ✓ | ✗ | ✗ | ✗ | ✗ | ✗ |
| (Cullen and Wong-Lin, 2015) | ★ | ✗ | ✗ | ✓ | ✗ | ✗ | ✗ |
| **Proposed model** | ✓ | ✓ | ✓ | ✓ | ✓ | ✓ | ✓ |

★ No ion channel dynamics but has spiking behavior (Izhikevich neuronal model (Izhikevich, 2003)).

### Different Regimes with Varying Energy Substrates

The proposed model with its biophysical framework shows four regimes of ATP dynamics by varying glucose and oxygen levels: (A) Unperturbed (no change in Basal ATP Concentration (BAC)), (B) adaptation (initial drop and a subsequent return to initial BAC) (Connolly et al., 2014), (C) no adaptation (initial drop and stabilized at a lower BAC, however, generally astrocytes and other energy sources (glycogen, glutamine) will restore ATP levels (Amaral et al., 2011)), and (D) oscillating (BAC fluctuates, where anaerobic respiration might occur) (Fadaka et al., 2017) and other regimes in which neuron undergoes degeneration (Figure 15A). The model also suggests that hypoglycemia plays a more crucial role in leading to ATP deficits than hypoxia (Figure 15B). From the modelling results, the relative levels of ATP consumption in different cellular processes can be described as: synaptic transmission > action potential propagation > endoplasmic reticulum calcium sequestration > protein degradation (Attwell and Laughlin, 2001; Harris et al., 2012).

**Figure 15:**
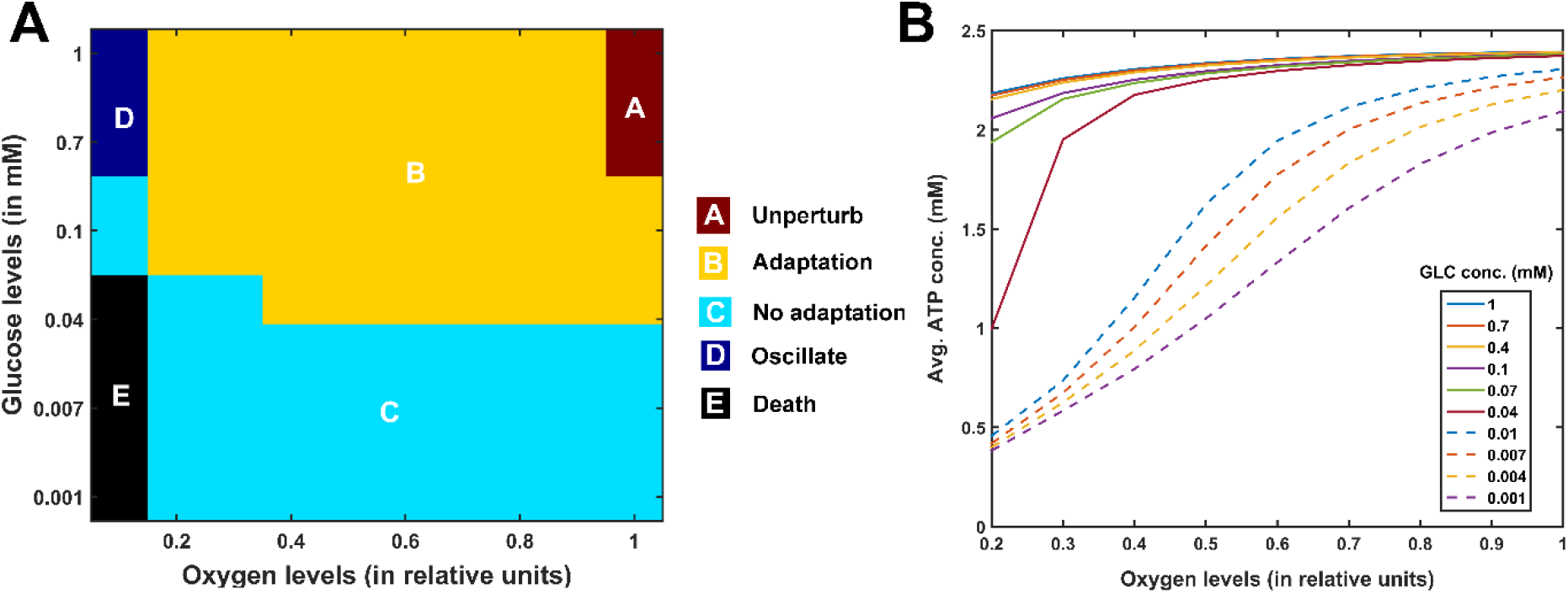
Model responses to hypoglycemia and hypoxia conditions. (A) Different regimes of the model response to hypoglycemia and hypoxia conditions, (B) Average ATP concentration for different initial glucose concentration concerning oxygen concentration. conc, concentration; mM, millimolar; GLC, glucose; ATP, adenosine triphosphate.

### Excitotoxicity Precipitated by Energy Deficiency

During chemical stimulation or synaptic evoked action potential, glutamate concentration varies from 0.03 *mM* to 0.1 *mM* which was in the range observed in the synaptic cleft (2*x*10^−3^ – 1 *mM*) and the binding affinities of NMDA (2*x*10^−3^ – 3*x*10^−3^ *mM*) and AMPA (0.4 – 0.5 *mM*) receptors (Meldrum, 2000). From the proposed model, the SNc neurons with both AMPA and NMDA receptors are more prone to apoptosis than SNc neurons with AMPA receptor alone (Surmeier et al., 2010; Zhou et al., 2013) (Figure 11C). Thus, the long-term influence of NMDA activation (longer time constant than that of AMPA) in the SNc neuron plays an important role in PD pathogenesis (Hallett and Standaert, 2004; Loopuijt and Schmidt, 1998; Surmeier et al., 2010). Under energy deficit conditions, SNc neurons undergo apoptosis due to overexcitation with even physiological concentrations of glutamate when compared to normal conditions (not shown here) (Connolly et al., 2014). We suggest that the excitotoxic loss of SNc neurons in PD might be precipitated by energy deficiency (Muddapu et al., 2019). Any therapeutic interventions that can reduce ionic flux through these glutamatergic receptors or enhance energy production can be neuroprotective in nature (Bathina and Das, 2015; Maiolino et al., 2019; Wallace et al., 2010).

### Insights into Different Phenotypes of PD (Determinants at Different Levels)

In genetics, the phenotype of an organism depends on the underlying genotype (Talbot et al., 2016). Similarly, the occurrence of different phenotypes of a disease can be driven by underlying dysfunctions occurring at different levels in the hierarchy such as molecular, cellular and systems levels (Angeli et al., 2013; Zuo et al., 2017). In PD, the loss of dopaminergic neurons in SNc results in the manifestation of PD symptoms and the cause of the SNc cell loss is still not clearly elucidated. The PD phenotypes are distinct, and this specificity might be arising out of a combination of interactions between key determinants at the same or different levels.

At the molecular level, the interactions among divergent key determinants such as adenosine triphosphate (ATP), cytoplasmic dopamine (DA_cyt_), alpha-synuclein (ASYN), reactive oxygen species (ROS) and cytoplasmic calcium (Ca^2+^) converges to common pathologies or pathways such as oxidative stress, mitochondrial impairment and protein mishandling (Greenamyre and Hastings, 2004; Levy et al., 2009; Post et al., 2018). The dysfunction causing interactions among different molecular determinants was elaborated in Figure 16 (Betzer et al., 2018; Brookes et al., 2004; Post et al., 2018).

**Figure 16:**
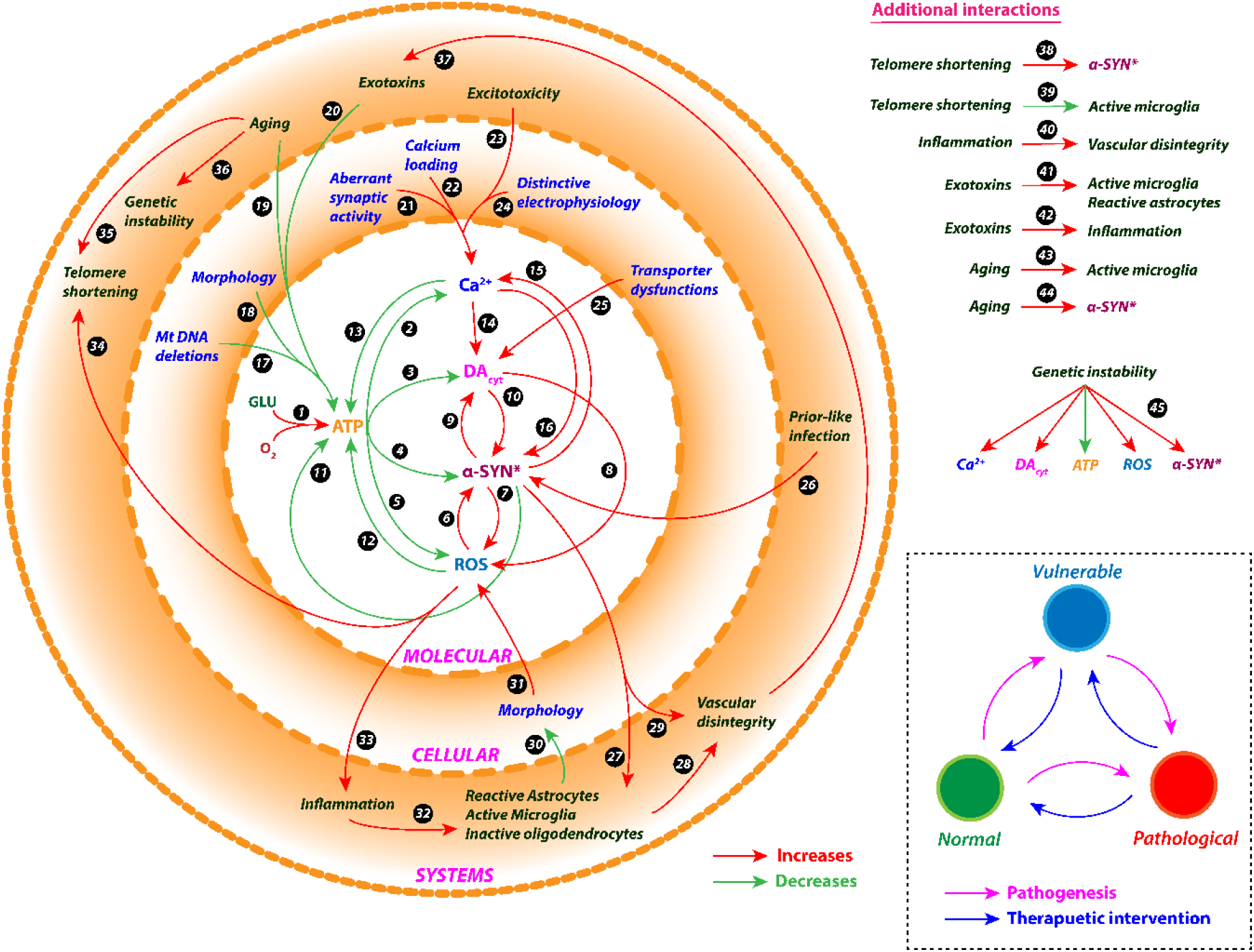
Interactions among the determinants at different levels of hierarchy. See Box-1 for description of the Figure.

At the cellular level, the determinants that might contribute to differential PD phenotypes are complex morphology (due to large axonal arborization and numerous synaptic connectivity) (Bolam and Pissadaki, 2012; Pacelli et al., 2015; Pissadaki and Bolam, 2013), lesser mitochondrial mass (due to higher level of mitochondrial DNA deletions) (Kraytsberg et al., 2006; Liang et al., 2007), high levels of reactive cytosolic dopamine (due to underexpression of vesicular monoamine transporter 2 and overexpression of dopamine transporter) (Anderegg et al., 2015; Brichta and Greengard, 2014; Liang et al., 2004; Peter et al., 1995), distinctive electrophysiology (due to broad spikes and pacemaking activity) (Bean, 2007; Chan et al., 2007; Guzman et al., 2009), calcium loading (due to presence of Cav1.3 calcium channels and low calcium buffering) (Anderegg et al., 2015; Foehring et al., 2009; Philippart et al., 2016) and aberrant excitatory synaptic activity (due to ineffective magnesium blockage of NMDA receptors and increased NMDA receptor subunit NR1) (Roselli and Caroni, 2015; Surmeier et al., 2010). These cellular determinants individually or collectively would result in higher basal metabolic rate and increased oxidative stress (Pacelli et al., 2015) which in turn converges to common pathologies (Duda et al., 2016) (Figure 16).

At the systems level, the determinants that might contribute to differential PD phenotypes are excitotoxicity (due to overexcitation by subthalamic nucleus or pedunculopontine nucleus) (Pahapill and Lozano, 2000; Rodriguez et al., 1998), aging (due to proteostatic dysfunction, mitochondrial dysfunction, genetic mutations or telomere shortening) (Birch et al., 2018; Surmeier, 2018), genetic instability (due to changes in nucleic acid sequences, chromosomal rearrangements or aneuploidy) (Mullin and Schapira, 2015; Selvaraj and Piramanayagam, 2019; Singleton et al., 2013), environmental toxins (due to exposure to insecticides, commercial solvents, metal exposure or traumatic head injury) (Goldman, 2014; Nandipati et al., 2016), neuroinflammation (due to traumatic head injury, exotoxins or immune dysfunctions) (Caggiu et al., 2019; Gardner et al., 2018), prion-like infection (bacteria or viruses) (Brugger et al., 2015; Caggiu et al., 2019), telomere shortening (due to aging or oxidative stress) (Kolyada et al., 2016; Scheffold et al., 2016), glial dysfunctions (due to phagocytic or inflammatory impairments, enteric glial dysfunction) (Clairembault et al., 2015; di Domenico et al., 2019; Lecours et al., 2018) and vascular dysfunctions (due to endothelial dysfunction or cardiovascular autonomic dysfunction) (Kim et al., 2017; Yang et al., 2017). These systems-level determinants interact among themselves and also across different levels in the hierarchy resulting in different PD phenotypes (Figure 16).

#### Box-1: Description of the Figure 16.

***(1)** ATP production by aerobic glucose metabolism, **(2)** Ca^2+^ efflux by ATP-dependent calcium pump, **(3)** DA_cyt_ packing into vesicles by VMAT using H^+^-ATPase-induced concentration gradient, **(4)** ATP-dependent protein degradation by UPS and autophagy, **(5)** ROS scavenging mechanism by glutathione, **(6)** α-syn* aggregation due to ROS-induced UPS impairment, **(7)** ROS formation due to α-syn* induced mitochondrial dysfunction, **(8)** ROS formation due to DA_cyt_ autoxidation, **(9)** DA_cyt_ accumulation due to α-syn* induced vesicle recycling impairment, **(10)**α-syn* aggregation due to DA_cyt_ induced CMA impairment, **(11)** Reduced ATP production due to α-syn* induced mitochondrial dysfunction, **(12)** Reduced ATP production due to ROS-induced mitochondrial dysfunction, **(13)** Reduced ATP production due to Ca^2+^ induced mitochondrial dysfunction, **(14)** DA_cyt_ accumulation due to Ca^2+^ induced DA synthesis, **(15)** Ca^2+^ accumulation due to α-syn* induced dysregulation of Ca^2+^homoestasis, **(16)** α-syn* aggregation due to Ca^2+^ induced calpain activation, **(17)** Reduced ATP production due to mitochondrial DNA deletions, **(18)** Reduced ATP production due to ROS formation induced by complex anatomical structures, **(19)** Reduced ATP production due to age-dependent hypoglucometabolism, **(20)** Reduced ATP production due to exotoxins-induced mitochondrial impairments, **(21)** Ca^2+^ accumulation due to aberrant excitatory synaptic activity, **(22)** Ca^2+^ accumulation due to poor calcium buffering mechanisms, **(23)** Ca^2+^accumulation due to distinctive electrophysiology, **(24)** Ca^2+^ accumulation due to overexcitation through afferent connections, **(25)** DA_cyt_ accumulation due to dysfunction of VMAT and DAT, **(26)** α-syn* aggregation due to prior-like infection, **(27)** α-syn* induced dysfunction of glial cells, **(28)**Vascular disintegration due to inflammatory response by glial cells, **(29)** Vascular disintegration due to α-syn*, **(30)** Disruption of anatomical structures due to inflammatory response by glial cells, **(31)** ROS formation induced by complex anatomical structures, **(32)** Inflammatory signals activates microglia and convert astrocytes into reactive astrocytes, **(33)** Inflammatory response due to ROS **(34)** ROS-induced shortening of telomere **(35)** Age-dependent shortening of telomere **(36)** Age-dependent instability in genome **(37)** Exotoxins accumulation due to vascular disintegrity **(38)** α-syn* aggregation due to telomere shortening **(39)** Microglia impairment due to telomere shortening **(40)** Vascular disintegration due to inflammation response **(41)**Exotoxins-induced activation of microglia **(42)** Exotoxins-induced inflammatory response **(43)** Age-dependent activation of microglia **(44)** Age-dependent α-syn* aggregation **(45)** Genetic instabilities affects Ca^2+^ homeostasis, DA_cyt_ accumulation, α-syn* aggregation, ROS formation and ATP production. **Inset:** Proposed states of SNc neuron. GLU, glucose; O_2_, oxygen; ATP, adenosine triphosphate; Ca^2+^, calcium; DA_cyt_, cytoplasmic dopamine; α-syn*, alpha-synuclein aggregates; ROS, reactive oxygen species; DA, dopamine; VMAT, vesicular monoamine transporter; DAT, dopamine transporter; UPS, ubiquitin-proteasome system; CMA, chaperone-mediated autophagy; DNA, deoxyribonucleic acid*.

Dysfunctions at any level of hierarchy would make SNc cell move from normal state to pathological state directly or indirectly via intermediate (vulnerable) state (Figure 16, inset). Any therapeutics that can bring back SNc neuron from pathological or vulnerable state to normal state can be beneficiary for the survival of SNc neurons.

### Potential Experimental Setup to Validate Predictions from the Proposed Model

Here, we suggest some experimental approaches to evaluate the behavior of dopaminergic neuron at single-cell or network level by capturing the dynamics of critical molecular players in different conditions. During energy-deficient conditions, monitoring important intracellular players such as ATP, glucose, AMP-activated protein kinase (AMPK), and lactate using single-cell imaging studies gives an insight into the progressive adaptation of dopaminergic neurons to the energy crisis by activating various compensatory mechanisms (Connolly et al., 2014; Connolly and Prehn, 2015). Also, we can determine all the cellular processes that are compromised during energy crisis. Mitochondria play a major role in maintaining cellular energy levels (Osellame et al., 2012) and monitoring its functioning capacity provides insights into cellular energy production. Using cellular models (Connolly et al., 2018), monitoring the mitochondrial calcium, ATP, NADPH, pH, membrane potential, oxygen consumption rate, ROS production and morphology gives better understanding of mitochondrial bioenergetic function in the neuron under energy deficits, oxidative stress and excitotoxicity (Connolly et al., 2018; Ludtmann et al., 2018; Pacelli et al., 2015; Theurey et al., 2019). During progressive energy deficiency, dopamine and its metabolites can be measured to check for production of ROS leading to oxidative stress in the neuron using toxin-induced animal pathological models (Puginier et al., 2019).

### Future Directions

In the proposed model, ketone metabolism (Morris, 2005) can be incorporated to make the model more robust to utilize different substrates as an energy source and understand the role of ketone bodies in PD pathogenesis (Phillips et al., 2018; Włodarek, 2019). Apart from ketone bodies, astrocytes also play an important role in maintaining neuronal energy demands (Jha and Morrison, 2018). Therefore, combining the SNc neuronal model with astrocyte model will provide a better understanding of compensation due to astrocyte involvement in energy deficit conditions (Kuter et al., 2019). The ischemic condition was implemented by modulating glucose and oxygen levels, which can be extended by introducing the vascular module (Cloutier et al., 2009), where ischemia condition can be simulated more realistically by varying cerebral blood flow. Cancer cells survive in low oxygen and acidic conditions (Weisz, 2015) where pH plays a vital role in the functioning of cellular processes (Putnam, 2012) thus considering potentiometric properties in formulating cellular processes could be more biologically realistic (pH plays an essential role in mitochondrial functioning).

## CONCLUSIONS

In conclusion, we believe that the proposed model provides an integrated modelling framework to understand the neurodegenerative processes underlying PD (Lloret-Villas et al., 2017). From the simulation results, it was observed that, under conditions of energy starvation, intracellular calcium, dopamine (cytoplasmic), alpha-synuclein, and ROS concentrations significantly deviated from normal values (equilibrium). There is a positive feedback loop formed with increased intracellular calcium, or dopamine levels lead to oligomerization of alpha-synuclein, while alpha-synuclein oligomers increased intracellular calcium and dopamine levels (Post et al., 2018). Any therapeutics that can reduce these key toxicity mediators can be beneficiary for the survival of SNc neurons (Catoni et al., 2019; Mosharov et al., 2009; Post et al., 2018). The big picture of developing such model was to develop a therapeutic computational testbed for PD wherein the proposed model of SNc will be the center of a larger framework, which will also be integrated to behavioural model (Muralidharan et al., 2018). This type of framework will help in providing personalized medicine for PD patients (Bloem et al., 2019) rather than the currently employed trial and error approaches.

## CODE ACCESSIBILITY

The comprehensive SNc model code is available in ModelDB database (McDougal et al., 2017).

## AUTHOR CONTRIBUTIONS

VRM and VSC: conceived, developed the model, and prepared the manuscript.

