## Supplementary material-1 for "Influence of Energy Deficiency on the Molecular Processes of *Substantia Nigra Pars Compacta* Cell for Understanding Parkinsonian Neurodegeneration: A Comprehensive Biophysical Computational Model"

**Table-1:** Published dopaminergic neuronal models.

| S.No. | Model | Ion channels | Pumps and exchangers (Ionic balance) | Synaptic currents | Reference(s) |
|---|---|---|---|---|---|
| 1. | Two-compartment – soma and dendrite | Soma: $I_{K,DR}$, $I_{Na}$<br>Dendrite: $I_L$ | Dendrite: $I_{NaKP}$<br>(sodium) | Dendrite: $I_{NMDA}$ | (Li et al., 1996) |
| 2. | Single-compartment soma with calcium buffering (CBP) | Soma: $I_{Ca,T}$, $I_{Ca,L}$, $I_{Ca,N}$, $I_{Ca,HVA}$, $I_{K,Ca}$, $I_{K,DR}$, $I_{K,A}$, $I_H$, $I_B$ | Soma: $I_{NaKP}$, $I_{CaP}$, $I_{NaCaX}$<br>(calcium) | - | (Amini et al., 1999) |
| 3. | Three compartments – Soma, proximal and distal dendrites | $I_{K,DR}$, $I_L$ | $I_{NaKP}$<br>(sodium in all) | Distal dendrite: $I_{NMDA}$, $I_{AMPA}$, $I_{GABAA}$ | (Canavier, 1999) |
| 4. | (Amini et al., 1999) model with calcium diffusion (also abstract version) | Soma: $I_{Ca}$, $I_{K,Ca}$, $I_K$, $I_L$ | Soma: (calcium) | - | (Medvedev et al., 2003; Medvedev and Kopell, 2001; Wilson and Callaway, 2000) |
| 5. | Two (Canavier, 1999) models coupled at distal dendrites | $I_{Na}$, $I_{K,DR}$, $I_L$, $I_{K,A}$ | $I_{NaKP}$ | Distal dendrite: $I_{NMDA}$ | (Komendantov and Canavier, 2002) |
| 6. | Soma with four identical branched dendrites with a single proximal and two distal branches | Soma: $I_{Na}$, $I_{K,A}$, $I_{K,DR}$, $I_L$, $I_{K,Ca}$, $I_{Ca,T}$, $I_{Ca,L}$, $I_{Ca,N}$<br>Dendrite: $I_{Na}$, $I_{K,A}$, $I_{K,DR}$, $I_L$ | Soma: $I_{NaKP}$, $I_{CaP}$<br>(calcium)<br>Dendrite: $I_{NaKP}$ | Soma: $I_{GABAA}$<br>Dendrite: $I_{NMDA}$, $I_{GABAA}$ | (Komendantov et al., 2004) |
| 7. | Modified (Komendantov | Soma: $I_{Na}$, $I_{K,A}$, $I_{K,DR}$, | Soma: $I_{NaKP}$, $I_{CaP}$ | Soma: $I_{GABAA}$ | (Canavier and Landry, 2006) |

| | | | | | |
|---|---|---|---|---|---|
| | et al., 2004) model with $I_{AMPA}$ synaptic current in dendrite | $I_{K,Ca}$, $I_L$, $I_{Ca,T}$, $I_{Ca,L}$, $I_{Ca,N}$<br><br>Dendrite: $I_{Na}$, $I_{K,A}$, $I_{K,DR}$, $I_L$ | (calcium)<br><br>Dendrite: $I_{NaKP}$<br><br>(sodium) | Dendrite: $I_{NMDA}$, $I_{AMPA}$, $I_{GABAA}$ | |
| 8. | Modified (Wilson and Callaway, 2000) model with $I_{AMPA}$ and $I_{NMDA}$ synaptic currents along with spiking generating ion channels | Soma:$I_{Ca}$, $I_{K,Ca}$, $I_K$, $I_L$, $I_{Na}$, $I_{K,DR}$ | Soma: (calcium) | Soma: $I_{NMDA}$, $I_{AMPA}$ | (Kuznetsov et al., 2006) |
| 9. | Single-compartment soma | Soma: $I_{Ca,L}$, $I_{Ca,B}$, $I_{K,ERG}$, $I_{K,Ca}$, $I_H$, $I_L$ | Soma: $I_{CaP}$<br><br>(calcium) | - | (Canavier et al., 2007) |
| 10. | Modified (Komendantov et al., 2004) model with pacemaking mechanism throughout soma and dendrites | Soma: $I_{Na}$, $I_A$, $I_{K,DR}$, $I_L$, $I_{K,Ca}$, $I_{Ca,L}$<br><br>Dendrite: $I_{Na}$, $I_A$, $I_{K,DR}$, $I_L$, $I_{K,Ca}$, $I_{Ca,L}$ | Soma: (calcium) | - | (Kuznetsova et al., 2010; Yu et al., 2014) |
| 11. | Single-compartment soma | Soma: $I_{Ca,L}$, $I_{Na}$, $I_{K,DR}$, $I_{K,Ca}$, $I_L$ | Soma: $I_{CaP}$<br><br>(calcium) | - | (Drion et al., 2011) |
| 12. | Single-compartment soma which is combines conductance mechanisms from (Amini et al., 1999) and (Kuznetsov et al., 2006) | Soma: $I_{Ca,L}$, $I_{Na}$, $I_{K,DR}$, $I_{K,Ca}$, $I_L$, $I_K$ | Soma: $I_{CaP}$<br><br>(calcium) | Soma: $I_{NMDA}$, $I_{GABAA}$ | (Oster and Gutkin, 2011) |
| 13. | Single-compartment | Soma: $I_{Ca,L}$, $I_{Na}$, $I_{Na,HCN}$, | Soma: $I_{NaKP}$, $I_{CaP}$, $I_{NaCaX}$ | - | (Francis et al., 2013) |

| | | | | | |
|---|---|---|---|---|---|
| | soma with calcium buffering (CBP) | $I_{L,Na}$, $I_{K,DR}$, $I_{L,IR}$, $I_{K,Ca}$ | (calcium, sodium, potassium, calbindin, calmodulin) | | |
| 14. | Modified (Kuznetsov et al., 2006) model with altered NMDA and $I_{K,ERG}$ along with full morphology of dendrite (reduced model) | Soma: $I_{Ca,L}$, $I_{Na}$, $I_{K,DR}$, $I_{K,Ca}$, $I_L$, $I_{K,ERG}$ | Soma: $I_{CaP}$ (calcium) | Soma: $I_{NMDA}$, $I_{AMPA}$ | (Ha and Kuznetsov, 2013; Zakharov et al., 2016) |
| 15. | Single-compartment soma | Soma: $I_{Na}$, $I_{K,DR}$, $I_L$, | - | Soma: $I_{NMDA}$, $I_{AMPA}$ | (Qian et al., 2014) |
| 15. | Single-compartment soma with full morphology of dendrite | Soma: $I_{Ca}$, $I_{Na}$, $I_{K,DR}$, $I_{K,Ca}$, $I_{L,Ca}$, $I_{K,ERG}$, $I_H$, $I_L$ | Soma: $I_{CaP}$ (calcium) | - | (Yu and Canavier, 2015) |
| 16. | Simple (spiking) dopaminergic neuronal model | Izhikevich (point neuron) – two variable neuronal model | - | - | (Cullen and Wong-Lin, 2015; Muddapu et al., 2019) |
| 17. | Modified (Ha and Kuznetsov, 2013) | Soma: $I_{Ca}$, $I_{Na}$, $I_{Na,S}$, $I_{K,DR}$, $I_{K,Ca}$, $I_L$, $I_K$, $I_H$ | Soma: $I_{CaP}$ (calcium) | Soma: $I_{NMDA}$, $I_{AMPA}$, $I_{GABAA}$ | (Morozova et al., 2016b, 2016a) |
| 18. | Single-compartment soma with calcium buffering (CBP) along $I_{K,ATP}$ mediated bursting | Soma: $I_{Ca,L}$, $I_{Na}$, $I_{K,DR}$, $I_{K,ATP}$, $I_{L,Ca}$, $I_L$ | Soma: $I_{CaP}$ (calcium) | Soma: $I_{NMDA}$ | (Knowlton et al., 2018) |

| | | | | | |
|---|---|---|---|---|---|
| 19. | Modified (Kuznetsova et al., 2010) model | $I_{Ca,L}$, $I_{Ca,T}$, $I_{Na}$, $I_{Na,HCN}$, $I_{K,DR}$, $I_{K,B}$, $I_{K,Ca}$, $I_{K,A}$, $I_{K,ERG}$, $I_L$ | Soma: $I_{CaP}$ (calcium) | - | (Rumbell and Kozloski, 2019) |

$I_{Ca,T}$ – T-type calcium current; $I_{Ca,L}$ – L-type calcium current; $I_{Ca,N}$ – N-type calcium current; $I_{Ca,HVA}$ – residual high-voltage activated calcium current; $I_{Ca}$ – calcium current; $I_{K,Ca}$ – calcium-activated (small conductance) potassium current; $I_{K,DR}$ – delayed rectifier potassium current; $I_{K,A}$ –transient outward (4-aminopyridine-sensitive) potassium current; $I_H$ – hyperpolarization-activated cation current; $I_B$ – background current (sodium, potassium, calcium); $I_{NaKP}$ – sodium-potassium pump; $I_{CaP}$ – calcium pump; $I_{NaCaX}$ – sodium-calcium exchanger; $I_L$ – leaky current; $I_{Na}$ – fast spiking (tetrodotoxin-sensitive) sodium current; $I_{NMDA}$ – N-methyl-D-aspartic acid (NMDA) current; $I_{AMPA}$ – alpha-amino-3-hydroxy-5-methyl-4-isoxazolepropionic acid (AMPA) current; $I_{GABAA}$ – gamma-aminobutyric acid A-class (GABAA) current; $I_{K,ERG}$ – ERG (ether-a-go-go-related gene) potassium current; $I_{Ca,B}$ – background calcium leak current; $I_{L,Ca}$ – leaky calcium current; CBP – calcium-binding proteins; $I_{L,Na}$ – leaky sodium current; $I_{K,IR}$ – inward rectifying potassium current; $I_{Na,HCN}$ – hyperpolarization-activated cyclic nucleotide (HCN) sodium current; $I_{Na,S}$ – subthreshold sodium current; $I_K$ – intrinsic potassium current; $I_{K,ATP}$ – ATP-sensitive potassium current; $I_{K,B}$ – large conductance potassium current;
