## Supplementary material-2 for "Influence of Energy Deficiency on the Molecular Processes of *Substantia Nigra Pars Compacta* Cell for Understanding Parkinsonian Neurodegeneration: A Comprehensive Biophysical Computational Model"

**Table-1:** Published dopaminergic terminal models.

| S.No. | Model | Metabolite balance (units) | Autoreceptors | Reference(s) |
|---|---|---|---|---|
| 1. | Two-compartment – cytoplasmic and extracellular | $DA_c$, $DA_v$, $DA_e$, $LDOPA$, $I1$, $I2$ <br> (nmol/g, min) | DA synthesis | (Porenta and Riederer, 1982) |
| 2. | Two-compartment – cytoplasmic and extracellular | $DA_i$, $DA_e$ <br> ($\mu$g/g, min) | DA firing | (King et al., 1984) |
| 3. | Three-compartment – cytoplasmic, vesicular and extracellular | $DA_c$, $DA_v$, $DA_e$, $DA_a$, $DA_g$, $3MT$, $LDOPA$, $DOPAC$, $HVA$ <br> (mM, ms) | - | (Justice et al., 1988) |
| 4. | Biochemical systems theory model | DA homeostasis <br> (relative units) | - | (Qi et al., 2008a, 2008b) |
| 5. | Three-compartment – cytoplasmic, vesicular and extracellular | $DA_c$, $DA_v$, $DA_e$, $TYR$, $LDOPA$, $BH_2$, $BH_4$, $HVA$, $TYRPOOL$ <br> ($\mu$M, hr) | DA synthesis | (Best et al., 2009; Reed et al., 2009) |
| 6. | Three-compartment – cytoplasmic, vesicular and extracellular | $DA_c$, $DA_v$, $DA_e$ <br> (mM, ms) | DA synthesis, DA release | (Tello-Bravo, 2012) |
| 7. | Modified (Best et al., 2009) model with DA and 5HT cell bodies and 5HT terminal | DA terminal: $DA_c$, $DA_v$, $DA_e$, $TYR$, $LDOPA$, $BH_2$, $BH_4$, $HVA$, $TYRPOOL$ | DA synthesis, DA firing, 5HT synthesis, 5HT firing | (Reed et al., 2012) |

| | | | | |
|---|---|---|---|---|
| | | 5HT terminal: $5HT_c$, $5HT_v$, $5HT_e$, $TRYP$, $5HTP$, $BH_2$, $BH_4$, $HIA$, $TRPPOOL$<br><br>($\mu$M, hr) | | |
| 8. | DA neurotransmission model | Volume transmission<br><br>($\mu$M, sec) | DA firing, DA release | (Dreyer et al., 2010; Dreyer and Hounsgaard, 2013) |
| 9. | Systems Biology Markup Language model | Flux balance analysis<br><br>($\mu$M, hr) | - | (Büchel et al., 2013) |
| 10. | Modified (Best et al., 2009) model with spiking neuronal model | DA terminal: $DA_c$, $DA_v$, $DA_e$, $TYR$, $LDOPA$, $BH_2$, $BH_4$, $HVA$, $TYRPOOL$<br><br>($\mu$M, hr) | DA synthesis, DA firing | (Cullen and Wong-Lin, 2015) |

$DA$ – dopamine; $5HT$ – serotonin; $DA_c$ – cytoplasmic DA; $DA_v$ – vesicular DA; $DA_e$ – extracellular DA; $DA_a$ – inactive DA; $DA_g$ – glial DA; $TH$ – tyrosine hydroxylase; $I1$ – competitive TH inhibitor 1; $I2$ – competitive TH inhibitor 2; $LDOPA$ – 3,4-dihydroxyphenylalanine; $3MT$ – 3-methoxytyramine; $DOPAC$ – 3,4-dihydroxyphenylacetic acid; $HVA$ – homovanillic acid; $TYR$ – tyrosine; $BH_2$ – dihydrobiopterin; $BH_4$ – tetrahydrobiopterin; $TYRPOOL$ – tyrosine pool; $5HT_c$ – cytoplasmic 5HT; $5HT_v$ – vesicular 5HT; $5HT_e$ – extracellular 5HT; $5HTP$ – 5-hydroxytryptophan; $HIA$ – 5-hydroxyindoleacetic acid; $TRYP$ – tryptophan; $TRYPPOOL$ – tryptophan pool; $\mu M$ – micromolar; $mM$ – millimolar; $ms$ – millisecond; $hr$ – hour; $DA_i$ – intracellular DA; nmol – nanomole; g – gram; min – minute.
