## Supplementary material-3 for "Influence of Energy Deficiency on the Molecular Processes of *Substantia Nigra Pars Compacta* Cell for Understanding Parkinsonian Neurodegeneration: A Comprehensive Biophysical Computational Model"

### Ion Channels

**Table-1:** Parameter values for ion-channel dynamics of SNc cell model (Francis et al., 2013).

| Constant | Symbol | Value | Units |
|---|---|---|---|
| Faraday's constant | $F$ | 96485 | $coulomb * mole^{-1}$ |
| SNc membrane capacitance | $C_{snc}$ | $9\ x\ 10^{7}$ | $pF * cm^{-2}$ |
| Cytosolic volume | $v_{cyt}$ | $\phi_{cyt} * v_{pmu}$ | $pl$ |
| Fraction of cytosolic volume | $\phi_{cyt}$ | 0.5 | $dimensionless$ |
| Pacemaking unit (PMU) volume | $v_{pmu}$ | 5 | $pl$ |
| PMU area | $\mathcal{A}_{pmu}$ | $\mathcal{S}_{pmu} * v_{pmu}$ | $cm^{2}$ |
| PMU surface area-to-volume ratio | $\mathcal{S}_{pmu}$ | $1.6667\ x\ 10^{4}$ | $cm^{-1}$ |
| Voltage defined thermodynamic entity | $V_D$ | $\frac{V}{V_\tau}$ | $dimensionless$ |
| Temperature defined thermodynamic entity | $V_\tau$ | $\frac{R * T}{F}$ | $mV$ |
| Universal gas constant | $R$ | 8314.472 | $mJ * mol^{-1} * K^{-1}$ |
| Physiological temperature | $T$ | 310.15 | $K$ |
| Maximal conductance of calcium channel | $\bar{g}_{Ca,L}$ | 2101.2 | $pA * mM^{-1}$ |
| Extracellular calcium concentration | $[Ca_e]$ | 1.8 | $mM$ |
| Reversal potential for calcium ion | $V_{Ca}$ | $\frac{1}{2} * \log\left(\frac{[Ca_e]}{[Ca_i]}\right)$ | $dimensionless$ |
| Valence of calcium ion | $z_{Ca}$ | 2 | $dimensionless$ |
| Maximal conductance of sodium channel | $\bar{g}_{Na}$ | 907.68 | $pA * mM^{-1}$ |
| Extracellular sodium concentration | $[Na_e]$ | 137 | $mM$ |
| Reversal potential for sodium ion | $V_{Na}$ | $\log\left(\frac{[Na_e]}{[Na_i]}\right)$ | $dimensionless$ |
| Valence of sodium ion | $z_{Na}$ | 1 | $dimensionless$ |

| | | | |
|---|---|---|---|
| Maximal conductance of sodium HCN channel | $\bar{g}_{NaHCN}$ | 51.1 | $pA * mM^{-1}$ |
| Maximal conductance of leaky sodium channel | $\bar{g}_{Nalk}$ | 0.0053 | $pA * mM^{-1}$ |
| Cyclic adenosine monophosphate concentration | $[cAMP]$ | $1\ x\ 10^{-5}$ | $mM$ |
| Maximal conductance of delayed rectifying potassium channel | $\bar{g}_{Kdr}$ | 31.237 | $nS$ |
| Extracellular potassium concentration | $[K_e]$ | 5.4 | $mM$ |
| Reversal potential for potassium ion | $V_K$ | $\log\left(\frac{[K_e]}{[K_i]}\right)$ | $dimensionless$ |
| Valence of potassium ion | $z_K$ | 1 | $dimensionless$ |
| Maximal conductance of inward rectifying potassium channel | $\bar{g}_{Kir}$ | 13.816 | $nS$ |
| Maximal conductance of small conductance potassium channel | $\bar{g}_{Ksk}$ | 2.2515 | $pA * mM^{-1}$ |
| Maximal conductance for sodium-potassium ATPase | $K_{nak}$ | 1085.7 | $pA$ |
| Reaction rates of $I_{NaK}$ | $k_{2,nak}$ | 0.04 | $ms^{-1}$ |
| | $k_{3,nak}$ | 0.01 | $ms^{-1}$ |
| | $k_{4,nak}$ | 0.165 | $ms^{-1}$ |
| Dissociation constants of $I_{NaK}$ | $K_{nak,nae}$ | 69.8 | $mM$ |
| | $K_{nak,nai}$ | 4.05 | $mM$ |
| | $K_{nak,ke}$ | 0.258 | $mM$ |
| | $K_{nak,ki}$ | 32.88 | $mM$ |
| Maximal conductance for calcium ATPase | $k_{pmca}$ | 2.233 | $pA * ms^{-1}$ |
| Reaction rates of $I_{pmca}$ | $k_{2,pc}$ | 0.001 | $ms^{-1}$ |
| | $k_{3,pc}$ | 0.001 | $ms^{-1}$ |
| | $k_{4,pc}$ | 1 | $ms^{-1}$ |
| Dissociation constants of $I_{pmca}$ | $K_{pc,e}$ | 2 | $mM$ |

| | | | |
|---|---|---|---|
| Maximal conductance for sodium-calcium exchanger | $k_{xm}$ | 0.0166 | $pA * ms^{-1}$ |
| Energy barrier parameter of $I_{NaCaX}$ | $\delta_{xm}$ | 0.35 | $dimensionless$ |
| Denominator factor of $I_{NaCaX}$ | $\mathcal{D}_{xm}$ | 0.001 | $dimensionless$ |

**Table-2:** Steady state values of ion-channel dynamics of SNc cell model (Francis et al., 2013).

| **Symbol** | **Value** | **Symbol** | **Value** |
|---|---|---|---|
| $V$ | $-49.42\ mV$ | $h_{Na}$ | 0.1848 |
| $[Ca_i]$ | $1.88\ x\ 10^{-4}\ mM$ | $O_{NaHCN}$ | 0.003 |
| $[Na_i]$ | $4.69\ mM$ | $m_{K,dr}$ | 0.003 |
| $[K_i]$ | $126.06\ mM$ | $y_{nak}$ | 0.6213 |
| $m_{Na}$ | 0.0952 | $y_{pc}$ | 0.483 |

#### Calcium Buffering Mechanisms

**Table-3:** Parameter values of calcium buffering mechanisms of SNc cell model (Francis et al., 2013; Marhl et al., 2000).

| **Constant** | **Symbol** | **Value** | **Units** |
|---|---|---|---|
| Calbindin reaction rates | $k_{1,calb}$ | 10 | $mM^{-1} * ms^{-1}$ |
| | $k_{2,calb}$ | $2\ x\ 10^{-3}$ | $ms^{-1}$ |
| Total cytosolic calbindin concentration | $[Calb_{tot}]$ | 0.005 | $mM$ |
| Calmodulin reaction rates | $k_{cam}^{cb}$ | 12000 | $mM^{-2} * ms^{-1}$ |
| | $k_{cam}^{nb}$ | $3.7\ x\ 10^{6}$ | $mM^{-2} * ms^{-1}$ |
| | $k_{cam}^{cd}$ | $3\ x\ 10^{-3}$ | $ms^{-1}$ |
| | $k_{cam}^{nd}$ | 3 | $ms^{-1}$ |
| Total cytosolic calmodulin concentration | $[Cam_{tot}]$ | 0.0235 | $mM$ |

| | | | |
|---|---|---|---|
| The maximal rate constant of SERCA | $k_{serca,er}$ | 0.02 | $mM^{-1} * ms^{-1}$ |
| Maximal permeability of calcium channels in the ER membrane | $k_{ch,er}$ | 3 | $ms^{-1}$ |
| Half saturation for calcium | $K_{ch,er}$ | 0.005 | $mM$ |
| Maximal rate constant for calcium leak flux through the ER membrane | $k_{leak,er}$ | $5 \, x \, 10^{-5}$ | $ms^{-1}$ |
| Ratio of free calcium to total calcium concentration in ER | $\beta_{er}$ | 0.0025 | $dimensionless$ |
| Volume ratio between the ER and cytosol | $\rho_{er}$ | 0.01 | $dimensionless$ |
| Maximal permeability of MCUs | $k_{mcu,mt}$ | $3 \, x \, 10^{-4}$ | $mM * ms^{-1}$ |
| Half saturation for calcium | $K_{mcu,mt}$ | $8 \, x \, 10^{-4}$ | $mM$ |
| Maximal rate of calcium flux through $[Na^{+}]/[Ca^{2+}]$ exchangers and mPTPs | $k_{out,mt}$ | 0.125 | $ms^{-1}$ |
| Half saturation for calcium | $K_{out,mt}$ | 0.005 | $mM$ |
| Maximal rate constant for calcium leak flux through the MT membrane | $k_{leak,mt}$ | $6.25 \, x \, 10^{-6}$ | $ms^{-1}$ |
| Ratio of free calcium to total calcium concentration in MT | $\beta_{mt}$ | 0.0025 | $dimensionless$ |
| Volume ratio between the MT and cytosol | $\rho_{mt}$ | 0.01 | $dimensionless$ |

**Table-4:** Steady state values of calcium buffering mechanisms of SNc cell model (Francis et al., 2013; Marhl et al., 2000).

| Symbol | Value | Symbol | Value |
|---|---|---|---|
| $[Ca_{er}]$ | $1 \, x \, 10^{-3} \, mM$ | $[Calb]$ | $26 \, x \, 10^{-4} \, mM$ |
| $[Ca_{mt}]$ | $4 \, x \, 10^{-4} \, mM$ | $[Cam]$ | $222 \, x \, 10^{-4} \, mM$ |

#### Energy Metabolism

**Table-5:** Parameter values of energy metabolism of SNc cell model (Cloutier and Wellstead, 2010, 2012).

| Constant | Symbol | Value | Units |
|---|---|---|---|
| Extracellular glucose concentration | $[GLC_e]$ | 1 | $mM$ |
| Hexokinase maximal flux | $\bar{v}_{hk}$ | $2.5\ x\ 10^{-3}$ | $mM * ms^{-1}$ |
| Affinity constant for ATP | $K_{m,ATP,hk}$ | 0.5 | $mM$ |
| Inhibition constant for F6P | $K_{i,F6P}$ | 0.068 | $mM$ |
| Phosphofructokinase maximal flux | $\bar{v}_{pfk}$ | $3.8\ x\ 10^{-3}$ | $mM * ms^{-1}$ |
| Affinity constant for F6P | $K_{m,F6P,pfk}$ | 0.18 | $mM$ |
| Affinity constant for ATP | $K_{m,ATP,pfk}$ | 0.05 | $mM$ |
| Affinity constant for F26P | $K_{m,F26P,pfk}$ | 0.01 | $mM$ |
| Activation constant for AMP | $K_{a,AMP,pfk}$ | 0.05 | $mM$ |
| Inhibition constant for ATP | $K_{i,ATP}$ | 1 | $mM$ |
| Coefficient constant for AMP | $nAMP$ | 0.5 | $dimensionless$ |
| Coefficient constant for ATP | $nATP$ | 0.4 | $dimensionless$ |
| Total energy shuttles concentration | $[ANP]$ | 2.51 | $mM$ |
| Coefficient constant for ADP | $Q_{adk}$ | 0.92 | $dimensionless$ |
| Phosphofructokinase-2 maximal forward flux | $\bar{v}_{pfk2,f}$ | $2\ x\ 10^{-7}$ | $mM * ms^{-1}$ |
| Phosphofructokinase-2 maximal reverse flux | $\bar{v}_{pfk2,r}$ | $1.036\ x\ 10^{-7}$ | $mM * ms^{-1}$ |
| Affinity constant for F6P | $K_{m,F6P,pfk2}$ | 0.01 | $mM$ |
| Affinity constant for ATP | $K_{m,ATP,pfk2}$ | 0.05 | $mM$ |
| Affinity constant for F26P | $K_{m,F26P,pfk2}$ | 0.0001 | $mM$ |
| Activation constant for AMP | $K_{a,AMP,pfk2}$ | 0.005 | $mM$ |
| Pyruvate kinase maximal flux | $\bar{v}_{pk}$ | $5\ x\ 10^{-3}$ | $mM * ms^{-1}$ |
| Affinity constant for GAP | $K_{m,GAP,pk}$ | 0.4 | $mM$ |
| Affinity constant for ADP | $K_{m,ADP,pk}$ | 0.005 | $mM$ |

| | | | |
|---|---|---|---|
| Oxidative phosphorylation maximal flux | $\bar{v}_{op}$ | $1\ x\ 10^{-3}$ | $mM * ms^{-1}$ |
| Maximal electron transport chain efficiency | $\bar{\eta}_{op}$ | 0.995 | $dimensionless$ |
| Maximal fraction of $asyn^{*}$ effect on the oxidative phosphorylation | $\beta_{op,asyn_{mis}}$ | 0.08 | $dimensionless$ |
| Affinity constant for $asyn^{*}$ | $K_{asyn_{mis}}$ | $8.5\ x\ 10^{-3}$ | $mM$ |
| Affinity constant for PYR | $K_{m,PYR,op}$ | 0.5 | $mM$ |
| Affinity constant for ADP | $K_{m,ADP,op}$ | 0.005 | $mM$ |
| Forward reaction constant of LDH | $k_{ldh,f}$ | $12.5\ x\ 10^{-3}$ | $ms^{-1}$ |
| Reverse reaction constant of LDH | $k_{ldh,r}$ | $2.5355\ x\ 10^{-3}$ | $ms^{-1}$ |
| Maximal lactate fermentation efficiency | $\bar{\eta}_{ldh}$ | 1 | $dimensionless$ |
| Maximal fraction of $ROS$ effect on the lactate fermentation | $\beta_{ldh,ROS}$ | 0.25 | $dimensionless$ |
| Affinity constant for $ROS$ | $K_{ldh,ROS}$ | $10\ x\ 10^{-3}$ | $mM$ |
| MCT maximal influx | $\bar{v}_{lac}$ | $3.55\ x\ 10^{-4}$ | $mM * ms^{-1}$ |
| Coefficient constant for MCT influx | $K_{lac,inf}$ | 0.641 | $dimensionless$ |
| Reaction constant for lactate efflux | $K_{lac,eff}$ | $7.1\ x\ 10^{-4}$ | $ms^{-1}$ |
| ATPase maximal flux | $\bar{v}_{ATPase}$ | $9.355\ x\ 10^{-4}$ | $mM * ms^{-1}$ |
| Affinity constant for ATP | $K_{m,ATP}$ | 0.5 | $mM$ |
| PPP maximal flux | $\bar{v}_{ppp}$ | $3.972\ x\ 10^{-4}$ | $mM * ms^{-1}$ |
| Inhibition constant for $\left(\frac{NADPH}{NADP}\right)$ | $K_{i,NADPH}$ | 20 | $dimensionless$ |
| Total NADPH and NADP concentration | $[NADPH_{tot}]$ | 0.25 | $mM$ |
| GR forward reaction constant | $k_{gr,f}$ | $1.8\ x\ 10^{-4}$ | $mM^{-1} * ms^{-1}$ |
| GR reverse reaction constant | $k_{gr,r}$ | $3.472\ x\ 10^{-7}$ | $mM^{-1} * ms^{-1}$ |
| Total GSH and GSSG concentration | $[GSH_{tot}]$ | 2.5 | $mM$ |
| Reaction constant of DOX | $K_{dox,ROS}$ | $7.5\ x\ 10^{-8}$ | $ms^{-1}$ |

| | | | |
|---|---|---|---|
| CK forward reaction constant | $k_{ck,f}$ | $3\ x\ 10^{-3}$ | $mM^{-1} * ms^{-1}$ |
| CK reverse reaction constant | $k_{ck,r}$ | $1.26\ x\ 10^{-3}$ | $mM^{-1} * ms^{-1}$ |
| Total PCr and Cr concentration | $[PCr_{tot}]$ | 20 | $mM$ |

**Table-6:** Steady state values of energy metabolism of SNc cell model (Cloutier and Wellstead, 2010, 2012).

| Symbol | Value | Symbol | Value |
|---|---|---|---|
| $[F6P]$ | $0.176\ mM$ | $[LAC]$ | $0.598\ mM$ |
| $[F26P]$ | $2.2\ x\ 10^{-3}\ mM$ | $[PCr]$ | $18.04\ mM$ |
| $[GAP]$ | $8.25\ x\ 10^{-2}\ mM$ | $[NADPH]$ | $0.25\ mM$ |
| $[PYR]$ | $0.124\ mM$ | $[GSH]$ | $2.5\ mM$ |
| $[ATP_i]$ | $2.4\ mM$ | | |

#### Dopamine Turnover Processes

**Table-7:** Parameter values for DA turnover processes of SNc cell model (Reed et al., 2012; Tello-Bravo, 2012).

| Constant | Symbol | Value | Units |
|---|---|---|---|
| Average release flux per vesicle | $\psi$ | 17.4391793 | $mM * ms^{-1}$ |
| Initial vesicular DA concentration | $DA_{v_o}$ | 500 | $mM$ |
| Sensitivity to vesicular DA concentration | $DA_{v_s}$ | 0.01 | $mM$ |
| Affinity constant of DA binding to receptors | $DA_{R_a}$ | $5\ x\ 10^{-5}$ | $mM$ |
| Binding sensitivity | $DA_{R_s}$ | 0.01 | $mM$ |
| Activation constant for ATP | $K_{a,RRP}$ | 1.4286 | $mM$ |
| Vesicle recycling maximal flux | $\bar{v}_{nrrp}$ | $1\ x\ 10^{-3}$ | $mM * ms^{-1}$ |
| Maximal vesicle recycling efficiency | $\bar{\eta}_{nrrp}$ | 0.995 | $dimensionless$ |
| Maximal fraction of $asyn^*$ effect on the vesicle | $\beta_{nrrp,asyn_{mis}}$ | 0.08 | $dimensionless$ |

| | | | |
|---|---|---|---|
| Affinity constant for $asyn^*$ | $K_{asyn_{mis}}$ | $8.5\ x\ 10^{-3}$ | $mM$ |
| Reaction constant of $DA_e$ clearance | $k_{comt}$ | 0.0083511 | $ms^{-1}$ |
| Tyrosine concentration | $[TYR]$ | $126\ x\ 10^{-3}$ | $mM$ |
| Affinity constant for $TYR$ | $K_{TYR}$ | $46\ x\ 10^{-3}$ | $mM$ |
| Inhibition constant for $DA_c$ | $K_{i,cda}$ | $11\ x\ 10^{-2}$ | $mM$ |
| Inhibition constant for $DA_e$ | $K_{i,eda}$ | $46\ x\ 10^{-3}$ | $mM$ |
| Maximal velocity of DA synthesis | $\bar{V}_{synt}$ | $25\ x\ 10^{-6}$ | $mM * ms^{-1}$ |
| Affinity constant for $Ca_i$ | $K_{synt}$ | $35\ x\ 10^{-4}$ | $mM$ |
| Maximal velocity of VMAT | $\bar{V}_{cda}$ | $4.67\ x\ 10^{-6}$ | $ms^{-1}$ |
| Affinity constant for $DA_c$ | $K_{cda}$ | $238\ x\ 10^{-4}$ | $mM$ |
| Scaling factor for VMAT | $\alpha_{vmat}$ | $1\ x\ 10^{-3}$ | $dimensionless$ |
| Scaling factor for $ATP_i$ | $\beta_{vmat}$ | 3 | $dimensionless$ |
| Reaction constant of $DA_c$ clearance | $k_{mao}$ | 0.00016 | $ms^{-1}$ |
| Maximal velocity of AADC | $\bar{V}_{aadc}$ | $9.73\ x\ 10^{-5}$ | $mM * ms^{-1}$ |
| Affinity constant for $LDOPA$ | $K_{aadc}$ | 0.13 | $mM$ |
| Maximal velocity of AAT | $\bar{V}_{aat}$ | $5.11\ x\ 10^{-7}$ | $mM * ms^{-1}$ |
| Affinity constant for $LDOPA_e$ | $K_{ldopa_e}$ | $3.2\ x\ 10^{-4}$ | $mM$ |
| Affinity constant for $TYR_e$ | $K_{tyr_e}$ | $6.4\ x\ 10^{-4}$ | $mM$ |
| Affinity constant for $TRP_e$ | $K_{trp_e}$ | $1.5\ x\ 10^{-4}$ | $mM$ |
| Serum concentration of TYR | $[TYR_e]$ | $6.3\ x\ 10^{-4}$ | $mM$ |
| Serum concentration of TRP | $[TRP_e]$ | $8.2\ x\ 10^{-4}$ | $mM$ |
| Serum concentration of LDOPA | $[sLD]$ | $3.6\ x\ 10^{-3}$ | $mM$ |

**Table-8:** Steady state values of DA turnover processes of SNc cell model (Reed et al., 2012; Tello-Bravo, 2012).

| Symbol | Value | Symbol | Value |
|---|---|---|---|
| $[DA_e]$ | $4\ x\ 10^{-6}\ mM$ | $[DA_v]$ | $500\ mM$ |
| $[DA_c]$ | $1\ x\ 10^{-4}\ mM$ | $[LDOPA]$ | $3.6\ x\ 10^{-4}\ mM$ |

#### Molecular Pathways Involved in PD Pathology

**Table-9:** Parameter values of PD pathology pathways of SNc cell model (Cloutier and Wellstead, 2012).

| Constant | Symbol | Value | Units |
|---|---|---|---|
| Activation constant for ATP | $K_{a,leak}$ | 0.5282 | $mM$ |
| Reaction constant for ROS production due to excess dopamine | $k_{dopa}$ | $4.167\ x\ 10^{-4}$ | $mM^{-1} * ms^{-1}$ |
| Affinity constant for $[DA_c]$ | $K_{dopa}$ | 8.5 | $mM$ |
| Reaction constant for catalase | $k_{cat}$ | $2.35\ x\ 10^{-5}$ | $ms^{-1}$ |
| Reaction constant for alpha-synuclein oxidation | $k_{syn}$ | $1.39\ x\ 10^{-8}$ | $mM * ms^{-1}$ |
| Reaction constant for alpha-synuclein consumption | $k_{to}$ | $1.39\ x\ 10^{-7}$ | $ms^{-1}$ |
| Reaction constant for alpha-synuclein aggregation | $k_{agg}$ | $2.08\ x\ 10^{-10}$ | $ms^{-1}$ |
| Affinity constant for $ASYN_{mis}$ | $K_{agg}$ | $7.5\ x\ 10^{-3}$ | $mM$ |
| Reaction constant for tagging of damaged protein | $k_{tag}$ | $7.64\ x\ 10^{-11}$ | $mM^{-1} * ms^{-1}$ |
| Total ubiquitin concentration | $[Ub_{tot}]$ | $10.5\ x\ 10^{-3}$ | $mM$ |
| Reaction constant for damaged protein disposal by the proteasome | $k_{prt}$ | $2.08\ x\ 10^{-10}$ | $ms^{-1}$ |
| Affinity constant for $ASYN_{agg}$ | $K_{prt}$ | $5\ x\ 10^{-3}$ | $mM$ |
| Fraction reduction of proteasome activity by $ASYN_{agg}$ | $\beta_{prt}$ | 0.25 | $dimensionless$ |
| Reaction constant for $ASYN_{agg}$ disposal by lysosome | $k_{lyso}$ | $2.08\ x\ 10^{-11}$ | $ms^{-1}$ |
| Reaction constant for Lewy bodies from $ASYN_{agg}$ | $k_{lb}$ | $2.08\ x\ 10^{-11}$ | $ms^{-1}$ |
| Affinity constant for $ASYN_{agg}$ | $K_{lb}$ | $5\ x\ 10^{-3}$ | $mM$ |

**Table-10:** Steady state values of PD pathology pathways of SNc cell model (Cloutier and Wellstead, 2012).

| Symbol | Value | Symbol | Value |
|---|---|---|---|
| $[ROS]$ | $1\ x\ 10^{-3}\ mM$ | $[ASYN_{tag}]$ | $1\ x\ 10^{-5}\ mM$ |
| $[ASYN]$ | $0.1\ mM$ | $[ASYN_{agg}]$ | $0\ mM$ |
| $[ASYN_{mis}]$ | $1\ x\ 10^{-3}\ mM$ | $[LB]$ | $0\ mM$ |

#### Apoptotic Pathways

**Table-11:** Parameter values of apoptotic pathways of SNc cell model (Hong et al., 2012).

| Constant | Symbol | Value | Units |
|---|---|---|---|
| Forward reaction constant for $[Ca_i.Calpain]$ | $k_1^+$ | 1 | $mM^{-1} * ms^{-1}$ |
| Reverse reaction constant for $[Ca_i.Calpain]$ | $k_1^-$ | $1\ x\ 10^{-3}$ | $ms^{-1}$ |
| Forward reaction constant for $[Calpain^*]$ | $k_2^+$ | $1\ x\ 10^{-3}$ | $ms^{-1}$ |
| Forward reaction constant for $[Calpain^*.Casp12]$ | $k_3^+$ | 1 | $mM^{-1} * ms^{-1}$ |
| Reverse reaction constant for $[Calpain^*.Casp12]$ | $k_3^-$ | $1\ x\ 10^{-3}$ | $ms^{-1}$ |
| Forward reaction constant for $[Casp12^*]$ | $k_4^+$ | $1\ x\ 10^{-3}$ | $ms^{-1}$ |
| Forward reaction constant for $[Casp12^*.Casp9]$ | $k_5^+$ | 10 | $mM^{-1} * ms^{-1}$ |
| Reverse reaction constant for $[Casp12^*.Casp9]$ | $k_5^-$ | $5\ x\ 10^{-4}$ | $ms^{-1}$ |
| Forward reaction constant for $[Casp9^*]$ | $k_6^+$ | $1\ x\ 10^{-3}$ | $ms^{-1}$ |
| Forward reaction constant for $[Casp9^*.Casp3]$ | $k_7^+$ | 10 | $mM^{-1} * ms^{-1}$ |
| Reverse reaction constant for $[Casp9^*.Casp3]$ | $k_7^-$ | $5\ x\ 10^{-4}$ | $ms^{-1}$ |
| Forward reaction constant for $[Casp3^*]$ | $k_8^+$ | $1\ x\ 10^{-4}$ | $ms^{-1}$ |
| Forward reaction constant for $[Apop]$ | $k_9^+$ | 1 | $mM^{-1} * ms^{-1}$ |
| Forward reaction constant for $[Casp9^*]$ | $k_{10}^+$ | $1\ x\ 10^{-3}$ | $ms^{-1}$ |
| Forward reaction constant for $[Casp9^*.IAP]$ | $k_{11}^+$ | 5 | $mM^{-1} * ms^{-1}$ |
| Reverse reaction constant for $[Casp9^*.IAP]$ | $k_{11}^-$ | $35\ x\ 10^{-7}$ | $ms^{-1}$ |
| Forward reaction constant for $[Casp3^*.IAP]$ | $k_{12}^+$ | 5 | $mM^{-1} * ms^{-1}$ |
| Reverse reaction constant for $[Casp3^*.IAP]$ | $k_{12}^-$ | $35\ x\ 10^{-7}$ | $ms^{-1}$ |
| Forward reaction constant for $[ROS_{mit}]$ | $k_{13}^+$ | 0.5 | $mM^{-1} * ms^{-1}$ |
| Forward reaction constant for $[PTP_{mit}^*]$ | $k_{14}^+$ | 0.5 | $mM^{-1} * ms^{-1}$ |
| Forward reaction constant for $[Cytc]$ | $k_{15}^+$ | 1 | $mM^{-1} * ms^{-1}$ |
| Forward reaction constant for $[Cytc.Casp9]$ | $k_{16}^+$ | 1 | $mM^{-1} * ms^{-1}$ |
| Reverse reaction constant for $[Cytc.Casp9]$ | $k_{16}^-$ | $1\ x\ 10^{-3}$ | $ms^{-1}$ |

**Table-12:** Steady state values of energy metabolism of SNc cell model (Hong et al., 2012).

| Symbol | Value | Symbol | Value |
|---|---|---|---|
| $[Calpain]$ | 1 | $[ROS_{mit}]$ | 0 |
| $[Ca_i.Calpain]$ | 0 | $[PTP^*_{mit}]$ | 1 |
| $[Calpain^*]$ | 0 | $[Cytc_{mit}]$ | 1 |
| $[Casp12]$ | 1 | $[Cytc]$ | 0 |
| $[Calpain^*.Casp12]$ | 0 | $[Cytc.Casp9]$ | 0 |
| $[Casp12^*]$ | 0 | $[Casp9]$ | 1 |
| $[Casp12^*.Casp9]$ | 0 | $[Casp9^*]$ | 0 |
| $[Casp3]$ | 1 | $[Casp9^*.Casp3]$ | 0 |
| $[Casp3^*]$ | 0 | $[IAP]$ | 1 |
| $[Casp9^*.IAP]$ | 0 | $[Casp3^*.IAP]$ | 0 |
| $[Apop]$ | 0 | | |

#### Energy Consumption

**Table-13:** Parameters for energy consumption processes of SNc cell model.

| Constant | Symbol | Value | Units |
|---|---|---|---|
| Faraday's constant | $F$ | 96485 | $coulomb * mole^{-1}$ |
| Cytosolic volume | $v_{cyt}$ | $\phi_{cyt} * v_{pmu}$ | $pl$ |
| Pacemaking unit (PMU) volume | $v_{pmu}$ | 5 | $pl$ |
| Fraction of cytosolic volume | $\phi_{cyt}$ | 0.5 | $dimensionless$ |
| Scaling factor for synaptic recycling | $\lambda_{sr}$ | 100 | $dimensionless$ |
| Scaling factor for neurotransmitter packing | $\lambda_{np}$ | 1 | $dimensionless$ |
| Ratio of free calcium to total calcium concentration in ER | $\beta_{er}$ | 0.0025 | $dimensionless$ |
| Volume ratio between the ER and cytosol | $\rho_{er}$ | 0.01 | $dimensionless$ |
| Scaling factor for proteasome | $\lambda_{prt}$ | 25 | $dimensionless$ |

| | | | |
|---|---|---|---|
| Scaling factor for ubiquitination | $\lambda_{tag}$ | 3 | $dimensionless$ |
| Scaling factor for lysosome | $\lambda_{lyso}$ | 10 | $dimensionless$ |
