## Supplementary material-4 for "Influence of Energy Deficiency on the Molecular Processes of *Substantia Nigra Pars Compacta* Cell for Understanding Parkinsonian Neurodegeneration: A Comprehensive Biophysical Computational Model"

### RECEPTOR MODELING

#### AMPA/Kainate Receptors

The simplest model that approximates the kinetics of the fast AMPA/kainate type of glutamate receptors can be represented by the two-state diagram:

$$C + T \overset{(\alpha/\beta)}{\Longleftrightarrow} O \tag{1}$$

where, $\alpha$ and $\beta$ are voltage-independent forward and backward rate constants, $C$ is the closed state of the receptor, $O$ is the open state of the receptor, and $T$ is the neurotransmitter. If $r$ is defined as the fraction of the receptors in the open state, it is then described by the following first-order kinetic equation:

$$\frac{d(r)}{dt} = \alpha * [T] * (1 - r) - \beta * r \tag{2}$$

and the postsynaptic current $(I_{AMPA})$ is given by,

$$I_{AMPA} = \bar{g}_{AMPA} * r * (V - E_{AMPA}) \tag{3}$$

where, $\bar{g}_{AMPA}$ is the maximal conductance, $E_{AMPA}$ is the reversal potential, $V$ is the postsynaptic membrane potential, $[T]$ is the neurotransmitter, and $r$ is the fraction of the receptors in the open state.

#### NMDA Receptors

The slower NMDA type of glutamate receptors can be represented with a two-state model similar to AMPA/kainate receptors, with a voltage-dependent term representing magnesium block. Using the scheme in Eqs. 1 and 2, the postsynaptic current is given by

$$I_{NMDA} = \bar{g}_{NMDA} * r * B(V) * (V - E_{NMDA}) \tag{4}$$

where, $\bar{g}_{NMDA}$ is the maximal conductance, $E_{NMDA}$ is the reversal potential, $B(V)$ is the magnesium block, $V$ is the postsynaptic membrane potential, and $r$ is the fraction of the receptors in the open state.

$$B(V) = \frac{1}{1 + \left(\frac{[Mg^{2+}]}{3.57} * e^{-0.062 * V}\right)} \tag{5}$$

where, $[Mg^{2+}]$ is the external magnesium concentration, and $V$ is the postsynaptic membrane potential.

**GABA$_A$ Receptors**

GABA$_A$ receptors can also be represented by the scheme in Eqs. 1 and 2, with the postsynaptic current given by

$$I_{GABA_A} = \bar{g}_{GABA_A} * r * \left(V - E_{GABA_A}\right) \tag{6}$$

where, $\bar{g}_{GABA_A}$ is the maximal conductance, $E_{GABA_A}$ is the reversal potential, $V$ is the postsynaptic membrane potential, and $r$ is the fraction of the receptors in the open state.

**GABA$_B$ Receptors**

The stimulus dependency of GABA$_B$ responses, unfortunately, cannot be handled correctly by a two-state model. The simplest model of GABA$_B$-mediated currents has two variables:

$$\frac{d(r)}{dt} = K_1 * [T] * (1 - r) - K_2 * r \tag{7}$$

$$\frac{d(s)}{dt} = K_3 * r - K_4 * s \tag{8}$$

and the postsynaptic current $\left(I_{GABA_B}\right)$ is given by,

$$I_{GABA_B} = \bar{g}_{GABA_B} * \frac{s^n}{s^n + K_d} * \left(V - E_{GABA_B}\right) \tag{9}$$

where, $\bar{g}_{GABA_B}$ is the maximal conductance, $E_{GABA_B}$ $(= V_K)$ is the reversal potential, $V$ is the postsynaptic membrane potential, $r$ is the fraction of the receptors in the open state, $s$ is the fraction of activated G-proteins, $K_d$ is the dissociation constant of the binding of $s$ on the $K^+$ channels, $K_1$ and $K_2$ are voltage-independent forward and backward rate constants for $r$, $K_3$

and $K_4$ are voltage-independent forward and backward rate constants for $s$, and $[T]$ is the neurotransmitter.

**Overall Synaptic Current**

The overall synaptic input current flux $(J_{syn})$ to SNc neuron is given by,

$$J_{syn} = -\frac{1}{F * v_{cyt}} * \left(I_{AMPA} + I_{NMDA} + I_{GABA_A} + I_{GABA_B}\right) \tag{10}$$

where, $I_{AMPA}$ is the excitatory AMPA synaptic current, $I_{NMDA}$ is the excitatory NMDA synaptic current, $I_{GABA_A}$ is the inhibitory GABA$_A$ synaptic current, $I_{GABA_B}$ is the inhibitory GABA$_B$ synaptic current, $F$ is the Faraday's constant, and $v_{cyt}$ is the cytosolic volume.

**Table-1:** Parameter values of receptor models

| Constant | Symbol | Value | Units |
|---|---|---|---|
| Faraday's constant | $F$ | 96485 | $coulomb * mole^{-1}$ |
| Cytosolic volume | $v_{cyt}$ | $\phi_{cyt} * v_{pmu}$ | $pl$ |
| Fraction of cytosolic volume | $\phi_{cyt}$ | 0.5 | $dimensionless$ |
| Pacemaking unit (PMU) volume | $v_{pmu}$ | 5 | $pl$ |
| Maximal conductance of AMPA receptor | $\bar{g}_{AMPA}$ | $0.35 - 1$ | $nS$ |
| Maximal conductance of NMDA receptor | $\bar{g}_{NMDA}$ | $0.01 - 0.6$ | $nS$ |
| Concentration of Magnesium | $[Mg^{2+}]$ | $1 - 2$ | $mM$ |
| Maximal conductance of GABA$_A$ receptor | $\bar{g}_{GABA_A}$ | $0.25 - 1.2$ | $nS$ |
| Maximal conductance of GABA$_B$ receptor | $\bar{g}_{GABA_B}$ | 0.06 | $nS$ |
| Dissociation constant of the binding of $s$ on the $K^+$ channels | $K_d$ | 100 | $\mu M^4$ |
| Voltage-independent forward rate constant for $r$ of GABA$_B$ | $K_1$ | $9 x 10^4$ | $M^{-1} * sec^{-1}$ |

| | | | |
|---|---|---|---|
| Voltage-independent backward rate constant for $r$ of $GABA_B$ | $K_2$ | 1.2 | $sec^{-1}$ |
| Voltage-independent forward rate constant for $s$ of $GABA_B$ | $K_3$ | 180 | $sec^{-1}$ |
| Voltage-independent backward rate constant for $s$ of $GABA_B$ | $K_4$ | 34 | $sec^{-1}$ |
| Cooperativity constant (binding sites) | $n$ | 4 | $dimensionless$ |
| Reversal potential of AMPA | $E_{AMPA}$ | 0 | $mV$ |
| Reversal potential of NMDA | $E_{NMDA}$ | 0 | $mV$ |
| Reversal potential of $GABA_A$ | $E_{GABA_A}$ | $-80$ | $mV$ |
| Reversal potential of $GABA_B$ | $E_{GABA_B}$ | $-95$ | $mV$ |
| Voltage-independent forward rate constant for $r$ $(\alpha)$ | AMPA | $1.1\ x\ 10^6$ | $M^{-1} * sec^{-1}$ |
| | NMDA | $7.2\ x\ 10^4$ | $M^{-1} * sec^{-1}$ |
| | $GABA_A$ | $5\ x\ 10^6$ | $M^{-1} * sec^{-1}$ |
| Voltage-independent backward rate constant for $r$ $(\beta)$ | AMPA | 190 | $sec^{-1}$ |
| | NMDA | 6.6 | $sec^{-1}$ |
| | $GABA_A$ | 180 | $sec^{-1}$ |
